# Structural dynamics and functional cooperativity of human NQO1 by ambient temperature serial crystallography and simulations

**DOI:** 10.1101/2023.12.21.572834

**Authors:** Alice Grieco, Sergio Boneta, José A. Gavira, Angel L. Pey, Shibom Basu, Julien Orlans, Daniele de Sanctis, Milagros Medina, Jose Manuel Martin-Garcia

## Abstract

The human NQO1 (hNQO1) is a FAD-dependent oxidoreductase that catalyzes the two-electron reduction of quinones to hydroquinones, being essential for the antioxidant defense system, stabilization of tumor suppressors, and activation of quinone-based chemotherapeutics, and it is over-expressed in several tumors, which makes it an attractive cancer drug target. To decipher new structural insights into the flavin reductive half-reaction of the catalytic mechanism of hNQO1, we have carried serial crystallography experiments at new ID29 beamline of the ESRF to determine, to the best of our knowledge, the first structure of the hNQO1 in complex with NADH. The use of room temperature serial crystallography with microcrystals has been key to study this mechanism. We have also performed molecular dynamics simulations of free hNQO1 and in complex with NADH. Both structural results and MD simulations have supported that the binding of NADH significantly decreases protein dynamics and stabilizes hNQO1 especially at the dimer core and interface. This is the first structural evidence that the hNQO1 functional cooperativity is driven by structural communication between the active sites through long-range propagation of cooperative effects across the hNQO1 structure. Altogether, these results pave the way for future time-resolved studies, both at XFELs and synchrotrons, of the dynamics of hNQO1 upon binding to NADH as well as during the FAD cofactor reductive half-reaction. This knowledge will allow us to reveal unprecedented structural information of the relevance of the dynamics during the catalytic function of hNQO1.

## 1 INTRODUCTION

The human NAD(P)H quinone oxidoreductase 1 (hNQO1) is a flavo-oxidoreductase essential for the antioxidant defense system, stabilization of tumor suppressors, and activation of quinone-based chemotherapeutics (Ross and Siegel 2017; Pey et al. 2016). Alterations in hNQO1 function are associated with cancer and with Alzheimer’s and Parkinson’s disease, making this enzyme an attractive target for drug discovery (Beaver et al. 2019). hNQO1 is a homodimer of 62 kDa with two identical 31 kDa interlocking protomers and two active sites. These active sites are located at the interface between the two protomers consisting of two domains that involve residues from both polypeptide chains and the corresponding two noncovalently bound flavin adenine nucleotide (FAD) cofactors, one at each catalytic site (Li et al. 1995; Asher et al. 2006; Faig et al. 2000; Skelly et al. 1999). hNQO1 primarily catalyzes the two-electron reduction of quinones to hydro-quinones by a two-step mechanism commonly known as “ping-pong” (Anoz-Carbonell et al. 2020), in which the electron donor molecule NAD(P)H enters the active site and donates a hydride to the FAD cofactor reducing it to the hydroquinone state (namely FADH^-^). Then the oxidized NAD^+^ leaves the active site, and it is eventually replaced by a quinone substrate that is subsequently reduced to tis hydroquinone form by FADH^-^.

Crystal structures of hNQO1 with and without the inhibitors dicoumarol and cibacron blue, chemotherapeutic drugs, and the substrate duroquinone, have shown that hNQO1 undergoes local conformational changes in which the active site opens and closes during the catalysis (Faig et al. 2000; Winski et al. 2001; Asher et al. 2006; W. D. Lienhart et al. 2014). However, the difficulty of obtaining structural and dynamic information on many of the intermediate states by standard macromolecular crystallography at synchrotrons, has limited the understanding of the redox mechanism of hNQO1. Other biophysical techniques including isothermal titration calorimetry, hydrogen/deuterium exchange, absorption spectroscopy, and stopped flow have supported the hypothesis that the two active/binding sites of hNQO1 act cooperatively and display highly collective inter-domain and inter-monomer communication and dynamics (Pey, Megarity, and Timson 2014; Clavería-Gimeno, Velazquez-Campoy, and Pey 2017; Pey 2018; Pacheco-Garcia et al. 2021). This coupled network of amino acids is sensitive to ligand binding as it decreases protein dynamics significantly and stabilizes hNQO1 especially at the dimer core and interface. Such observation provides experimental evidence for the two active sites in the homodimer catalyzing the hydride exchange with different rates being driven by structural communication and the breaking and formation of hydrogen bonds and salt bridges across the hNQO1 protein structure (Pacheco-Garcia et al. 2021; Anoz-Carbonell et al. 2020; Pacheco-garcia et al. 2022). For this reason, understanding the hNQO1 redox mechanism and the conformational dynamics upon interaction with substrates is critical to unravel its roles as an antioxidant and potential target to treat cancer, neurological disorders and cardiovascular diseases (Beaver et al. 2019; Betancor-Fernández et al. 2018). However, a detailed structural explanation for these observations, and the derived mechanistic implications, is difficult to provide by standard crystallography as crystal freezing might select one conformation above others (Fraser et al. 2011).

Despite X-ray crystallography being the main way of uncovering the 3D structures of biomacromolecules, traditionally, it has been viewed as a static technique, with limited applicability to study protein dynamics, which fundamentally relies on multiple conformational states and the transitions between them. However, owing to the convergence of several recent experimental and technical developments, X-ray crystallography is increasingly well-positioned to provide insights into the connection between protein flexibility and function. In this regard, the development of new light sources, such as X-ray free electron lasers (XFELs) and sample handling procedures, has brought about a resurgence of room temperature crystallography through so-called serial femtosecond crystallography (SFX). Also, the high intensity of state-of-the-art synchrotron sources, combined with ultrafast detectors, has enabled similar experiments in the form of serial synchrotron crystallography (SSX). To date, the number of room temperature SSX experiments (more than 60 experiments) has grown exponentially since the first experiment was conducted in 2014 (Gati et al. 2014). A further development is represented by the advent of Diffraction Limited Storage rings (Raimondi et al. 2023), capable of providing a much more brilliant beam at the sample position and allowing to perform diffraction experiments at short time scales. The new ID29 is the first beamline completely dedicated to room temperature serial crystallography at a fourth-generation storage ring. Recently, we have shown the benefit of room temperature data collection using serial femtosecond crystallography (SFX) by determining the SFX structure of hNQO1 (Doppler et al. 2023). Our results revealed that several residues in the catalytic site, described to play a key role in the function of the protein, show an unexpected flexibility within the crystals (Doppler et al. 2023). This high plasticity of hNQO1 in the catalytic site provided us with the first structural evidence that the hNQO1 functional negative cooperativity is driven by structural communication between the active sites through long-range propagation of cooperative effects across the protein structure.

In the work presented here, we have conducted one of the first SSX experiments at the new ID29 beamline at the ESRF Extremely Brilliant Source (ESRF-EBS) to determine, to the best of our knowledge, the first structure of the hNQO1 in complex with NADH. Until now, the only structure of an NQO1 protein in complex with NADP^+^ was reported by Amzel and *co-workers* from rat liver (Li et al. 1995). However, no structure was ever deposited in the PDB. With the goal of obtaining the first structure of the hNQO1 in complex with NADH, we have conducted SSX experiments at ID29 beamline of the ESRF-EBS, as well as performed molecular dynamics (MD) simulations. The results presented here will pave the way for future TR-SX studies both at XFELs and synchrotrons to study the dynamics of hNQO1 upon binding of NADH. This knowledge will allow us to reveal unprecedented structural information of the kinetic mechanism that determines the catalytic function of hNQO1. Furthermore, gaining insights into these functional aspects of hNQO1 at the molecular level will be valuable to advance in the design of new, more potent, and effective inhibitors that can be used in clinical realm.

## 2 MATERIALS AND METHODS

### 2.1 Expression and purification of human NQO1 protein

Protein expression and purification of hNQO1 (EC 1.6.99.2) were performed as previously described (Doppler et al. 2023). Briefly, *Escherichia coli BL21* (DE3) cells containing the DNA sequence of hNQO1 were grown overnight in LB culture medium supplemented with 0.1 mg/mL ampicillin (LBA) at 37°C. This culture was diluted in 4 L of fresh LBA and grown at 37 °C until the optical density at 600 nm reached a value between 0.6 and 0.8. Expression was then induced with IPTG (isopropyl β-D-1-thiogalactopyranoside) at a final concentration of 0.5 mM. Induced cells were further incubated for 4 h at 28 °C, harvested by centrifugation and resuspended in 40 mL of binding buffer (BB, 20 mM sodium phosphate, 300 mM NaCl and 50 mM imidazole at pH 7.4) containing one EDTA-free protease Inhibitor tablet. Cells were lysed by sonication on ice. Lysate was cleared by centrifugation at 30,000 rpm at 4 °C for 40 min. The supernatant containing hNQO1 was filtered through 0.45 μm filters and subsequently loaded onto immobilized Ni^2+^ affinity chromatography column previously equilibrated with BB. After collecting the flowthrough, the column was washed with 20 column volumes (CVs) of BB and eluted with 10 CVs of elution buffer (BB containing 500 mM imidazole). The elute protein was dialyzed against 50 mM K-HEPES at pH 7.4. NQO1 was further purified by size exclusion chromatography using a HiLoad 16/600 Superdex 200 prep grade (GE Healthcare) column using the elution buffer 20 mM K-HEPES pH 7.4, 200 mM NaCl, 1 mM FAD. Pure protein was concentrated to a final concentration of 25 mg/ml, flash frozen and stored at −80 °C.

### 2.2 Protein crystallization

Microcrystals of hNQO1 were obtained on-site at the ID29 Serial Macromolecular Crystallography (SMX) beamline at the ESRF-EBS by the batch with agitation method as follows: in a 3 mL glass vial, 150 μL of the protein solution at 25 mg/mL were added dropwise to 450 μL of the precipitant solution (0.1 M Tris pH 8.5, 0.2 M sodium acetate, 20 % polyethylene glycol (PEG) 3350) while stirring at 200 rpm. Upon addition of the protein, the solution turned turbid immediately and needle-shaped crystals of about 10-30 μm in their longest dimension (Figure S1) grew at room temperature in about 1 h. Microcrystals of hNQO1 with the ligand NADH were obtained by the soaking method as follows: a NADH solution of 100 mM was prepared in the protein crystallization buffer, then 5 μL of this solution were taken out, mixed with 50 μL of the original crystal slurry and incubated for 1 h prior to data collection. Microcrystals, initially yellow, turned clear immediately upon mixing, indicating the reduction of the enzyme by NADH.

### 2.3 Sample delivery using a small SOS-chip

Microcrystals of hNQO1 and hNQO1 with NADH at the concentration of 10^8^ crystals/mL were loaded by pipetting 3.5 μL of a crystal slurry and sandwiched between two 13 μm mylar films on a small version of the sheet-on-sheet sandwich chip (SOS-chip) (Doak et al. 2018) with an opening of 5 x 10 mm. The mylar-mylar sandwich containing the crystal slurry was held in place and sealed by the sample holder. The mylar-sample-mylar sandwich was placed between the two half frames of the chip holder leading to further spreading and thinning of the film by capillary action. The two frame halves were then sealed together with one clamping screw. Figure 1A illustrates how sample loading in the SOS-chip was carried out.

**Figure 1.**
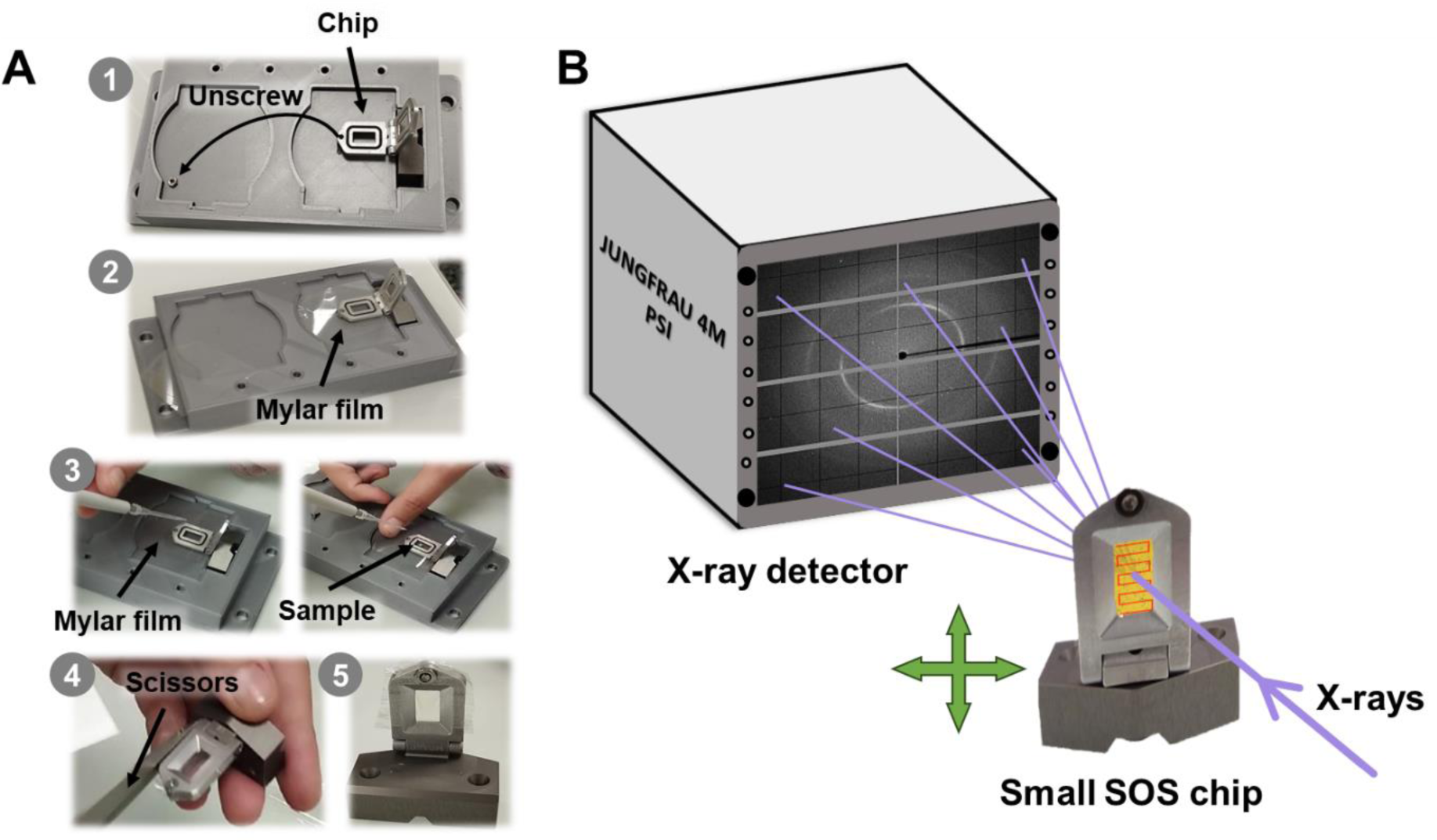
Sample loading and data collection setup at the ID29 beamline. **A)** Steps 1-5 for loading the small SOS chip: 1) unscrew and place the chip in the chip holder; 2) place a mylar film over the chip; 3) apply the sample on the mylar film, place another mylar film on top and screw again; 4) remove the excess mylar film; and 5) chip loaded and mounted. **B)** Schematic diagram of the setup. A 1% bandwidth X-ray beam with a pulse length of 90 µs at repetition rate of 231.25 Hz was used. The X-rays hit the sample that is immobilized in the small SOS chip, which moves from left to right in a zig-zag pattern across the X-Y axes. Diffraction is collected on a JUNGFRAU 4M detector.

### 2.4 Serial synchrotron data collection at the new ID29 beamline

The serial synchrotron crystallography data were collected at the new ID29 SMX beamline (*Orlans et al., Manuscript in preparation*) of the ESRF-EBS using a wavelength of 1.072 Å with a pulse length of 90 µs at repetition rate of 231.25 Hz. The X-ray beam flux at the sample was about 8 x 10^14^ ph/s and focused to 4 x 2 (H x V) μm (FWHM). Data were recorded with a newly developed MD3upSSX diffractometer (Arinax and EMBL-GR) and a JUNGFRAU 4M detector (Mozzanica et al. 2018) with integration time of 95 μs to ensure full recording of the X-ray pulse and automatically corrected for pedestal and geometrical reconstruction with a LImA2 data acquisition library developed at the ESRF (https://limagroup.gitlab-pages.esrf.fr/lima2/). The sample-to-detector distance was set to 150 mm. The final data collection statistics for both proteins, free hNQO1 and hNQO1 in complex with NADH (hNQO1-NAD^+^/H) are given in Table 1. The experimental set-up is shown in Figure 1B.

**Table 1.** Serial X-ray data collection and refinement statistics. Values for the outer shell are given in parentheses.

|  | Free hNQO1 | hNQO1-NAD<sup>+</sup>/H |
| --- | --- | --- |
| *Data collection statistics* |  |  |
| X-ray source / beamline | ID29 | ID29 |
| No. chips used | 5 | 6 |
| Data collection time (min) | 40 | 48 |
| Wavelength (Å) | 1,072 Å | 1,072 Å |
| Temperature (K) | 295 K | 295 K |
| Detector | JUNGFRAU 4M | JUNGFRAU 4M |
| Space group | P2<sub>1</sub>2<sub>1</sub>2<sub>1</sub> | P2<sub>1</sub>2<sub>1</sub>2<sub>1</sub> |
| a, b, c (Å) | 62.0, 108.0, 200.0 | 61.1, 106.8, 196.0 |
| α, β, γ (°) | α = β = γ = 90 | α = β = γ = 90 |
| Resolution range (Å) | 100.0-2.5 Å (20.56-20.50 Å) | 98.0-2.7 Å (20.77-20.70 Å) |
| Completeness (%) | 97.1% (96.0%) | 100.0% (100.0%) |
| CC* (%) | 0.989 (0.532) | 0.977 (0.523) |
| Multiplicity | 154.1 (42.3) | 95.4 (33.9) |
| R<sub>split</sub> | 14.2 (150.3) | 15.4 (97.6) |
| Avg. I/σ (I) | 9.9 (1.8) | 7.6 (0.8) |
| Overall B factor from Wilson plot (Å<sup>2</sup>) | 73.7 | 75.6 |
| *Refinement Statistics* |  |  |
| Resolution range (Å) | 100.0-2.5 Å (20.56-20.50 Å) | 98.0-2.7 Å (20.77-20.70 Å) |
| No. of reflections, working set | 41,455 | 32,476 |
| No. of reflections, test set | 4,581 | 3,618 |
| R<sub>work</sub>/R<sub>free</sub> (%) | 19.87 % / 23.87 % | 19.85 % / 24.25 % |
| *No. of non-H atoms* |  |  |
| Protein | 8,996 | 9,024 |
| Water | 50 | 17 |
| Others | 15 | -- |
| Ligands | 208 | 296 |
| *R.m.s. deviations* |  |  |
| Bond length (Å) | 0.006 | 0.005 |
| Bond angles (°) | 1.372 | 1.178 |
| Average B factors (Å<sup>2</sup>) | 70.0 | 66.0 |
| *Ramachandran plot* |  |  |
| Favored (%) | 97% | 96% |
| Allowed (%) | 3% | 4% |
| Outliers (%) | 0% | 0% |
| PDB code | 8RFN | 8RFM |

### 2.5 Data processing and structure determination

All data processing was carried out remotely, after the experiment, using the new Virtual Infrastructure for Scientific Analysis (VISA, https://visa.esrf.fr). Initial hit-finding was performed using a GPU version of NanoPeakCell (Coquelle et al. 2015) implementing Peakfinder8 algorithm (Kieffer et al. 2022). As hit finding parameters we used an SNR of 5.0 and a threshold of 1200. The Bragg reflections were integrated using the software package CrystFEL (White et al. 2012; White 2019) (version 0.10.1) after indexing was attempted with CrystFEL’s indexamajig using the algorithms XGANDALF (Gevorkov et al. 2019), MOSFLM (Powell 1999) and ASDF, in that order. The detector geometry file was further optimized by the *geoptimizer tool* (Yefanov et al. 2015).

MTZ files for phasing and refinement were generated by the CTRUNCATE program (Evans 2011) from the CCP4 software package (Winn et al. 2011) and a fraction of 5 % reflections were included in the generated R_free_ set. Initial phases of free hNQO1 and hNQO1-NAD^+^/H were obtained by molecular replacement with MOLREP (Vagin and Teplyakov 1997). For the free hNQO1, we used our recently published serial femtosecond crystallography structure (PDB 8C9J (Doppler et al. 2023)) as the search model. In the case of hNQO1-NAD^+^/H, we used the free hNQO1 structure from this study as the search model. The obtained models were refined using alternate cycles of automated refinement using non-crystallographic symmetry (NCS) with REFMAC5 (Murshudov et al. 2011) and manual inspection was performed with COOT (Emsley et al. 2010). The final refined structures were validated using the Protein Data Bank (PDB) validation service prior to deposition. The atomic coordinates and structure factors have been deposited in the PDB with accession codes PDBs 8RFN and 8RFM for the free hNQO1 and the complex hNQO1-NAD^+^/H, respectively. The final refinement statistics of both proteins are given in Table 1. Electron-density and POLDER maps were calculated with the MAPS tool in the PHENIX software suite (Adams et al. 2010). All figures of the free hNQO1 and hNQO1-NAD^+^/H structures presented in this manuscript were generated with UCSF CHIMERAX (version 1.2) (Meng et al. 2023) or PYMOL (version 2.4.0) (Schrödinger LLC).

### 2.6 Molecular dynamics simulations

Structural models of the hNQO1 protein were derived from former crystallographic data, PDB: 1D4A, selecting chains A and C to form the homodimer structure. The FAD cofactors were retained in their respective active sites. The positioning of NADH was manually adjusted based on similar protein structures with ligands (PDB: 2F1O, 1DXO) and the stereochemistry of the model early reported by Amzel and co-workers (Li et al. 1995), and similarly adjusted in both protomers. Before initiating any MD simulation, the structural models were meticulously prepared. This involved the removal of all crystallographic water molecules and the fine-tuning of the protonation of residues at pH of 7.0 utilizing the PROPKA software (Olsson et al. 2011). MD simulations were carried out using GROMACS 2018.4 software (Abraham et al. 2015), and the protein parameters were defined with the AMBER ff03 force field (Duan et al. 2003). Standard residue topologies were assigned using the GROMACS pdb2gmx tool, while FAD and NADH molecules were parametrized using ab initio methods. Atomic charges for each atom were determined through the restrained electrostatic potential (RESP) technique using Multiwfn (Lu and Chen 2012), following structure optimization at the HF/6-31G(d,p) level in Gaussian 09 (Frisch et al. 2009). These charges served as the initial parameters for the subsequent parametrization using the GAFF force field (Wang et al. 2006) via ACPYPE (Sousa da Silva and Vranken 2012). To set periodic boundary conditions, the system was enclosed within a rhombic dodecahedron box. Explicit water molecules were modeled using the TIP3P model. Neutralization of the system involved the replacement of random solute molecules with sodium ions. Prior to running production simulations, the studied systems underwent steepest descent minimization to alleviate close contacts or clashes. Equilibration was achieved through short 500 ps simulations under the NVT ensemble, with initial velocities generated according to a Boltzmann distribution at 310 K. Subsequently, a 500 ps simulation under the NPT ensemble at 1 atm was conducted. During both equilibration stages, harmonic potentials with a force constant of 1,000 kJ/(mol·nm) were applied to restrict the movement of the heavy atoms in the protein and ligand. Once the desired equilibrated conditions were met, the productive MD phase began with unrestrained positions for all atoms except hydrogen atoms, which were constrained using the LINCS algorithm (Hess 2008). Simulations were performed under an NPT ensemble at 310 K with a leap-frog integrator using 2 fs time steps, and data were collected every 10 ps. Long-range electrostatic interactions were computed using the Particle Mesh Ewald method, pressure was controlled via the Parrinello-Rahman method, and temperature equilibration was maintained through a modified Berendsen scheme. The simulations extended up to 200 ns, and each trajectory was replicated five times. All MD simulations were executed on the CIERZO supercomputer system at BIFI. Analysis of the simulations was performed using custom scripts using GROMACS tools, the MDTraj library (McGibbon et al. 2015) and PyMOL (Schrödinger LLC).

## 3 RESULTS

### 3.1 Experimental set-up at the new ID29 beamline

Serial crystallography data collection was performed at ID29, a brand-new energy tunable beamline at the 6 GeV storage ring of the new ESRF-EBS (Raimondi et al. 2023), the first high-energy fourth-generation synchrotron (Grenoble, France). A more detailed description of the beamline will be found in Orlans et al. (*Manuscript in preparation*). For the study presented here, we used the fixed target serial crystallography data collection platform developed at ID29. Crystal slurry was delivered to the X-ray beam using a small version of the SOS chip (Doak et al. 2018) and room temperature (290 K). The exposure area on the chip was 4 x 8 mm and was sampled collecting diffraction images every 10 μm steps apart, in x and y, respectively. The scanning speed, which is defined by the beam pulses repetition rate and spacing, was 2.3 mm/s, implying that the sample is moving 0.2 μm during the X-ray pulse. The sample-to-detector distance was 150 mm that was equivalent to 1.8 Å at the detector corners. This will change in the near future, so that data sets to resolutions as high as 1.2 Å in the detector corners at 1.072 Å wavelength can be achieved. A schematic view of the experimental set-up used at ID29 is shown in Figure 1B. For the free hNQO1 sample, a total of 21 μL (6 chips) of crystal slurry were consumed for a complete data set. In the case of NQO1 with NADH, we consumed 24.5 μL of crystal slurry from seven chips.

### 3.2 Room temperature crystal structures of free hNQO1 and hNQO1-NAD^+^/H

Despite the mylar sheets leading to three narrow rings on the detector, these were of sufficiently low intensity as to cause no difficulties in data analysis. A representative diffraction pattern for the complex hNQO1-NAD^+^/H is shown in Figure S2. The hit rate varied from 80 to 87 %. To prevent crystals from damaging effects, we collected data using a 10 x 10 μm raster scan spacing at the centers along horizontal and between rows, respectively. A total of 82,000 images were collected per chip in 8 min, which is too short to even observe dehydration effects so that no humidification was needed. Some images with multi-crystal diffraction were observed. All data collection statistics are listed in Table 1.

Microcrystals of free hNQO1 belonged to the space group P2_1_2_1_2_1_ with the following unit cell dimensions: a = 62.0 Å, b = 108.0 Å, c = 200.0 Å, α = β = γ = 90°, and diffracted to a resolution of ∼ 2.5 Å resolution. To solve the structure of the free hNQO1, a total of 492,000 frames were collected from five SOS chips (40 min), of which 302,781 were classified as hits by CrystFEL. About 33 % of the identified hits could then be indexed, giving rise to a total of 101,165 indexed, integrated, and merged lattices. The structure was solved by molecular replacement using the PDB entry 8C9J of the SFX structure of free hNQO1 previously reported by us (Doppler et al. 2023), as a search model without the FADs, water, and ion molecules. The structure was refined at a resolution of 2.7 Å with R_work_ and R_free_ of 19.87 % and 23.87 %, respectively. Figure S3 illustrates the two physiological homodimers in the asymmetric unit (ASU) related by a non-crystallographic two-fold axis of symmetry and showing the typical folding characteristic of hNQO1 (Bianchet, Faig, and Amzel 2004). In the case of the complex hNQO1-NAD^+^/H, the microcrystals belonged to the space group P2_1_2_1_2_1_ with shorter unit cell lengths dimensions: a = 61.1 Å, b = 106.8 Å, c = 196.0 Å, showing that crystals shrink upon the binding of the NADH. The same hit finding and indexing procedure used for free hNQO1 was applied for the case of hNQO1-NAD^+^/H. With 574,000 frames collected from six chips (48 min), an initial number of 344,574 hits were identified, of which ∼60 % were successfully indexed giving rise to individual 105,083 crystal lattices. The structure was solved by molecular replacement using the structure of hNQO1 (this study) as search model. The structure was refined at a resolution of 2.7 Å with R_work_ and R_free_ of 19.85 % and 24.25 %, respectively. The final data collection and refinement statistics for this structure are given in Table 1, while Figure 2 illustrates the two physiological homodimers in the ASU.

**Figure 2.**
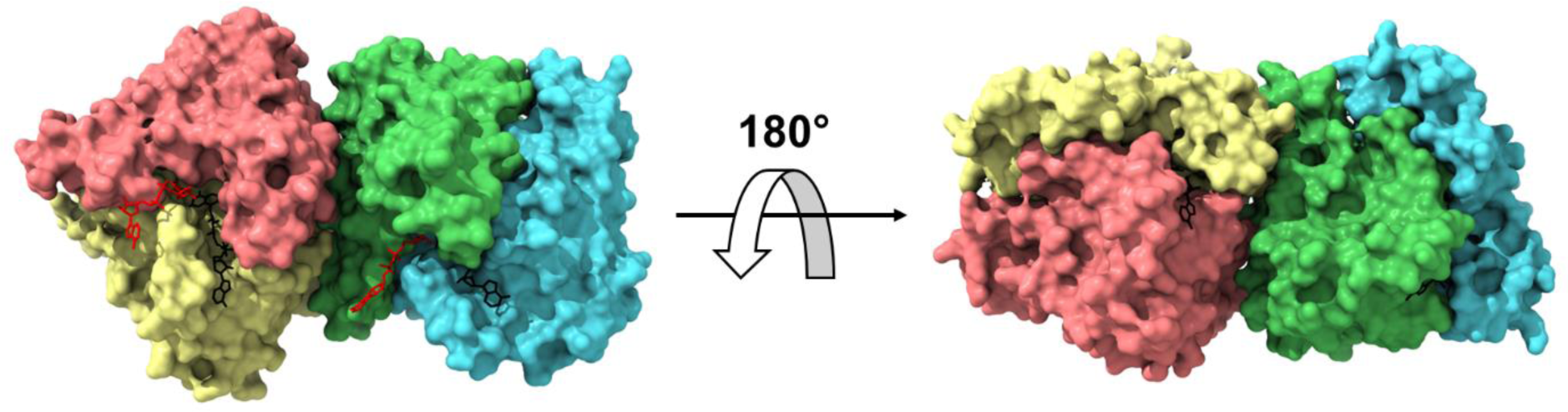
Room temperature structure of the complex hNQO1-NAD^+^/H obtained at ID29. **A)** The two homodimers of hNQO1 found in the ASU (A:B and C:D) are depicted as surface representation. The individual monomers are highlighted in salmon (chain A), yellow (chain B), green (chain C), and cyan (chain D). The coenzyme molecules bound to chain B (NAD^+^/H_B_) and chain D (NAD^+^/H_D_), are represented as sticks in red. The cofactor FADs are shown as black sticks.

The two structures could be modeled in the electron density from the N-terminus to the C-terminus without interruptions and with excellent stereochemistry (Table 1). The Ramachandran diagrams place nearly all residues within the favored and allowed regions, with only a few of them falling into the outlier region (Table 1). The resulting experimental maps were of excellent quality revealing, besides the presence of FADs, other solvent molecules such as water molecules: 76 in the free hNQO1 and 17 in the hNQO1-NAD^+^/H models. The high-quality of the hNQO1 structures can be assessed from the electron 2mF_o_–DF_c_ density maps shown for the catalytic site residues and the FAD cofactors (Figure 3A and S4 for hNQO1-NAD^+^/H and free hNQO1 structures, respectively). POLDER maps of the NAD^+^/H molecules in the final model of the complex are shown in Figure 3B and C, which clearly confirm the lack of model bias and, thus, their binding to the enzyme. Due to fact that the free hNQO1 and hNQO1-NAD^+^/H datasets are non-isomorphous (unit cell lengths a, b, and c vary 1 Å, 1.2 Å and 4 Å, respectively), F_o_-F_o_ maps were not calculated (see section 3.6 for an explanation of this). It is important to note that the NAD^+^/H molecules found in our structure are not bound to the same homodimer, but one of them is bound to one active site of one of the homodimers and the other one is bound to one of the active sites of the other homodimer in the ASU (Figure 2 and 3). Also, the exact disposition of both NAD^+^/H molecules differs from one to another.

**Figure 3.**
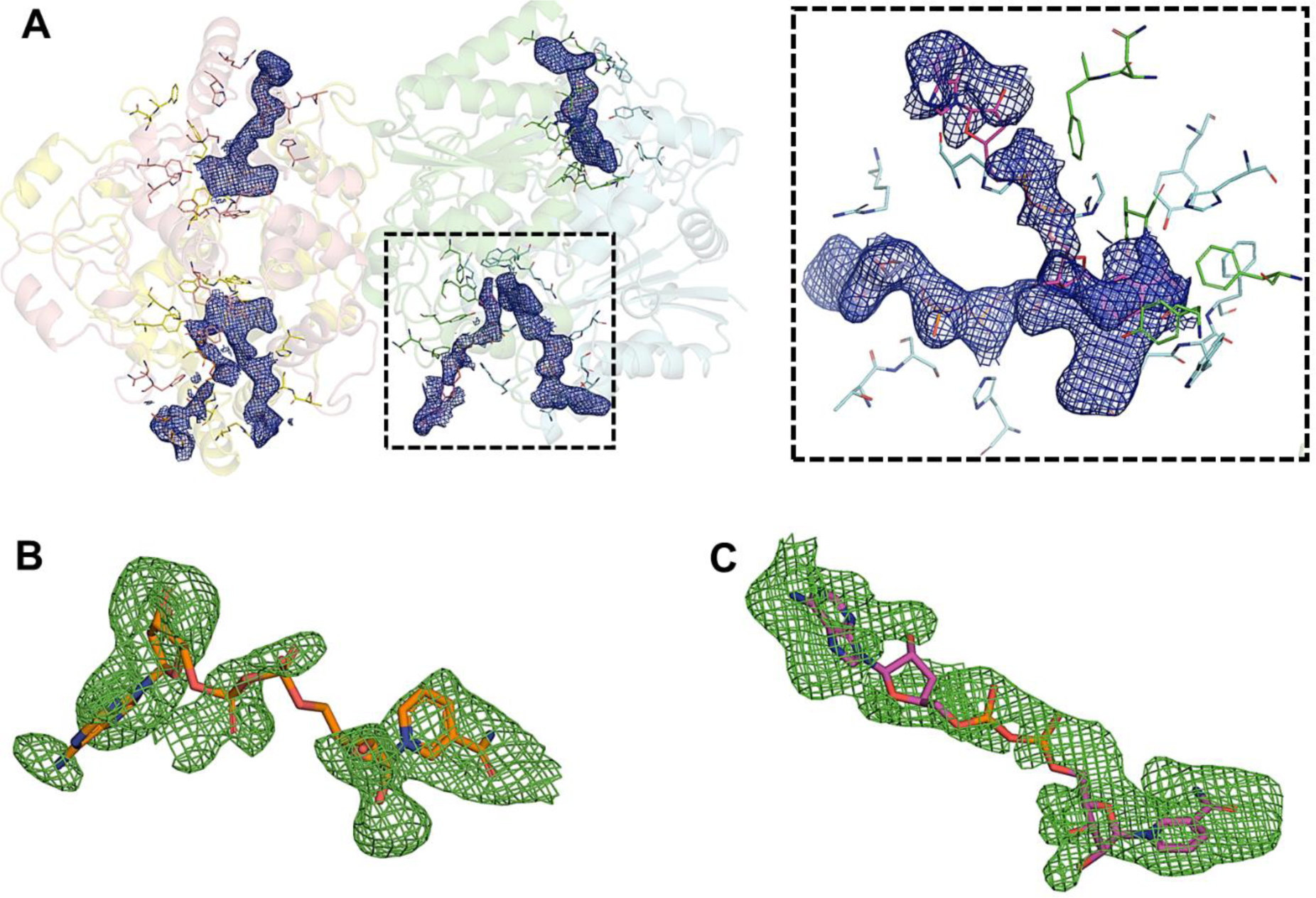
Electron density maps of the NQO1-NAD^+^/H structure. **A)** Cartoon representation of the two homodimers of hNQO1. Electron density maps 2mF_o_-DF_c_ contoured at 1 σ of the FAD and NAD^+^/H molecules are shown as blue meshes. Residues in the catalytic sites are shown as sticks. The right dash boxed panel shows a close-up view of the catalytic site dash boxed in the left panel. **B)** POLDER maps contoured at 3 σ of the NAD^+^/H molecule bound to chain D. **C)** POLDER maps contoured at 3 σ of the NAD^+^/H molecule bound to chain B.

### 3.3 Structural comparison of free hNQO1 and hNQO1-NAD^+^/H and with related structures

As mentioned above, the binding of the hydride donor NADH to hNQO1 led to an important reduction or shrinking of the unit cell lengths of up to 4 Å in the *c* dimension (Table 1), which shows the high flexibility of hNQO1 molecules within the crystal lattice. These differences in the unit cell dimensions are reflected when aligning the two homodimers of the free hNQO1 and hNQO1-NAD^+^/H structures (Figure 4A). A further analysis of the two structures with PISA server (Krissinel and Henrick 2005) was carried out in which the FAD and NAD^+^/H molecules were removed. This analysis showed that the number of total contacts in the complex hNQO1-NAD^+^/H (6,242) was significantly higher than that for the free structure (4,588), indicating that the binding of the NADH molecules rigidifies hNQO1. Consequently, the higher number of contacts observed in the complex led to an average reduction in the surface and buried areas of 1,000 Å^2^ and 100 Å^2^, respectively (Table S1). Also, the number of hydrogen bonds throughout the structures was 399 in the complex and 351 in the free structure. A further analysis of the contacts occurring at the homodimer interfaces shows that, overall, although the total number of inter- and intra-molecular hydrogen bonds is just slightly higher in the case of the complex (171 hydrogen bonds for the complex and 167 for the free structure) (Tables S2-S13), a redistribution of the hydrogen bonds across the interface was produced. Figure 4B and C illustrate the intermolecular hydrogen bonds observed for the two homodimers of the free hNQO1 and hNQO1-NAD^+^/H structures.

**Figure 4.**
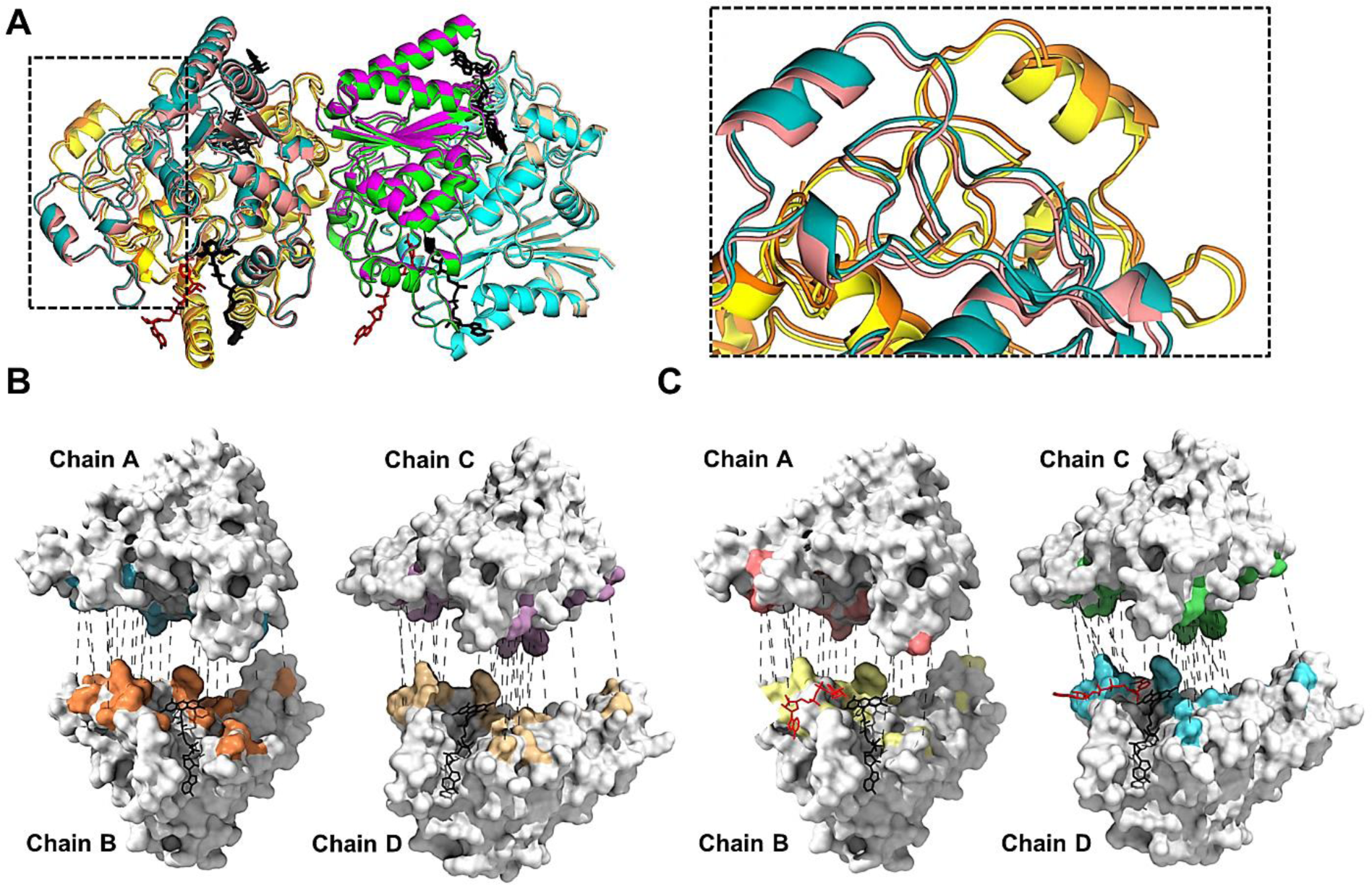
Structural comparison of the free hNQO1 and the hNQO1-NAD^+^/H structures. **A)** Cartoon representation of the superposition of the free hNQO1 and the complex hNQO1-NAD^+^/H structures. Color code used for the two structures is the same as that shown in Figures 2 and S3. All FAD and NAD^+^/H molecules are shown in stick representation in black and red, respectively. The right panel is a closer view of the inlet highlighted in the left panel. **B)** Hydrogen bonds (dashed grey lines) at the interfaces of the two homodimers in the free hNQO1 structure. **C)** Hydrogen bonds (dashed grey lines) at the interfaces of the two homodimers in the hNQO1-NAD^+^/H structure. Both in B) and C), protein monomers are shown as white surface representation with the interaction surfaces shown with the same color code as that in Figures 2 and S3, the protomers have been separated for clarity and all FAD and NAD^+^/H molecules are shown in stick representation in black and red, respectively.

A further evaluation of the hNQO1-NAD^+^/H complex was carried out by comparing it with previously reported crystal structures of hNQO1 both unliganded (free hNQO1 (this study), PDB 8C9J (Doppler et al. 2023), and PDB 1D4A (Faig et al. 2000)), and in complex with NADH (computational model (this study)), the inhibitor PMSF (8OK0 (Grieco et al. 2023), the natural substrate duroquinone (PDB 1DXO (Faig et al. 2000)), and various inhibitor molecules such as dicoumarol (PDB 5FUQ (Medina-Carmona et al. 2017)) and Cibracron blue (PDB 4CF6 (W. D. Lienhart et al. 2014)), as well as a prodrug molecule (PDBs 1GG5 (Faig et al. 2001)). Overall, all NQO1 homodimer structures aligned very well with each other, with RMSD values for all atoms between 0.301 Å and 0.852 Å and with an average value of 0.423 Å. A superimposition of all structures is shown in Figure S5A. As one could expect, the higher structural differences are mainly found in the loop regions, the solvent exposed regions, as well as in the catalytic site regions, where the side chains of several residues differ in conformation (Figure S5B).

### 3.4 Structural changes in the active site of hNQO1 upon binding of the hydride donor NADH

Since we have recently observed the existence of a high conformational heterogeneity of hNQO1 in the catalytic site with some of the residues being modeled in different conformations (Doppler et al. 2023), the structural changes of the catalytic sites of both structures, the free hNQO1 and hNQO1-NAD^+^/H, were analyzed individually. Analysis of the hNQO1 protein with CASTp program (Tian et al. 2018), shows that, overall, the catalytic pocket of hNQO1 buries a total area of 824.02 Å^2^ (volume 694.64 Å^3^) that is found at the dimer interface and comprises the binding pocket for the FAD molecule and the NAD^+^/H molecule.

#### The FAD binding site

Firstly, we carried out the analysis of the FAD catalytic pockets in the free hNQO1 and the hNQO1-NAD^+^/H structures. As described previously by Amzel and *co-workers* (Li et al. 1995; Bianchet, Faig, and Amzel 2004), the FADs are stabilized by establishing hydrogen bond interactions with residues from one of the monomers of the homodimer that are located, primarily, at the bottom of the active site. In that study, it was assumed that the interactions established by the FAD and the protein residues were equivalent in the two FAD binding sites, however, since our structure is a room temperature structure and based on previous results from our group (Doppler et al. 2023), we decided to analyze the FAD sites from all chains in the ASU. In fact, we have observed some differences between the two binding sites of the two homodimers of both structures (Figures 5, S6 and Table 2) that might be relevant to the hNQO1 function. Overall, the interactions between the FAD and the free hNQO1 include in general the two oxygen and all nitrogen atoms of the isoalloxazine ring of FAD and some of the residues located in one of the monomers including Trp105, Phe106, Gly149, Gly150, and His161. In addition, two of the oxygens of the ribitol fragment of the FAD are hydrogen bonded to Leu103 and Thr147. The diphosphate moiety hydrogen bonds, in general, with His11, Phe17 and Asn18 of the main protomer, and occasionally with Gln66 of the other protomer. Finally, the ribose/adenine moiety lies in a pocket composed of residues Thr15, Ser16, Phe17 and Arg200 of the same protomer together with Asn64 and Gln66 of the neighbor protomer, interacting mainly with Arg200 through the adenine, although interactions with Asn64 of the second protomer through the ribose are also observed. A more detailed illustration of all hydrogen bond interactions established by the FAD and the protein in free hNQO1 are shown in Figure S6 and Table 2. A similar interaction signature was observed in the FAD binding site of the hNQO1-NAD^+^/H structure (Figures 5, S7 and Table 2), which indicates that the binding of the NAD^+^/H molecules to the protein does not alter, at least not significantly, the FAD binding site. However, some variations in the hydrogen bond distances established between the FADs and the protein residues were observed when comparing both the free hNQO1 and the complex hNQO1-NAD^+^/H (Table 2).

**Figure 5.**
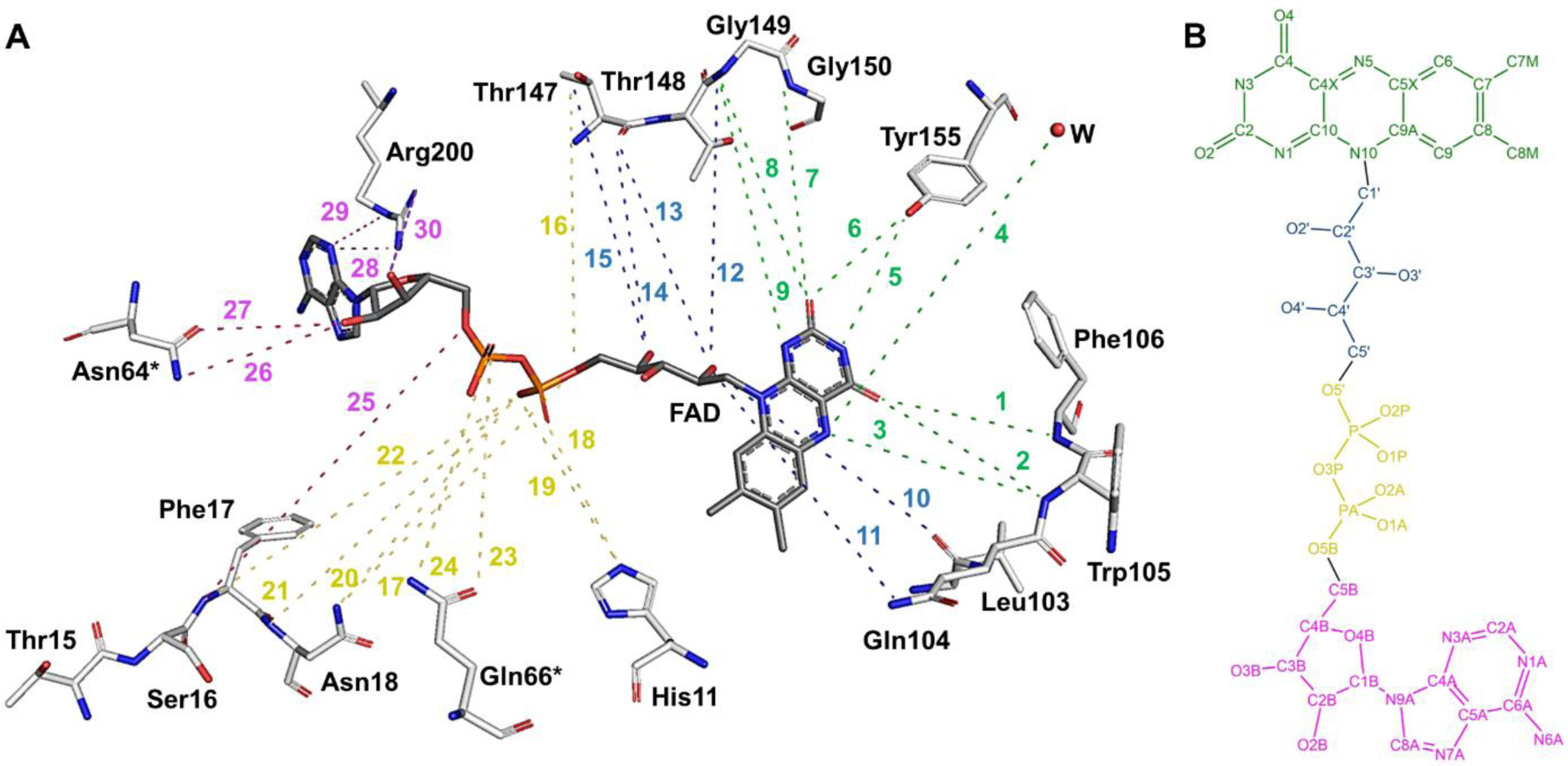
Schematic representation of all hydrogen bond interactions occurring at the FAD binding site. **A)** The binding site comprises most of the residues of one protomer and only two residues (Asn64* and Asn66*) of the other protomer. All interactions have been numbered from 1 to 30 (as listed in Table 2) and highlighted using the same color code as in Table 2: Green for all interactions at the isoalloxazine moiety, blue for all interactions at the ribitol moiety, yellow for all interactions at the phosphates moiety, and pink for all interactions at the ribose and adenine moieties. Important to note that, for clarity, all residues, the FAD and the water molecule have been moved and displaced from their original positions in the structure, as well as all the interactions have been exaggerated. **B)** 2D representation of the chemical structure of the FAD molecule. Atoms have been colored with the same code used for the interactions in A).

**Table 2:** Hydrogen bond interactions established between the FAD molecules and the protein residues in the free hNQO1 and the complex hNQO1-NAD^+^/H.

| # | FAD (atom) | Residue (atom) | Distance (Å) |  |  |  |  |  |  |  |
| --- | --- | --- | --- | --- | --- | --- | --- | --- | --- | --- |
|  |  |  | free hNQO1 |  |  |  | hNQO1-NAD<sup>+</sup>/H |  |  |  |
|  |  |  | Chain A | Chain B | Chain C | Chain D | Chain A | Chain B | Chain C | Chain D |
| 1 | O4 | Phe106[N] | 3.3 | 3.2 | 3.5 | 3.5 | 3.3 | 3.2 | 3.4 | 3.2 |
| 2 | O4 | Trp105[N] | 3.0 | 3.0 | 3.2 | 3.1 | 3.0 | 3.1 | 3.1 | 2.9 |
| 3 | N5 | Trp105[N] | 3.0 | 3.0 | 3.3 | 3.0 | 2.9 | 2.9 | 2.9 | 2.8 |
| 4 | N5 | Water[O] | -- | 3.3 | -- | 3.4 | -- | -- | -- | -- |
| 5 | N3 | Try155[OH] | 3.0 | 3.2 | 3.0 | 3.1 | 3.1 | 3.1 | 3.1 | 3.2 |
| 6 | O2 | Try155[OH] | 2.7 | 2.8 | 2.7 | 2.9 | 2.6 | 2.6 | 2.5 | 2.7 |
| 7 | O2 | Gly150[N] | 3.4 | 3.6 | 3.2 | 3.3 | 3.2 | 3.4 | 3.2 | 3.3 |
| 8 | O2 | Gly149[N] | 3.5 | 3.6 | 3.4 | 3.4 | 3.5 | 2.5 | 3.5 | 3.5 |
| 9 | N1 | Gly149[N] | 3.4 | -- | 3.3 | 3.5 | 3.4 | 3.4 | 3.3 | 3.4 |
| 10 | O2' | Gly149[N] | 3.4 | 3.5 | 3.6 | 3.5 | 3.2 | -- | 3.6 | 3.4 |
| 11 | O2' | Leu103[O] | 2.8 | 3.1 | 2.9 | 3.8 | 2.9 | 2.7 | 2.7 | 2.8 |
| 12 | O2' | Leu103[N] | -- | -- | 3.6 | -- | -- | -- | 3.6 | 3.6 |
| 13 | O2' | Thr147[O] | 3.3 | 3.5 | 3.2 | 3.3 | 3.2 | 3.4 | 3.2 | 3.2 |
| 14 | O4' | Thr147[O] | 2.9 | 3.0 | 3.0 | 3.0 | 2.9 | 3.1 | 3.0 | 2.9 |
| 15 | O4' | Thr147[OG1] | 2.7 | 2.7 | 2.6 | 2.6 | 2.5 | 2.6 | 2.3 | 2.4 |
| 16 | O5' | Thr147[OG1] | 3.3 | 3.3 | 3.4 | 3.4 | 3.1 | 3.2 | 3.1 | 3.2 |
| 17 | O5' | Asn18[ND2] | -- | -- | -- | -- | 3.6 | 3.5 | 3.5 | -- |
| 18 | O2P | His11[NE2] | 2.7 | 2.5 | 2.8 | 2.6 | 2.8 | 2.6 | 2.7 | 2.8 |
| 19 | O1P | His11[NE2] | -- | -- | 3.6 | 3.6 | -- | -- | 3.5 | 3.5 |
| 20 | O1P | Asn18[ND2] | 3.1 | 3.2 | 3.1 | 3.0 | 3.1 | 3.1 | 3.0 | 2.9 |
| 21 | O1P | Asn18[N] | 3.2 | 3.1 | 3.2 | 3.3 | 3.1 | 3.0 | 3.2 | 3.2 |
| 22 | O1P | Phe17[N] | 3.4 | 3.2 | 3.3 | 3.6 | 3.2 | 3.1 | 3.3 | 3.3 |
| 23 | O2A | Gln66[OE1] | -- | -- | -- | 2.7 | -- | -- | -- | 2.4 |
| 24 | O2A | Gln66[NE2] | 3.3 | 2.7 | 2.8 | -- | 3.1 | 3.0 | 3.0 | -- |
| 25 | O5B | Phe17[N] | 3.6 | 3.6 | -- | 3.5 | 3.5 | 3.5 | 3.5 | 3.4 |
| 26 | O2B | Asn64[ND2] | 3.5 | 3.2 | -- | 3.5 | -- | -- | -- | -- |
| 27 | O2B | Asn64[OD1] | -- | -- | -- | -- | -- | 3.1 | 3.6 | -- |
| 28 | N3A | Arg200[NH2] | -- | 3.1 | 3.3 | -- | -- | 2.8 | -- | -- |
| 29 | N3A | Arg200[NE] | 3.0 | 2.8 | -- | 2.9 | 3.1 | 2.9 | 3.1 | 3.1 |

A further comparison of the FAD binding sites of the two homodimers in the ASU of both the free hNQO1 and the complex hNQO1-NAD^+^/H is shown in Figure S8A and B. As can be seen, no significant differences were found for the protein residues. However, although all FAD molecules showed a similar conformation from one to another within the active sites of each structure, with the biggest differences found in the ribose and adenine moieties (Figure S8C and D), some relevant differences were found when comparing the FAD molecules between the free hNQO1 and the hNQO1-NAD^+^/H structures (Figures S9 and S10). As can be seen, the binding of the NADH induces conformational changes in all the FAD molecules, being more significant, overall, in the isoalloxazine ring (Figures S9) and the ribose and adenine moieties (Figure S10). Also, and rather surprisingly, the biggest changes are produced in the FADs of chains A and C, in which no NAD^+^/H molecules are bound.

Furthermore, water molecules also seem to play a relevant role in the stabilization of the FADs. Amzel and *co-workers* reported the presence of three water molecules coordinating the ribitol and diphosphate moieties of FAD (Bianchet, Faig, and Amzel 2004). Although none of these water molecules were found in the structures reported in this study, we found others. In this regard, in our free hNQO1 structure there is a conserved water molecule right above the isoalloxazine ring that interacts, in two of the chains (chains B and D), with its N5 atom (Figure 5). In the other two chains (chains A and C), this water molecule hydrogen bonds with Tyr128 (not shown). This is the first time these water molecules have been reported for hNQO1, which may have important implications in the stabilization of FAD and/or binding of NADH. Unlike in the free hNQO1, no water molecules were found in the FAD binding sites of the hNQO1-NAD^+^/H structure. The absence of such water molecules in chains B and D is justified by the presence of the NAD^+^/H molecules, which may displace them. In the case of the water molecule found in chains A and C, the absence of this water molecule could be justified by conformational changes caused in these active sites by NADH binding at the others.

#### The NAD^+^/H binding site

In our structure, we have captured the hydride donor NAD^+^/H in two different positions. We have observed a coenzyme molecule bound to the catalytic site of one of the homodimers through chain B (NAD^+^/H_B_), and another coenzyme molecule bound to the catalytic site of the other homodimer through chain D (NAD^+^/H_D_) (Figures 2 and 3A). The fact that the hydride donor molecule is found in just one catalytic site of each homodimer agrees with the negative cooperativity previously reported in literature during hydride transfer (Anoz-Carbonell et al. 2020). Because the interaction of the coenzyme NADH is transient, as it must leave the catalytic site upon hydride transfer so that the redox reaction continues, it must have fewer specific interactions with the protein than FAD.

##### 1) NAD^+^/H_D_ molecule

The electron density maps 2mF_o_-DF_c_ around the NAD^+^/H_D_ (Figure 3) indicate that the side face of the nicotinamide motif of the NAD^+^/H_D_ is stacked right on top of the middle ring of the isoalloxazine of FAD, thus establishing a Van der Waal interaction of 3.4 Å distance with FAD (Figure 6A). It is important to note that in this conformation the distance between the two atoms involved in the hydride exchange, the C4 of the nicotinamide (C4N) and the N5 of the FAD, is 3.4 Å. In addition, the NAD^+^/H_D_ is further stabilized by five more interactions that involve residues of the same protomer that binds FAD and residues of the other protomer (Figure 6A). The carboxyamide moiety makes hydrogen bonds with His161 and Tyr128. One hydroxyl of the ribose makes hydrogen bond with an oxygen of the ribose of the FAD molecule. The oxygen of the ribose also establishes an interaction with Tyr128. Lastly, the second phosphate moiety of NAD^+^/H interacts with His194. All these interactions are shown in Figure 6A, and their distances listed in Table 3. As the microcrystals, initially yellow color, turned clear right after they were mixed with the NADH and the coenzyme molecule is pointed with its A face towards the isoalloxazine, we assume that the NAD^+^/H_D_ molecule found at the catalytic site may presumably be NAD^+^, which has transferred the Pro-R hydride ion to the FAD, reducing it to FADH^-^ (Anoz-Carbonell et al. 2020).

**Figure 6.**
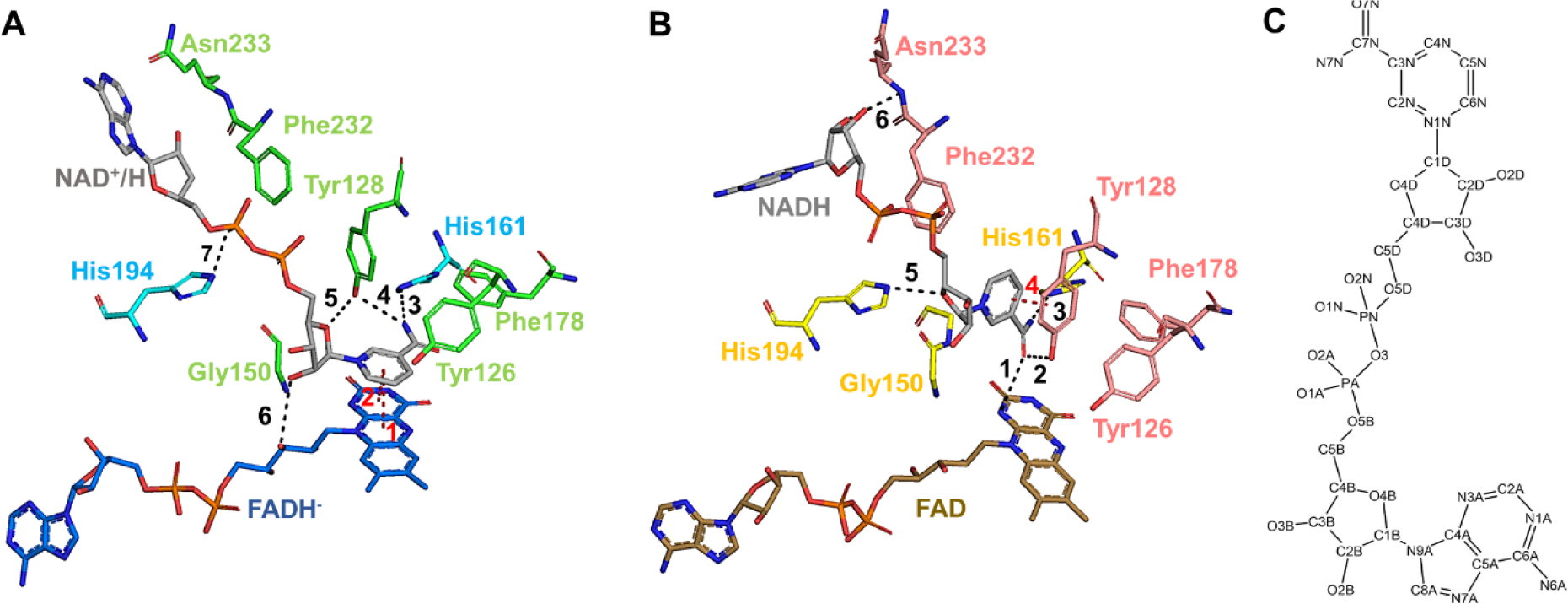
Hydrogen bond interactions at the NAD^+^/H binding sites in the two homodimers of the complex NQO1-NAD^+^/H. **A)** Stick representation of the molecule NAD^+^/H_D_, the FADs and all residues involved in the formation of the NAD^+^/H binding site of the homodimer C:D. **B)** Stick representation of the molecule NAD^+^/H_B_, the FADs and all residues involved in the formation of the NAD^+^/H binding site of the homodimer A:B. The color code of the protein residues is the same as that shown in Figure 2. The color code of the FADs is the same as that shown in Figure S6. All hydrogen bond interactions established between the NAD^+^/H molecules and both the protein residues and the FADs have been labeled and shown as black dashes. The *pi-interactions* are labeled and dashed in red. **C)** 2D representation of the chemical structure of the NADH molecule.

**Table 3:** Hydrogen bond and *pi-interactions* established between the NAD^+^/H_D_ molecule and the protein residues.

| Interaction* | Residue (chain) /atom | Residue (chain) /atom | Distance (Å) |
| --- | --- | --- | --- |
| **1\*\*** | **Nicotinamide ring** | **Isoalloxazine ring** | **3.6** |
| **2\*\*** | **Nicotinamide ring** | **Isoalloxazine ring** | **3.9** |
| 3 | NAD (D) / N7N | His161 (D) / NE2 | 3.5 |
| 4 | Tyr128 (C) / OH | NAD (D) / N7N | 3.1 |
| 5 | Tyr128 (C) / OH | NAD (D) / O4D | 3.3 |
| 6 | NAD (D) / O2D | FAD (D) / O3' | 3.7 |
| 7 | NAD (D) / O2A | His194 (D) / NE2 | 2.8 |
| * All interactions in this table correspond to those from Figure 6B |  |  |  |
| ** *pi*-interactions are highlighted in bold. |  |  |  |

##### 2) NAD^+^/H_B_ molecule

As mentioned above, there is an additional hydride donor molecule bound to chain B of the other homodimer (Figure 2). Figure 6B shows the interactions established between this NAD^+^/H_B_ molecule and the enzyme. The nicotinamide ring is parallel to the benzyl ring of the residue Tyr128, thus establishing a *pi-interaction*. The carboxyamide is interacting with His161 and the isoalloxazine ring of FAD through its N1F. Two more interactions are established between the two ribose sugars and protein residues His194 and Asn233. All these interactions are shown in Figure 6B, and their distances listed in Table 4. Based on our complex structure and the orientation of this hydride donor molecule, it is hard to know whether it has reacted (or not) with the FAD so that this molecule could be either on its way to the catalytic site (as NADH) or on its way out from it upon reaction (as NAD^+^).

**Table 4:** Hydrogen bond and *pi-interactions* established between the NAD^+^/H_B_ molecule and the protein residues.

| Interaction* | Residue (chain) /atom | Residue (chain) /atom | Distance (Å) |
| --- | --- | --- | --- |
| 1 | NAD (B) / N7N | FAD (B) / N1 | 3.6 |
| 2 | NAD (B) / N7N | Tyr128 (A) / OH | 2.9 |
| 3 | NAD (B) / N7N | His161 (B) / NE2 | 2.8 |
| **4\*\*** | **Nicotinamide ring** | **Tyr128 ring** | **3.3** |
| 5 | NAD (B) / O4D | His194 (B) / NE2 | 2.9 |
| 6 | Asn233 (A) / N | NAD (B) / O2B | 3.4 |
| * All interactions in this table correspond to those from Figure 6A |  |  |  |
| ** *pi*-interactions are highlighted in bold. |  |  |  |

### 3.5 Dynamic properties of FAD and NAD^+^/H and how they correlate with X-ray structures

The static picture provided by the crystal structures is not sufficient to fully understand highly dynamic processes such as the conformational changes occurring at and near the catalytic site during competent allocation of NADH in the active site to achieve subsequent hydride transfer from the C4N of the nicotinamide of NADH to the N5 of the FAD isoalloxazine. To further characterize the network interactions and the dynamic properties of the FAD and NAD^+^/H sites, the structure and dynamics of hNQO1 homodimers, in the free and in the NADH complex states, were investigated by MD simulations. Five MD replicates of 200 ns each were run for each structure, with overall atomic positions remaining close to the starting crystallographic structures with equilibrated average C_α_ RMSD fluctuations of 1.9 ± 0.2 Å for the free hNQO1 and 1.6 ± 0.2 Å for the hNQO1-NADH complex. Figure S11A shows the RMSDs for the Cα of the five replicates along the entire MD simulation for the free hNQO1and hNQO1-NADH models. Noticeably, the most relevant differences occur at the FAD and NADH binding sites (Figure 7 and Molecular movies V1 and V2). In good agreement with what we observed in the crystallographic results (Figure S5), residues at the FAD and, particularly, NADH binding sites show high flexibility. Among the highest ones are those observed at residues Asn64, Gln66 and Arg200, interacting with the ribose and adenine motifs of the FAD, and Tyr126, Tyr128, Phe232 and Asn233, interacting with the nicotinamide and adenine of NADH (Figures 7 and S11B and C, and Molecular movies V1 and V2). It is important to note that, in both, the free hNQO1 and the hNQO1-NADH models, the two catalytic sites behave as independent sites throughout the simulations, showing different protein-FAD and protein-NADH signatures in terms of interaction frequency and motional flexibility, as reflected in Figures 7 and S12-S14, and Tables S14-S18. A more detailed analysis of what happens during the simulations at both the FAD and NADH binding sites is described below.

**Figure 7.**
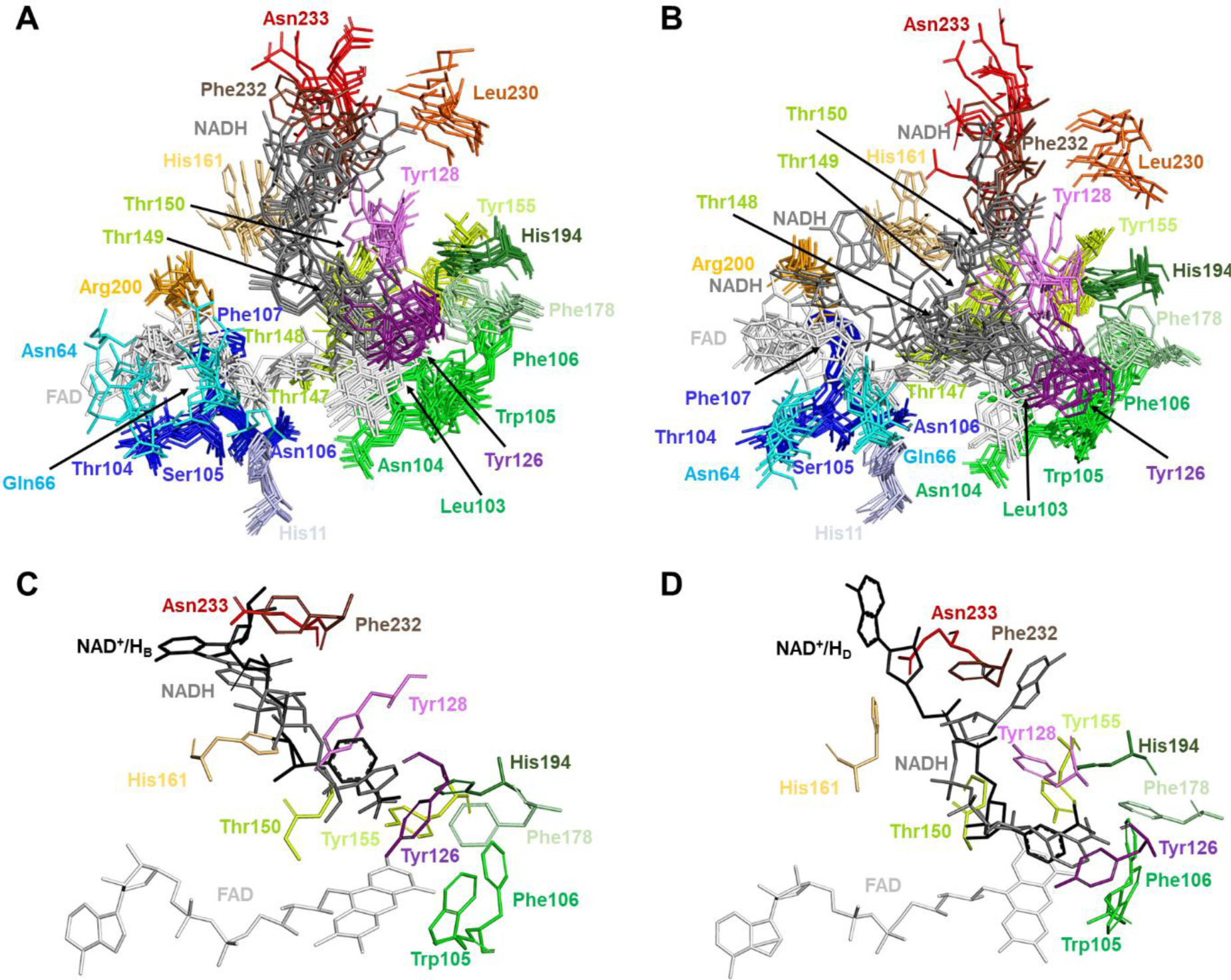
Structural dynamics of the FAD and NADH binding sites in the MD simulations. **A)** Superposition of the FAD and NADH binding sites for two randomly picked times from each of the five replicates. All residues comprising the FAD and NADH binding sites of the protomer at the active site labelled as 1 are shown in stick representation. **B)** Superposition of the FAD and NADH binding sites for two randomly picked times from each of the five replicates. All residues comprising the FAD and NADH binding sites of the protomer at the active site labelled as 2 are shown in stick representation. Both A) and B) show two randomly picked times from each of the five replicates analyzed in the simulations. **C)** Superposition of the NAD^+^/H_B_ molecule (black sticks) in the hNQO1-NAD^+^/H complex to the NADH binding site 1 at 60 ns simulation from replicate 4. **D)** Superposition of the NAD^+^/H_D_ molecule (black sticks) in the hNQO1-NAD^+^/H complex to the NADH binding site 1 at 30 ns simulation from replicate 2. All residues in C) and D) comprising the NAD^+^/H binding site are shown as sticks in the same color as that in A) and B).

#### The FAD binding site

Regarding the FAD, in agreement with the crystal structures, the position of residues interacting with the isoalloxazine ring is relatively stable, but there is quite flexibility in those interacting with the ribitol moiety and particularly with the phosphates, ribose and especially the adenine. All this can be seen in Tables S14 and S15, which illustrate how the polar interactions shown in Figure 5 for each of the fragments of FAD are maintained or not during the entire simulation. In addition, the adenine moiety moves a lot with no fixed position. In this regard, the Arg200 has been seen to interact sometimes with the ribose ring and sometimes with the adenine. Furthermore, during the simulations, the analysis of the RMSDs and RMSFs show that the rearrangement and movement of the two FADs are not symmetrical (Figures S12A and B, and S13A and B). In other words, the conformation/position of the FAD seen in one of the active sites of the protein is not the same as that seen in the other active site at the same time. In addition, in the presence of NADH, an overall decrease in FAD fluctuation is also observed (Figures S12A and B, and S13A and B), with again the FAD molecule of one catalytic site showing more fluctuation than the other.

#### The NADH binding site

Regarding NADH, the nicotinamide ring moves around quite a bit (Figures 7 and S14, Tables S16-S18). In general, it is almost kind of parallel to the isoalloxazine ring (Molecular movie V2). Nonetheless, it explores alternative conformations (Molecular movie V2), being even observed in some frames of replicate 4 at one of the active sites in positions more perpendicular to the FAD (Figure 7C), adopting, interestingly, a conformation that resembles that observed for the NAD^+^/H_B_ molecule of the crystal structure (Figure 6B). The position of the residues that interact with NADH changes quite a bit, being mainly the nicotinamide moiety that is anchored to the protein in a parallel conformation to the FAD isoalloxazine (Figure 7D and Molecular movies V1 and V2), like what was observed for the NAD^+^/H_D_ molecule of the crystal structure (Figure 6A). Same as in the crystal structures, Tyr126 and Tyr128 show a high flexibility, but the OH of Tyr126 usually stacks against the nicotinamide ring along with residues Phe178 and Trp105, keeping the nicotinamide buried (Figure S15). Also, Tyr128 pairs its OH with opposite nicotinamide and stays generally in that position, although sometimes it flips over and interacts its OH with the O7N and/or N7N atoms of NADH (Molecular movie V2). The rest of the NADH molecule is very flexible, exploring multiple conformations, and having few polar interactions with protein residues, as well as with the FAD. This was also observed in the crystal structure of the complex hNQO1-NAD^+^/H (this study). All this can be seen in Figure S14 and Tables S16-S18, which illustrate how the polar and *pi-stacking* interactions shown in Figure 6 for the crystal structure of the hNQO1-NAD^+^/H, vary during the entire simulation. Furthermore, as observed with FAD, the conformation/position of the NADH molecule, and particularly of its nicotinamide regarding the isoalloxazine of FAD, is not symmetrical throughout the simulations, thus showing differential fluctuation at one of the catalytic sites than at the other (Figure S14).

### 3.6 Crystal packing of hNQO1 homodimers in microcrystals differs from that in large crystals

A further evaluation of the hNQO1 crystal packing was carried out by comparing the two hNQO1 crystal structures presented in this study with the crystal structures recently published by our group for hNQO1 in the absence of NADH coenzyme (PDB 8C9J, (Doppler et al. 2023)) and hNQO1 in complex with the inhibitor PMSF (PDB 8OK0) (Grieco et al. 2023). Although all these structures were obtained from crystals grown in the exact same crystallization conditions, the rearrangement of the double hNQO1 homodimer observed in the ASU for the structure with PMSF differs from that observed in structures from microcrystals (Figure 8A). Regardless of the crystal structure, a structural alignment of the individual two hNQO1 homodimers leads to all hNQO1 homodimers aligning very well with each other (not shown). However, when the two homodimers are subjected, at once, to a structural alignment, clear differences arise in that one of the homodimers always aligns perfectly well while the other does not (Figure 8A). A visual analysis of the two homodimers showed that the homodimers rearrangement in the microcrystals differs significantly from that observed in the large crystals.

**Figure 8.**
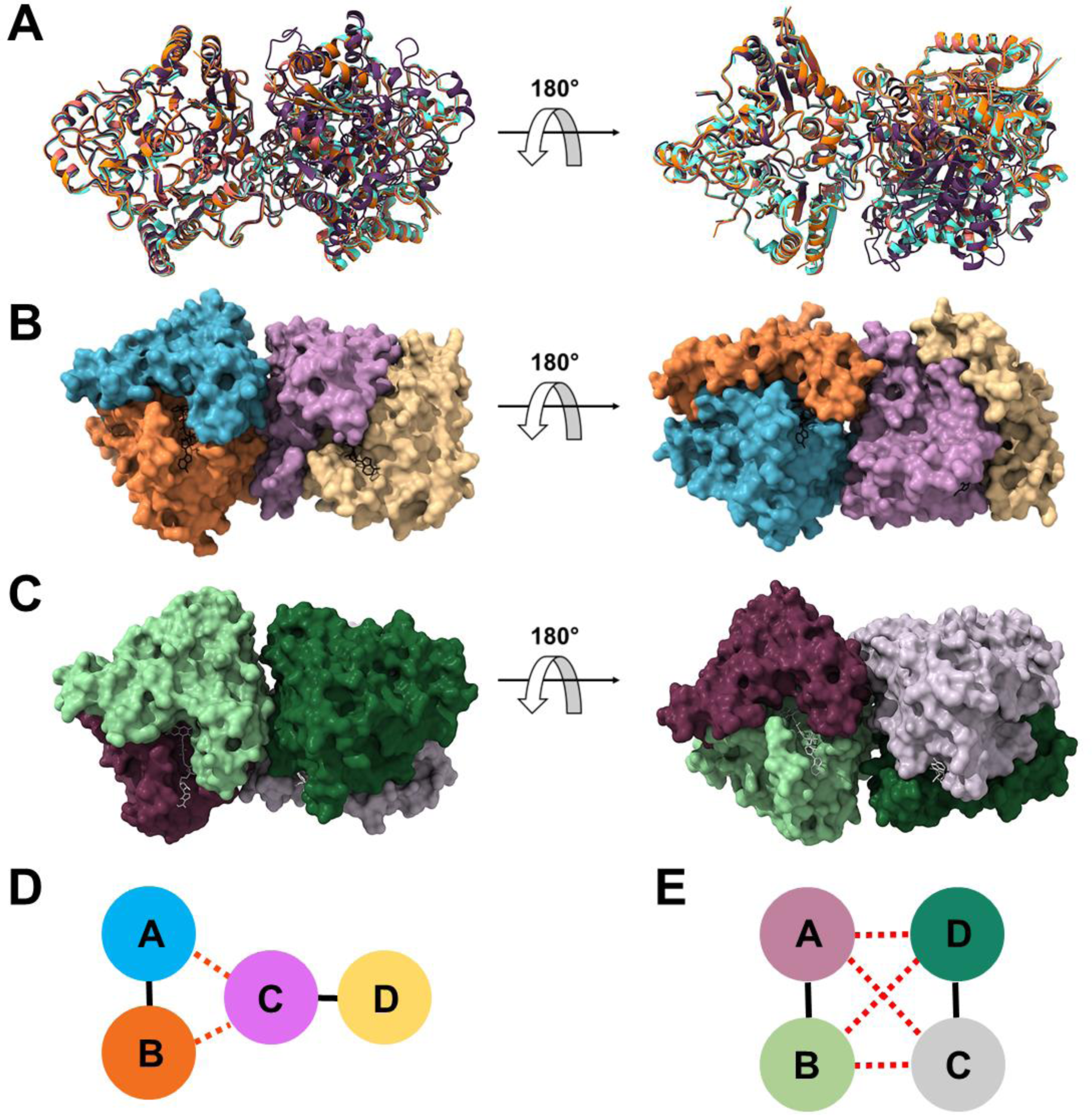
Comparison of the packing in the two homodimers of hNQO1 from different structures. **A)** Superposition of the two homodimers in the microcrystals of the free hNQO1 (orange) and hNQO1-NAD^+^/H (red), this study, and PDB 8C9J (light blue) (Doppler et al. 2023), to the two homodimers in the large crystals of PDB 8OK0 (purple) (Grieco et al. 2023). **B)** Surface representation of the free hNQO1 structure (this study). The FADs are shown as black sticks. **C)** Surface representation of the hNQO1 in complex with PMSF (PDB 8OK0) (Grieco et al. 2023). The FADs are shown as grey sticks. **D)** Schematic diagram of the intra-homodimer (solid black lines) and inter-homodimer (dashed red lines) interactions established in the microcrystals. As shown in A), the two homodimers are stabilized through interactions between chains A and B of one of the homodimers and chain C of the other homodimer with no participation of monomer D (yellow). **E)** Schematic diagram of the intra-homodimer (solid black lines) and inter-homodimer (dashed red lines) interactions established in the large crystals. As shown in B), the two homodimers are stabilized through interactions between chains A and B of one of the homodimers and chains C and D of the other homodimer.

A further analysis of the homodimers interface with the PISA server (Krissinel and Henrick 2005) shows that the two homodimers in the room temperature structures of the free hNQO1 (PDB 8C9J (Doppler et al. 2023); and this study) and the complex hNQO1-NAD^+^/H (this study), enclose a surface area that is similar in size (930 Å^2^ on average), while the cryo-structure of hNQO1 in complex with PMSF (PDB 8OK0, (Grieco et al. 2023)) buries a smaller surface area of 775 Å^2^. From this analysis, we have also found out that there are significant differences in the residues that participate in the interfaces between the two homodimers. As can be seen in Figure 8B for the homodimers in the microcrystals, they are kept by interactions established by the two protomers of one of the homodimers and just one protomer of the other homodimer with no participation from the other one. However, in the case of the large and cryo-cooled crystals, the two homodimers are kept through interactions established by all the protomers of the two homodimers (Figure 8C). A schematic diagram of how this homodimer rearrangement occurs in the microcrystals and in the large crystals can be seen better in Figure 8D and E. Putting all this together, we hypothesize that the way we produce our microcrystals at high supersaturation reached very fast increasing the nucleation rate, influences the way the two homodimers are rearranged within the crystals.

To further confirm this hypothesis, the crystal packing analysis was extended to other structures of hNQO1 in which large crystals were obtained from very similar or identical crystallization conditions such as PDB 5A4K (Lienhart et al. 2017) and 5EA2 (Pidugu et al. 2016) for the free hNQO1, and 6FY4 (Strandback et al. 2020) for the hNQO1 in complex with the ligand BPPSA, a small molecular chaperone. All these three structures showed two homodimers in the ASU as well. A comparison of these three structures with that of hNQO1 in complex with PMSF (PDB 8OK0) (Grieco et al. 2023), shows that the homodimers exhibit the same rearrangement within the large crystals (Figure S17A), this being different from the homodimer rearrangement observed in the microcrystals (Figure S17B).

## 4 DISCUSSION

### 4.1 hNQO1 is fully active in microcrystals showing they are suitable for time-resolved experiments

Proteins are not static entities but rather they fluctuate between alternative conformational states defined by a complex energy landscape (Frauenfelder, Sligar, and Wolynes 1991) to accomplish their biological functions. The redistribution or ‘population shift’ of the conformational ensemble is the basis of cooperativity and allosterism (Hilser 2010; Motlagh et al. 2014). Structural and biochemical studies of oxygen-carrying tetrameric hemoglobins have had a pivotal role in explaining the phenomenon of cooperativity and allostery (Kavanaugh, Rogers, and Arnone 2005). X-ray crystallography has traditionally been viewed as a static technique, with limited applicability to allostery and cooperativity. The reason for that is because, typically, protein crystals tend to be seen as static entities and therefore unsuitable for studying allosteric and cooperativity, given that these events involve conformational changes. However, it has been known for a long time that most enzymes remain catalytically active within crystals (Doscher and Richards 1963), although the environment within a crystal lattice has little resemblance to the actual cellular environment of a protein. Proteins are highly dynamic molecules, but even though the lattice restrains their accessible conformational space, they retain a remarkable degree of plasticity (Forneris, Burnley, and Gros 2014) and often display surprisingly large-scale conformational changes during ligand binding (Stagno et al. 2017). The accessible conformational space depends on how the protein is packed in a given crystal form, and analysis of multiple crystal forms gives a more complete picture of the conformational landscape (Tyka et al. 2011).

However, owing to the convergence of several recent experimental and technical developments, X-ray crystallography is increasingly well-positioned to provide insights into the connection between protein flexibility and function. In this regard, it has been shown that crystal packing in microcrystals is key to studying reaction mechanisms by mix-and-inject serial crystallography technology (Olmos et al. 2018; Schmidt 2020). To this end, in the study presented here, we report the crystallographic structures of the hNQO1 in its free form and in complex with the hydride donor NADH as obtained from microcrystals at room temperature by serial crystallography experiments. We have noticed that, for crystals grown in the same crystallization conditions, the rearrangement of the two hNQO1 homodimers in the microcrystals is different to that observed in hNQO1’s large crystals (Figure 8). Although this has never been reported in the literature, we believe that, for this case, the way we grow the microcrystals, much faster than we typically used to grow large crystals, affects the packing of the molecules within the crystals. Also, because of this, the catalytic site is more accessible in microcrystals than in large crystals, enabling the large hydride donor molecule to diffuse into the crystals and, thus, allowing us to determine the structure of the complex hNQO1-NAD^+^/H.

A question that may arise from our hNQO1-NAD^+^/H structure is whether the two NAD^+^/H molecules found at the catalytic pockets (NAD^+^/H_B_ and NAD^+^/H_D_) reacted with the enzyme. Based on the static structures presented in our study, it is hard to reach a conclusion. To assess whether hNQO1 protein remains enzymatically active within microcrystals grown from the batch-with-agitation method, the microcrystals, initially yellow due to the presence of the oxidized FAD cofactor, turned clear immediately upon mixing them with NADH. This color change confirms the reduction of the enzyme by NADH and, thus, indicates that hNQO1 is catalytically active within crystals. Based on this finding we could assume that the reaction has finished at least at one of the catalytic sites, so the hydride transfer from NADH to FAD has occurred. Thus, we hypothesized that at least one but probably the two NADH molecules (NAD^+^/H_B_ and NAD^+^/H_D_) must be NAD^+^ and the two FAD molecules FADH^-^.

### 4.2 The binding of the coenzyme NADH reduces the plasticity of hNQO1 in microcrystals

So far, the only structure reported on the literature (but not deposited in PDB) of a NQO1 protein in complex with the coenzyme was produced with the NADP^+^ oxidized form of the coenzyme (Li et al. 1995). NADP^+^ is not the NQO1’s substrate but the product of the reductive half-reaction produced after the reduced substrate, NADPH, transfers a hydride to the enzyme FAD cofactor. With that said, the crystal structure of the complex hNQO1-NAD^+^/H reported in this study is, to the best of our knowledge, the first structure of a natural relevant substrate, NADH in this case, being observed in complex with an NQO1 enzymés family. An overall comparative structural analysis of the hNQO1 in the crystal structures of the free hNQO1 with the complex hNQO1-NAD^+^/H, revealed that the binding of the hydride donor NADH reduces the plasticity of the enzyme homodimers, as reflected by the shrinking of the unit cell dimensions (Table 1). A more careful analysis reveals that, upon binding of the NADH, there is a rearrangement in the signature of the hydrogen bond interactions (in number and length) observed at the catalytic sites between the protein residues and the FAD molecules in the two homodimers (Figures 5, S6 and S7, and Table 2). More interestingly, this structural rearrangement propagates across the homodimeric interfaces of the two homodimers, as reflected by the number of inter- and intra-molecular hydrogen bonding interactions (Figure 4 and Tables S2-S13).

### 4.3 New structural and dynamics insights into the function of hNQO1

hNQO1 catalyzes the two-electron reduction of quinones to hydro-quinones by a two-step redox mechanism (Anoz-Carbonell et al. 2020), which has been widely studied by multiple biophysical and biochemical techniques and has been shown to use cooperative regulation (Pey, Megarity, and Timson 2014; Clavería-Gimeno, Velazquez-Campoy, and Pey 2017; Pey 2018; Pacheco-Garcia et al. 2021; Anoz-Carbonell et al. 2020). Essentially, cooperativity is manifested when the binding of a ligand to a protein alters the affinity for subsequent binding of the same or a different ligand. In this process a chemical or molecular signal is transmitted from one site in a protein to another to alter its structure and/or dynamics and therefore its function. Many protein enzymes, like hNQO1, utilize cooperative regulation to modulate their catalytic activity by binding to a regulatory molecule, an activator or inhibitor, that modifies binding sites for reaction substrates. However, it remains unclear how to decipher which local regions of a protein structure are conformationally coupled to each other. Therefore, the understanding of the molecular mechanism of hNQO1 has been limited by the complexity of its redox mechanism and by the difficulty in obtaining structural and dynamic information on the intermediate states by standard macromolecular crystallography.

Interestingly, the structure of the complex hNQO1-NAD^+^/H shows a molecule of the hydride donor bound to just one of the protomers of each of the two homodimers found in the ASU (Figure 2), which agrees well with the negative cooperativity from studies of hNQO1 in solution (Pey, Megarity, and Timson 2014; Clavería-Gimeno, Velazquez-Campoy, and Pey 2017; Pey 2018; Pacheco-Garcia et al. 2021; Anoz-Carbonell et al. 2020). Furthermore, the two molecules of the hydride donor (named NAD^+^/H_B_ and NAD^+^/H_D_ in this study) have been captured in two different conformations.

As reported by our group recently for the free hNQO1, the residues Tyr128 and Phe232, which gate the pocket, swinging in and out, showed a high-flexibility and thus were modeled in various conformations (Doppler et al. 2023). In the structures presented here, these two residues also showed a high flexibility (Figure S5C). This observation was also made in the MD simulations, in which more dramatic changes are particularly seen for residues Tyr128 and Phe232. In contrast to crystal structures, the Tyr126 was also observed to be highly flexible during the simulations. It is important to note that, these three residues, Tyr126, Ty128 and Ph232, are strictly conserved and envisaged as key players in the enzymatic function of NQO1 (Asher et al. 2005; Li et al. 1995; Pandey et al. 2020).

In addition to the structural changes observed at the protein level, conformational changes have been observed at the FAD level upon binding of NADH, being these changes more relevant for the isoalloxazine ring and the ribose and adenine moieties (Figures S9 and S10), especially in those protomers where no NAD^+^/H molecules were found. In a recent study by Maestre-Reyna and co-workers, they reported the first structural description of the microscopic processes of the photoreduction of a DNA photolyase by using pump-probe time-resolved serial crystallography at an X-ray free electron laser (XFEL) (Maestre-Reyna et al. 2022). DNA photolyases are flavoproteins containing, like NQO1, an FAD molecule as cofactor. In their study, they observed big geometric changes of the isoalloxazine moiety of FAD molecule during its reduction to FADH^-^ upon light excitation (Maestre-Reyna et al. 2022). In the crystal structure of our complex hNQO1-NAD^+^/H, although conformational changes have been observed for the isoalloxazine moiety (Figure S9), these are too subtle, which may indicate that the reaction has simply finished. However, similar geometric changes of the isoalloxazine have been occasionally captured in some snapshots of our MD simulations, even though they only account of FAD molecules in their oxidized state (Molecular movie V1 and V2). So that, to investigate these changes in the isoalloxazine experimentally, time-resolved experiments at XFELs would be needed.

It is well known that computational approaches complement experimental methods and provide powerful tools to study cooperativity, with MD simulations providing dynamic details. The large number of snapshots generated from MD simulations captures the motion of the proteins, thus providing insights into the population shift of the protein conformational ensemble. To this end, a more careful analysis of the complex of hNQO1 with NADH was carried out using MD simulations. It is important to note that both our structural and computational results match perfectly well. In this regard, from this analysis we have noticed that both the FAD and the NADH molecules are extremely dynamic even when bound to the enzyme (Figures 7, S12 and S13). Regarding the FAD, the MD simulations show, overall, an extremely more flexible FAD than that observed in the crystals, particularly in its adenine nucleotide moiety. In the case of the NADH molecules, they are mainly anchored to the protein through its nicotinamide moiety, which stacks against the FAD isoalloxazine in a binding pocket formed by numerous aromatic residues (Figures S14 and S15), being the rest of the NADH molecule highly flexible and hardly interacting with protein residues throughout the simulations (Tables S16, S17 and S18). This high plasticity of the NADH is also reflected in our crystals and explains why the electron density surrounding the two NAD^+^/H molecules is either poor or lacking, especially in the ribitol and the phosphates moieties, as well as the ribose next to the adenine (Figure 3). This effect is more pronounced in the case of the NAD^+^/H_B_ molecule.

## 5 CONCLUSIONS

The crystal structures reported in this study for hNQO1 are the first structures coming out from the new beamline ID29. More importantly, both structural results and MD simulations have supported that the two catalytic sites of hNQO1 act cooperatively and display highly collective inter-domain and inter-monomer communication and dynamics. This coupled network of amino acids is sensitive to ligand binding. In fact, the binding of NADH significantly decreases protein dynamics and stabilizes hNQO1 especially at the dimer core and interface, providing the first structural evidence that the hNQO1 functional cooperativity is driven by structural communication between the active sites through long-range propagation of cooperative effects across the hNQO1 protein structure. The flexibility and negative cooperativity of hNQO1 seen in the room temperature structure of the complex and the MD simulations are important features in obtaining a more complete picture of the function and dynamics of this enzyme, and the approaches delineated are critical steps needed for pursuing time-resolved crystallographic studies. In this sense, we are currently developing time-resolved experiments with X-ray free electron lasers and the results from these experiments will be published elsewhere. Therefore, understanding the dynamics of the redox mechanism of hNQO1 and of its interaction with ligands is critical to unravel its role as an antioxidant and as a potential target to treat common diseases in which hNQO1 is involved, advancing the design of new, more potent, and effective inhibitors that can be used in the clinical setting.

## DATA AVAILABILITY

The structural data for the free hNQO1 and the complex hNQO1-NAD^+^/H generated in this study are available in the Protein Data Bank repository (https://www.rcsb.org/) under accession codes PDB 8RFN and 8RFM, respectively.

## SUPPLEMENTAL INFORMATION

The following information is provided as supplemental: Figures S1-S17, Tables S1-S18, and molecular movies V1 and V2.

## AUTHOR CONTRIBUTIONS

JMMG and MM conceived the experiments. SBa, JO, DdS, ALP, MM and JMMG designed the experiments. AG, JLPG, and JAG produced and crystallized the protein. SBa, JO, DdS, AG, JAG, and JMMG participated in the beamtime experiment. SBa, JO, and DdS participated as beamline scientists. SBa, JO, DdS, AG, and JMMG analyzed diffraction data. AG and JMMG solved the structures. SBo, and MM performed MD simulations. AG, SBo, MM and JMMG made the figures of the manuscript. AG, SBo, ALP, MM and JMMG wrote the manuscript with input from all other co-authors.

## ACKNOWLEDGEMENTS

JO, SBa and DdS would like to acknowledge members of the ESRF-EMBL Joint Structural Biology Group (JSBG) and other support groups in the commissioning and operation of the new ID29. The results reported here were collected from the BAG proposal number MX2427. SBo and MM would like to acknowledge the use of the instrumentation provided by the National Facility ICTS, Centro de Supercomputación de Aragón (CESAR), at Universidad de Zaragoza.

## REFERENCES

Abraham, Mark James, Teemu Murtola, Roland Schulz, Szilárd Páll, Jeremy C Smith, Berk Hess, and Erik Lindahl. 2015. “GROMACS: High Performance Molecular Simulations through Multi-Level Parallelism from Laptops to Supercomputers.” SoftwareX 1–2: 19–25. 10.1016/j.softx.2015.06.001.

Adams, P D, P V Afonine, G Bunkoczi, V B Chen, I W Davis, N Echols, J J Headd, et al. 2010. “PHENIX: A Comprehensive Python-Based System for Macromolecular Structure Solution.” Acta Crystallogr D Biol Crystallogr 66 (Pt 2): 213–21. 10.1107/S0907444909052925.

Anoz-Carbonell, Ernesto, David J. Timson, Angel L. Pey, and Milagros Medina. 2020. “The Catalytic Cycle of the Antioxidant and Cancer-Associated Human NQO1 Enzyme: Hydride Transfer, Conformational Dynamics and Functional Cooperativity.” Antioxidants 9 (9): 1–22. 10.3390/antiox9090772.

Asher, Gad, Orly Dym, Peter Tsvetkov, Julia Adler, and Yosef Shaul. 2006. “The Crystal Structure of NAD(P)H Quinone Oxidoreductase 1 in Complex with Its Potent Inhibitor Dicoumarol.” Biochemistry 45 (20): 6372–78. 10.1021/bi0600087.

Asher, Gad, Peter Tsvetkov, Chaim Kahana, and Yosef Shaul. 2005. “A Mechanism of Ubiquitin-Independent Proteasomal Degradation of the Tumor Suppressors P53 and P73.” Genes and Development 19 (3): 316–21. 10.1101/gad.319905.

Beaver, Sarah K., Noel Mesa-Torres, Angel L. Pey, and David J. Timson. 2019. “NQO1: A Target for the Treatment of Cancer and Neurological Diseases, and a Model to Understand Loss of Function Disease Mechanisms.” Biochimica et Biophysica Acta - Proteins and Proteomics 1867 (7–8): 663–76. 10.1016/j.bbapap.2019.05.002.

Betancor-Fernández, Isabel, David J Timson, Eduardo Salido, and Angel L Pey. 2018. “Natural (and Unnatural) Small Molecules as Pharmacological Chaperones and Inhibitors in Cancer.” In Targeting Trafficking in Drug Development, edited by Alfredo Ulloa-Aguirre and Ya-Xiong Tao, 155–90. Cham: Springer International Publishing. 10.1007/164_2017_55.

Bianchet, Mario A., Margarita Faig, and L. Mario Amzel. 2004. “Structure and Mechanism of NAD[P]H:Quinone Acceptor Oxidoreductases (NQO).” Methods in Enzymology 382 (1997): 144–74. 10.1016/S0076-6879(04)82009-3.

Clavería-Gimeno, Rafael, Adrian Velazquez-Campoy, and Angel Luis Pey. 2017. “Thermodynamics of Cooperative Binding of FAD to Human NQO1: Implications to Understanding Cofactor-Dependent Function and Stability of the Flavoproteome.” Archives of Biochemistry and Biophysics 636 (September): 17–27. 10.1016/j.abb.2017.10.020.

Coquelle, Nicolas, Aaron S. Brewster, Ulrike Kapp, Anastasya Shilova, Britta Weinhausen, Manfred Burghammer, and Jacques Philippe Colletier. 2015. “Raster-Scanning Serial Protein Crystallography Using Micro- and Nano-Focused Synchrotron Beams.” Acta Crystallographica Section D: Biological Crystallography 71: 1184–96. 10.1107/S1399004715004514.

Doak, R Bruce, Gabriela Nass Kovacs, Alexander Gorel, Lutz Foucar, Thomas R M Barends, Marie Luise Grünbein, Mario Hilpert, et al. 2018. “Crystallography on a Chip – without the Chip: Sheet-on-Sheet Sandwich.” Acta Crystallographica Section D Structural Biology 74 (10): 1000–1007. 10.1107/S2059798318011634.

Doppler, Diandra, Mukul Sonker, Ana Egatz-gomez, Alice Grieco, Sahba Zaare, Rebecca Jernigan, Jose Domingo Meza-aguilar, et al. 2023. “Lab on a Chip Modular Droplet Injector for Sample Conservation Conformational Heterogeneity in the Disease-Associated NQO1 Enzyme †.” 10.1039/d3lc00176h.

Doscher, Marilynn S., and Frederic M. Richards. 1963. “The Activity of an Enzyme in the Crystalline State: Ribonuclease S.” Journal of Biological Chemistry 238 (7): 2399–2406. 10.1016/s0021-9258(19)67984-6.

Duan, Yong, Chun Wu, Shibasish Chowdhury, Mathew C Lee, Guoming Xiong, Wei Zhang, Rong Yang, et al. 2003. “A Point-Charge Force Field for Molecular Mechanics Simulations of Proteins Based on Condensed-Phase Quantum Mechanical Calculations.” Journal of Computational Chemistry 24 (16): 1999–2012. 10.1002/jcc.10349.

Emsley, P, B Lohkamp, W G Scott, and K Cowtan. 2010. “Features and Development of Coot.” Acta Crystallographica Section D 66 (4): 486–501. doi:10.1107/S0907444910007493.

Evans, Philip R. 2011. “An Introduction to Data Reduction: Space-Group Determination, Scaling and Intensity Statistics.” Acta Crystallographica Section D: Biological Crystallography 67 (4): 282–92. 10.1107/S090744491003982X.

Faig, Margarita, Mario A. Bianchet, Paul Talalay, Shiuan Chen, Shannon Winski, David Ross, and L. Mario Amzel. 2000. “Structures of Recombinant Human and Mouse NAD(P)H:Quinone Oxidoreductases: Species Comparison and Structural Changes with Substrate Binding and Release.” Proceedings of the National Academy of Sciences of the United States of America 97 (7): 3177–82. 10.1073/pnas.97.7.3177.

Faig, Margarita, Mario A. Bianchet, Shannon Winski, Robert Hargreaves, Christopher J. Moody, Anna R. Hudnott, David Ross, and L. Mario Amzel. 2001. “Structure-Based Development of Anticancer Drugs: Complexes of NAD(P)H:Quinone Oxidoreductase 1 with Chemotherapeutic Quinones.” Structure 9 (8): 659–67. 10.1016/S0969-2126(01)00636-0.

Forneris, Federico, B Tom Burnley, and Piet Gros. 2014. “Ensemble Refinement Shows Conformational Flexibility in Crystal Structures of Human Complement Factor D.” Acta Crystallographica Section D 70 (3): 733–43. 10.1107/S1399004713032549.

Fraser, James S., Henry Van Den Bedem, Avi J. Samelson, P. Therese Lang, James M. Holton, Nathaniel Echols, and Tom Alber. 2011. “Accessing Protein Conformational Ensembles Using Room-Temperature X-Ray Crystallography.” Proceedings of the National Academy of Sciences of the United States of America 108 (39): 16247–52. 10.1073/pnas.1111325108.

Frauenfelder, Hans, Stephen G Sligar, and Peter G Wolynes. 1991. “The Energy Landscapes and Motions of Proteins.” Science 254 (5038): 1598–1603. 10.1126/science.1749933.

Frisch, M.J., G.W. Trucks, H.B. Schlegel, G.E. Scuseria, M.A. Robb, J.R. Cheeseman, G. Scalmani, et al. 2009. “Gaussian 09, Revision A.1.”

Gati, Cornelius, Gleb Bourenkov, Marco Klinge, Dirk Rehders, Francesco Stellato, Dominik Oberthür, Oleksandr Yefanov, et al. 2014. “Serial Crystallography on *in Vivo* Grown Microcrystals Using Synchrotron Radiation.” IUCrJ 1 (2): 87–94. 10.1107/S2052252513033939.

Gevorkov, Yaroslav, Oleksandr Yefanov, Anton Barty, Thomas A. White, Valerio Mariani, Wolfgang Brehm, Aleksandra Tolstikova, Rolf Rainer Grigat, and Henry N. Chapman. 2019. “XGANDALF - Extended Gradient Descent Algorithm for Lattice Finding.” Acta Crystallographica Section A: Foundations and Advances 75: 694–704. 10.1107/S2053273319010593.

Grieco, Alice, Miguel A. Ruiz-Fresneda, Atanasio Gómez-Mulas, Juan Luis Pacheco-García, Isabel Quereda-Moraleda, Angel L. Pey, and Jose M. Martin-Garcia. 2023. “ Structural Dynamics at the Active Site of the Cancer-associated Flavoenzyme NQO1 Probed by Chemical Modification with PMSF.” FEBS Letters, 1–12. 10.1002/1873-3468.14738.

Hess, Berk. 2008. “P-LINCS: A Parallel Linear Constraint Solver for Molecular Simulation.” Journal of Chemical Theory and Computation 4 (1): 116–22. 10.1021/ct700200b.

Hilser, Vincent J. 2010. “An Ensemble View of Allostery.” Science 327 (5966): 653–54. 10.1126/science.1186121.

Kavanaugh, Jeffrey S, Paul H Rogers, and Arthur Arnone. 2005. “Crystallographic Evidence for a New Ensemble of Ligand-Induced Allosteric Transitions in Hemoglobin: The T-to-THigh Quaternary Transitions,.” Biochemistry 44 (16): 6101–21. 10.1021/bi047813a.

Kieffer, J, N Coquelle, G Santoni, S Basu, S Debionne, A Homs, and D De Sanctis. 2022. “Real-Time Pre-Processing of Serial Crystallography.” Acta Crystallographica Section A 78 (a2): e263. 10.1107/S2053273322094530.

Krissinel, Evgeny, and Kim Henrick. 2005. “Detection of Protein Assemblies in Crystals BT - Computational Life Sciences.” In, edited by Michael R. Berthold, Robert C Glen, Kay Diederichs, Oliver Kohlbacher, and Ingrid Fischer, 163–74. Berlin, Heidelberg: Springer Berlin Heidelberg.

L. Pey, Angel, Clare F. Megarity, Encarnación Medina-Carmona, and David J. Timson. 2016. “Natural Small Molecules as Stabilizers and Activators of Cancer-Associated NQO1 Polymorphisms.” Current Drug Targets 17 (13): 1506–14. 10.2174/1389450117666160101121610.

Li, Rongbao, Mario A. Bianchet, Paul Talalay, and L. Mario Amzel. 1995. “The Three-Dimensional Structure of NAD(P)H:Quinone Reductase a Flavoprotein Involved in Cancer Chemoprotection and Chemotherapy: Mechanism of the Two-Electron Reduction.” Proceedings of the National Academy of Sciences of the United States of America 92 (19): 8846–50. 10.1073/pnas.92.19.8846.

Lienhart, Wolf-Dieter, Emilia Strandback, Venugopal Gudipati, Karin Koch, Alexandra Binter, Michael K Uhl, David M Rantasa, et al. 2017. “Catalytic Competence, Structure and Stability of the Cancer-Associated R139W Variant of the Human NAD(P)H:Quinone Oxidoreductase 1 (NQO1).” The FEBS Journal 284 (8):1233–45. 10.1111/febs.14051.

Lienhart, Wolf Dieter, Venugopal Gudipati, Michael K. Uhl, Alexandra Binter, Sergio A. Pulido, Robert Saf, Klaus Zangger, Karl Gruber, and Peter Macheroux. 2014. “Collapse of the Native Structure Caused by a Single Amino Acid Exchange in Human NAD(P)H:Quinone Oxidoreductase.” FEBS Journal 281 (20): 4691–4704. 10.1111/febs.12975.

Lu, Tian, and Feiwu Chen. 2012. “Multiwfn: A Multifunctional Wavefunction Analyzer.” Journal of Computational Chemistry 33 (5): 580–92. 10.1002/jcc.22885.

Maestre-Reyna, Manuel, Cheng-Han Yang, Eriko Nango, Wei-Cheng Huang, Eka Putra Gusti Ngurah Putu, Wen-Jin Wu, Po-Hsun Wang, et al. 2022. “Serial Crystallography Captures Dynamic Control of Sequential Electron and Proton Transfer Events in a Flavoenzyme.” Nature Chemistry 14 (6): 677–85. 10.1038/s41557-022-00922-3.

McGibbon, Robert T., Kyle A. Beauchamp, Matthew P. Harrigan, Christoph Klein, Jason M. Swails, Carlos X. Hernández, Christian R. Schwantes, Lee-Ping Wang, Thomas J. Lane, and Vijay S. Pande. 2015. “MDTraj: A Modern Open Library for the Analysis of Molecular Dynamics Trajectories.” Biophysical Journal 109 (8): 1528–32. 10.1016/j.bpj.2015.08.015.

Medina-Carmona, Encarnación, Julian E. Fuchs, Jose A. Gavira, Noel Mesa-Torres, Jose L. Neira, Eduardo Salido, Rogelio Palomino-Morales, Miguel Burgos, David J. Timson, and Angel L. Pey. 2017. “Enhanced Vulnerability of Human Proteins towards Disease-Associated Inactivation through Divergent Evolution.” Human Molecular Genetics 26 (18): 3531–44. 10.1093/hmg/ddx238.

Meng, Elaine C, Thomas D Goddard, Eric F Pettersen, Greg S Couch, Zach J Pearson, John H Morris, and Thomas E Ferrin. 2023. “UCSF ChimeraX: Tools for Structure Building and Analysis.” Protein Science 32 (11): e4792. 10.1002/pro.4792.

Motlagh, Hesam N., James O. Wrabl, Jing Li, and Vincent J. Hilser. 2014. “The Ensemble Nature of Allostery.” Nature 508 (7496): 331–39. 10.1038/nature13001.

Mozzanica, A, M Andrä, R Barten, A Bergamaschi, S Chiriotti, M Brückner, R Dinapoli, et al. 2018. “The JUNGFRAU Detector for Applications at Synchrotron Light Sources and XFELs.” Synchrotron Radiation News 31 (6): 16–20. 10.1080/08940886.2018.1528429.

Murshudov, Garib N., Pavol Skubák, Andrey A. Lebedev, Navraj S. Pannu, Roberto A. Steiner, Robert A. Nicholls, Martyn D. Winn, Fei Long, and Alexei A. Vagin. 2011. “REFMAC5 for the Refinement of Macromolecular Crystal Structures.” Acta Crystallographica Section D: Biological Crystallography 67 (4): 355–67. 10.1107/S0907444911001314.

Olmos, J.L., S. Pandey, J.M. Martin-Garcia, G. Calvey, A. Katz, J. Knoska, C. Kupitz, et al. 2018. “Enzyme Intermediates Captured ‘on the Fly’ by Mix-and-Inject Serial Crystallography.” BMC Biology 16 (1). 10.1186/s12915-018-0524-5.

Olsson, Mats H M, Chresten R Søndergaard, Michal Rostkowski, and Jan H Jensen. 2011. “PROPKA3: Consistent Treatment of Internal and Surface Residues in Empirical PKa Predictions.” Journal of Chemical Theory and Computation 7 (2): 525–37. 10.1021/ct100578z.

Pacheco-Garcia, Juan Luis, Ernesto Anoz-Carbonell, Pavla Vankova, Adithi Kannan, Rogelio Palomino-Morales, Noel Mesa-Torres, Eduardo Salido, et al. 2021. “Structural Basis of the Pleiotropic and Specific Phenotypic Consequences of Missense Mutations in the Multifunctional NAD(P)H:Quinone Oxidoreductase 1 and Their Pharmacological Rescue.” Redox Biology 46 (May): 102112. 10.1016/j.redox.2021.102112.

Pacheco-garcia, Juan Luis, Dmitry S. Loginov, Ernesto Anoz-carbonell, Pavla Vankova, Rogelio Palomino-morales, Eduardo Salido, Petr Man, Milagros Medina, Athi N. Naganathan, and Angel L. Pey. 2022. “Allosteric Communication in the Multifunctional and Redox NQO1 Protein Studied by Cavity-Making Mutations.” Antioxidants 11 (6). 10.3390/antiox11061110.

Pandey, Pankaj, Bharathi Avula, Ikhlas A. Khan, Shabana I. Khan, Victor J. Navarro, Robert J. Doerksen, and Amar G. Chittiboyina. 2020. “Potential Modulation of Human NAD[P]H-Quinone Oxidoreductase 1 (NQO1) by EGCG and Its Metabolites - A Systematic Computational Study.” Chemical Research in Toxicology 33 (11): 2749–64. 10.1021/acs.chemrestox.9b00450.

Pey, Angel L. 2018. “Biophysical and Functional Perturbation Analyses at Cancer-Associated P187 and K240 Sites of the Multifunctional NADP(H):Quinone Oxidoreductase 1.” International Journal of Biological Macromolecules 118: 1912–23. 10.1016/j.ijbiomac.2018.07.051.

Pey, Angel L., Clare F. Megarity, and David J. Timson. 2014. “FAD Binding Overcomes Defects in Activity and Stability Displayed by Cancer-Associated Variants of Human NQO1.” Biochimica et Biophysica Acta - Molecular Basis of Disease 1842 (11): 2163–73. 10.1016/j.bbadis.2014.08.011.

Pidugu, Lakshmi Swarna Mukhi, J. C. Emmanuel Mbimba, Muqeet Ahmad, Edwin Pozharski, Edward A. Sausville, Ashkan Emadi, and Eric A. Toth. 2016. “A Direct Interaction between NQO1 and a Chemotherapeutic Dimeric Naphthoquinone.” BMC Structural Biology 16 (1): 1–10. 10.1186/s12900-016-0052-x.

Powell, Harold R. 1999. “The Rossmann Fourier Autoindexing Algorithm in {\it MOSFLM}.” Acta Crystallographica Section D 55 (10): 1690–95. 10.1107/S0907444999009506.

Raimondi, Pantaleo, Chamseddine Benabderrahmane, Paul Berkvens, Jean Claude Biasci, Pawel Borowiec, Jean-Francois Bouteille, Thierry Brochard, et al. 2023. “The Extremely Brilliant Source Storage Ring of the European Synchrotron Radiation Facility.” Communications Physics 6 (1): 82. 10.1038/s42005-023-01195-z.

Ross, David, and David Siegel. 2017. “Functions of NQO1 in Cellular Protection and CoQ10 Metabolism and Its Potential Role as a Redox Sensitive Molecular Switch.” Frontiers in Physiology 8 (AUG): 1–10. 10.3389/fphys.2017.00595.

Schmidt, Marius. 2020. “Reaction Initiation in Enzyme Crystals by Diffusion of Substrate.” Crystals 10 (2): 1–15. 10.3390/cryst10020116.

Skelly, Jane V., Mark R. Sanderson, David A. Suter, Ulrich Baumann, Martin A. Read, David S.J. Gregory, Matthew Bennett, Stephen M. Hobbs, and Stephen Neidle. 1999. “Crystal Structure of Human DT-Diaphorase: A Model for Interaction with the Cytotoxic Prodrug 5-(Aziridin-1-Yl)-2,4-Dinitrobenzamide (CB1954).” Journal of Medicinal Chemistry 42 (21): 4325–30. 10.1021/jm991060m.

Sousa da Silva, Alan W, and Wim F Vranken. 2012. “ACPYPE - AnteChamber PYthon Parser InterfacE.” BMC Research Notes 5 (1): 367. 10.1186/1756-0500-5-367.

Stagno, Jason R, Yuba R Bhandari, Chelsie E Conrad, Yu Liu, and Yun-Xing Wang. 2017. “Real-Time Crystallographic Studies of the Adenine Riboswitch Using an X-Ray Free-Electron Laser.” The FEBS Journal 284 (20): 3374–80. 10.1111/febs.14110.

Strandback, Emilia, Wolf-Dieter Lienhart, Altijana Hromic-Jahjefendic, Benjamin Bourgeois, Anja Högler, Daniel Waltenstorfer, Andreas Winkler, et al. 2020. “A Small Molecule Chaperone Rescues the Stability and Activity of a Cancer-Associated Variant of NAD(P)H:Quinone Oxidoreductase 1 in Vitro.” FEBS Letters 594 (3): 424–38. 10.1002/1873-3468.13636.

Tian, Wei, Chang Chen, Xue Lei, Jieling Zhao, and Jie Liang. 2018. “CASTp 3.0: Computed Atlas of Surface Topography of Proteins.” Nucleic Acids Research 46 (W1): W363–67. 10.1093/nar/gky473.

Tyka, Michael D., Daniel A. Keedy, Ingemar André, Frank Dimaio, Yifan Song, David C. Richardson, Jane S. Richardson, and David Baker. 2011. “Alternate States of Proteins Revealed by Detailed Energy Landscape Mapping.” Journal of Molecular Biology 405 (2): 607–18. 10.1016/j.jmb.2010.11.008.

Vagin, A, and A Teplyakov. 1997. “MOLREP: An Automated Program for Molecular Replacement.” Journal of Applied Crystallography 30 (6): 1022–25. doi:10.1107/S0021889897006766.

Wang, Junmei, Wei Wang, Peter A Kollman, and David A Case. 2006. “Automatic Atom Type and Bond Type Perception in Molecular Mechanical Calculations.” Journal of Molecular Graphics and Modelling 25 (2): 247–60. 10.1016/j.jmgm.2005.12.005.

White, Thomas A. 2019. “Processing Serial Crystallography Data with CrystFEL: A Step-by-Step Guide.” Acta Crystallographica Section D: Structural Biology 75: 219–33. 10.1107/S205979831801238X.

White, Thomas A, Richard A Kirian, Andrew V Martin, Andrew Aquila, Karol Nass, Anton Barty, and Henry N Chapman. 2012. “*CrystFEL* : A Software Suite for Snapshot Serial Crystallography.” Journal of Applied Crystallography 45 (2): 335–41. 10.1107/S0021889812002312.

Winn, Martyn D., Charles C. Ballard, Kevin D. Cowtan, Eleanor J. Dodson, Paul Emsley, Phil R. Evans, Ronan M. Keegan, et al. 2011. “Overview of the CCP4 Suite and Current Developments.” Acta Crystallographica Section D: Biological Crystallography 67 (4): 235–42. 10.1107/S0907444910045749.

Winski, S. L., M. Faig, M. A. Bianchet, D. Siegel, E. Swann, K. Fung, M. W. Duncan, C. J. Moody, L. M. Amzel, and D. Ross. 2001. “Characterization of a Mechanism-Based Inhibitor of NAD(P)H:Quinone Oxidoreductase 1 by Biochemical, x-Ray Crystallographic, and Mass Spectrometric Approaches.” Biochemistry 40 (50): 15135–42. 10.1021/bi011324i.

Yefanov, Oleksandr, Valerio Mariani, Cornelius Gati, Thomas A White, Henry N Chapman, and Anton Barty. 2015. “Accurate Determination of Segmented X-Ray Detector Geometry.” Opt. Express 23 (22): 28459–70. 10.1364/OE.23.028459.

