## Supplemental Information for "Structural dynamics and functional cooperativity of human NQO1 by ambient temperature serial crystallography and simulations"

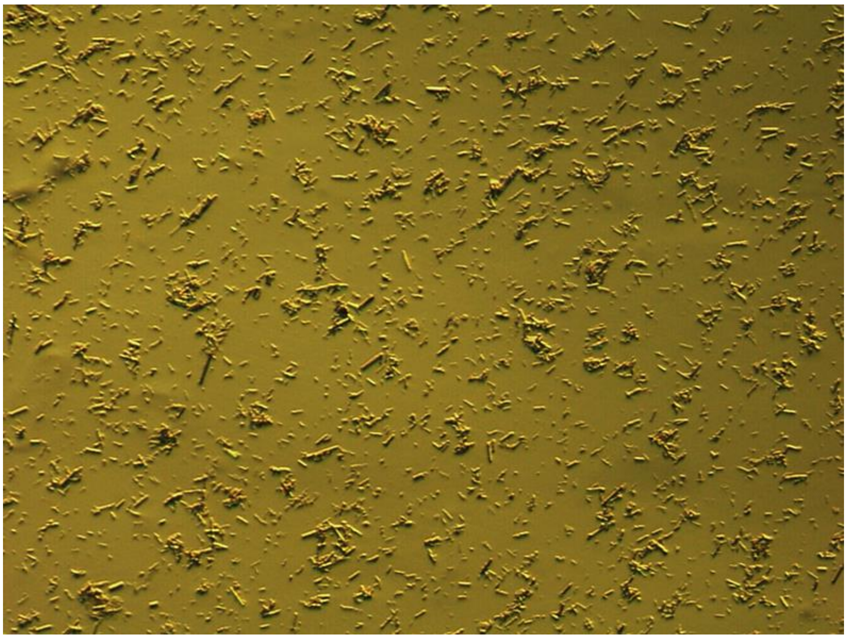

**Figure S1.** Microcrystals of hNQO1. Crystals of 10-30 μm in size were grown by the batch method as previously described (Doppler et al. 2023) in 0.1 M Tris pH 8.5, 0.2 M sodium acetate, 20 % PEG 3350.

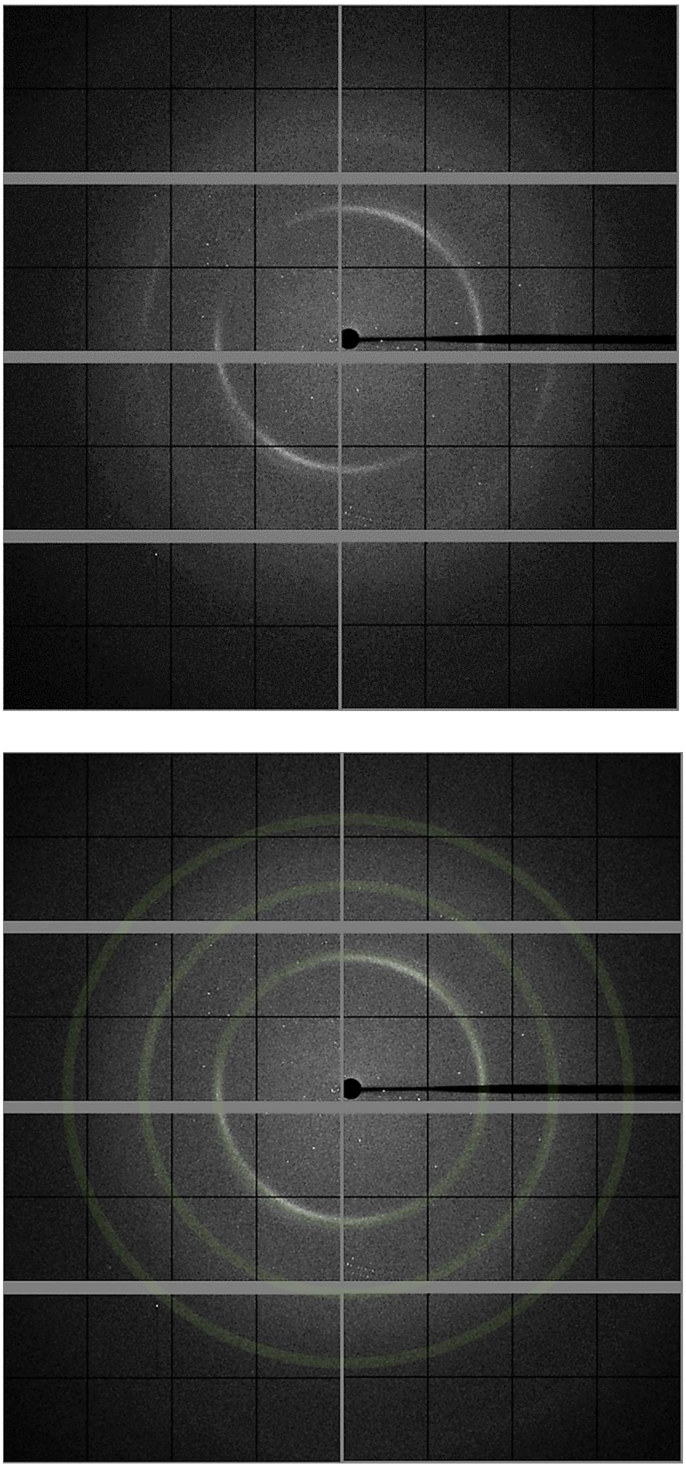

**Figure S2. Representative diffraction pattern of hNQO1 with NADH.** Crystals were seen to diffract up to ~2.5 Å resolution. Using the same diffraction pattern, the three background rings caused by the mylar film have been highlighted in transparent green on the bottom panel.

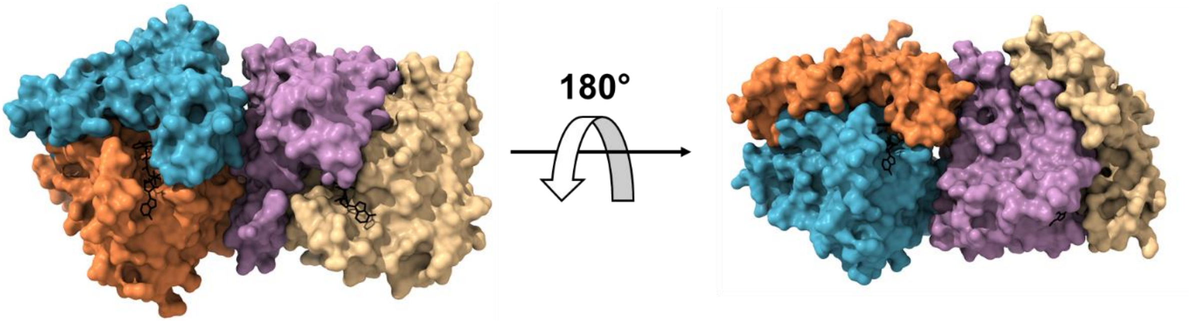


**Figure S3. Room temperature structure of the free hNQO1 protein obtained at ID29 beamline. A)** The two homodimers (chains A:B and C:D) of hNQO1 found in the ASU are depicted in surface representation. The individual monomers are highlighted in blue (chain A), orange (chain B), light magenta (chain C), and pale-yellow (chain D). The FAD cofactors are shown as black sticks.

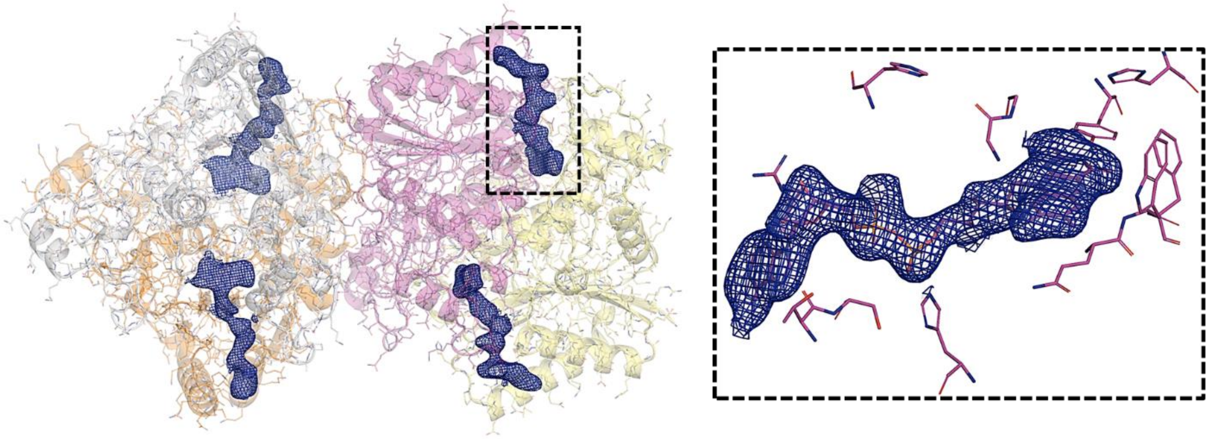

**Figure S4. Electron density maps of the free hNQO1 structure.** Cartoon and stick representation of the two homodimers of hNQO1 (left) found in the asymmetric unit. Electron density maps $2mF_o$-$DF_c$ contoured at 1 σ of the FAD are shown as blue meshes. The right panel shows a closer view of the catalytic site highlighted with the dashed box in the left panel. The FAD and the residues in the catalytic sites are shown as sticks.

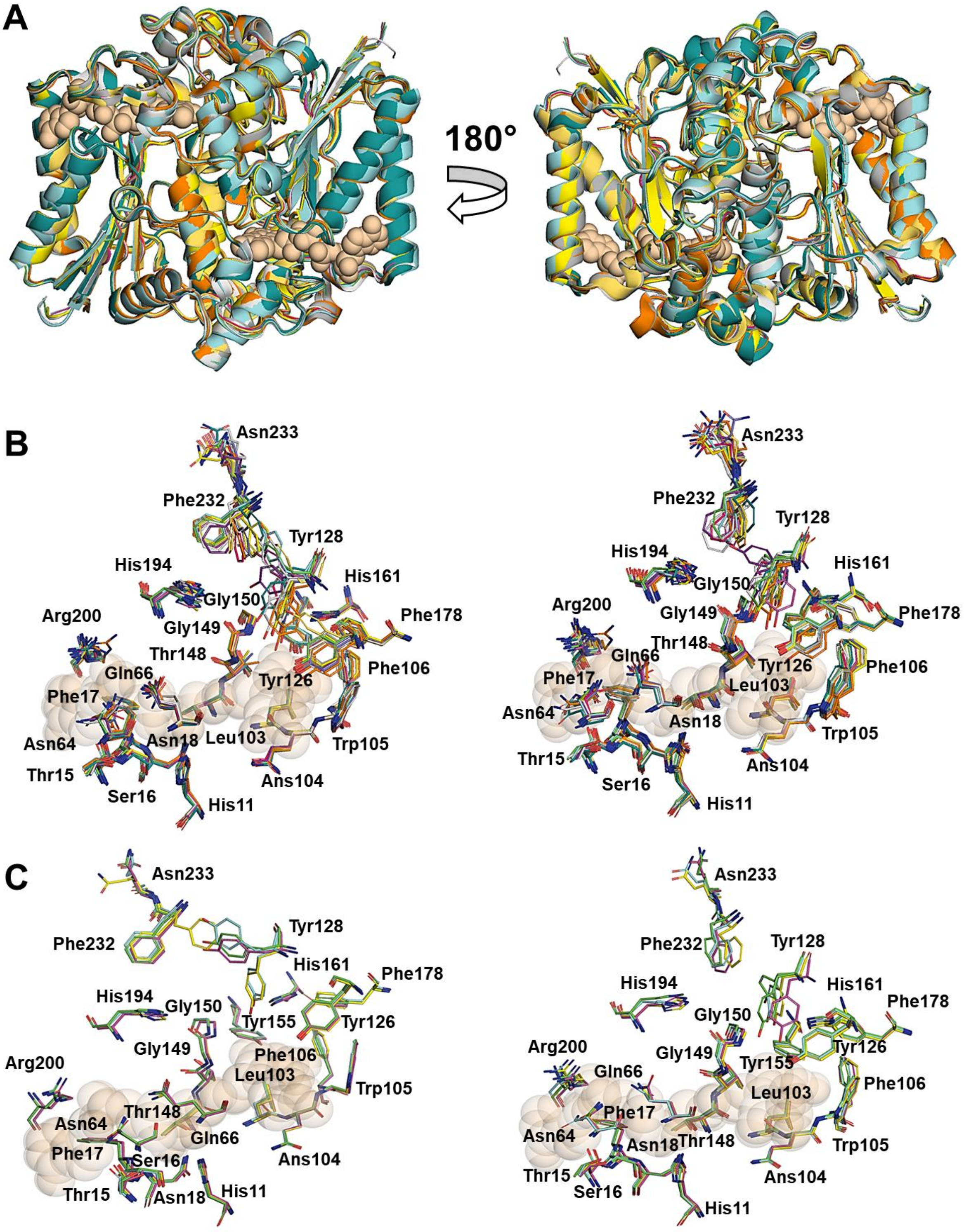


**Figure S5. Structural comparison of the hNQO1-NAD$^+$/H structure with other related hNQO1 structures. A)** Superposition of the NQO1-NAD$^+$/H structure with hNQO1 structures both unliganded (free hNQO1 (this study), PDB 8C9J (Doppler et al. 2023), and PDB 1D4A (Faig et al. 2000)), and in complex with NADH (computational model (this study )), the inhibitor PMSF (8OK0 (Grieco et al. 2023), the natural substrate duroquinone (PDB 1DXO (Faig et al. 2000)), various inhibitor molecules such as dicoumarol (PDB 5FUQ (Medina-Carmona et al. 2017)) and cibracron blue (PDB 4CF6 (Lienhart et al. 2014)), as well as a prodrug molecule (PDBs 1GG5 (Faig et al. 2001)). All protein molecules are represented as ribbons. All ligands from complex structures have been removed for clarity and only a FAD molecule is shown as pale-orange spheres representation to indicate where the catalytic site is located. **B)** Structural differences observed in the catalytic sites of all hNQO1 structures shown in A). All ligands in the complex structures have been removed for clarity. In all cases, the residues of the catalytic site are shown in stick representation and just an FAD molecule is shown in pale-orange spheres for clarity. **C)** Structural differences observed in the catalytic sites of the hNQO1 crystal structures reported in this study. In all cases, the residues of the catalytic site are shown in stick representation and just an FAD molecule is shown in pale-orange spheres for clarity.

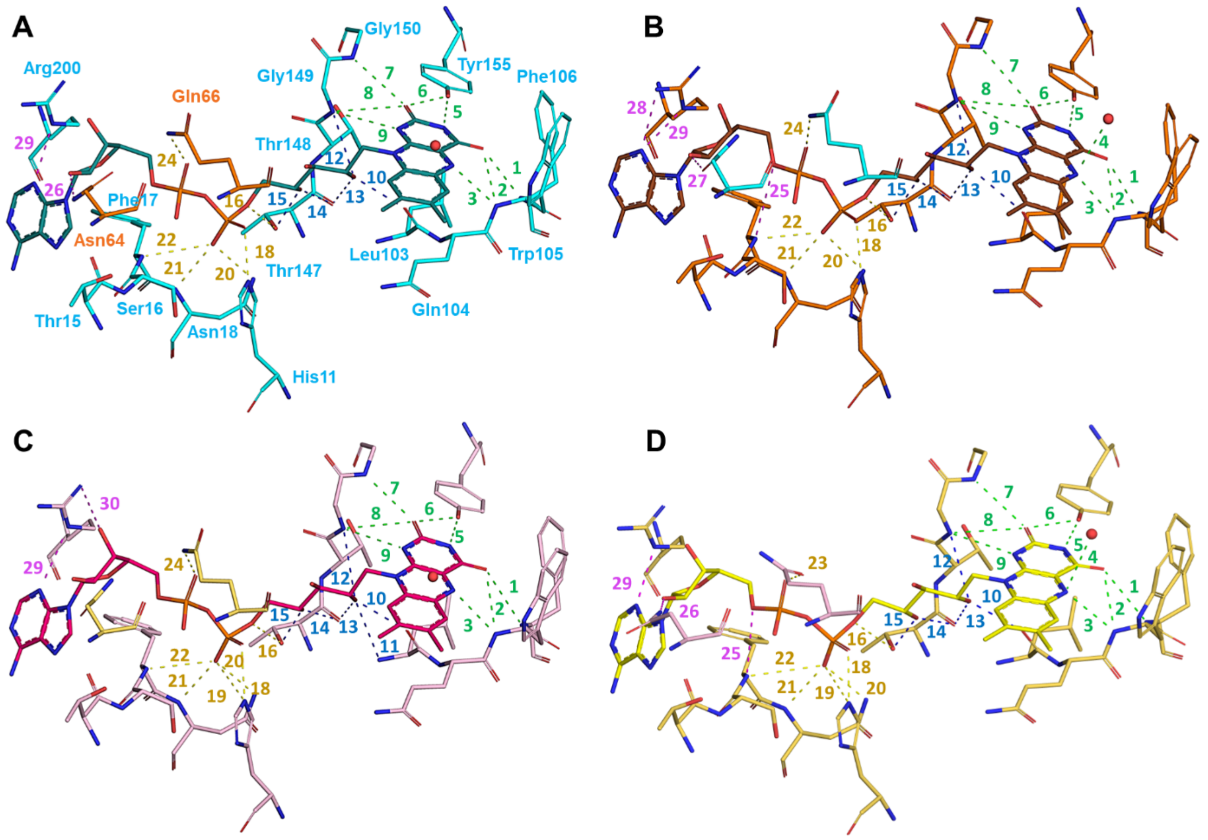


**Figure S6. Hydrogen bond interactions at the FAD binding site in the two homodimers of the free hNQO1 structure.** Stick representation of the FADs and all residues involved in the formation of the FAD binding site in one of the homodimers (**A)** and **B)** for chains A (cyan) and B (orange)), and in the other one (**C)** and **D)** for chains C (pink) and D (pale yellow)). Only residues of panel A) have been labeled for clarity. The color code of the protein residues is the same as that shown in Figure S3. All hydrogen bond interactions established between the FAD molecules and the protein residues have been labeled and are shown as dashes in the same color code as in Figure 5. The water molecule is represented as red sphere.

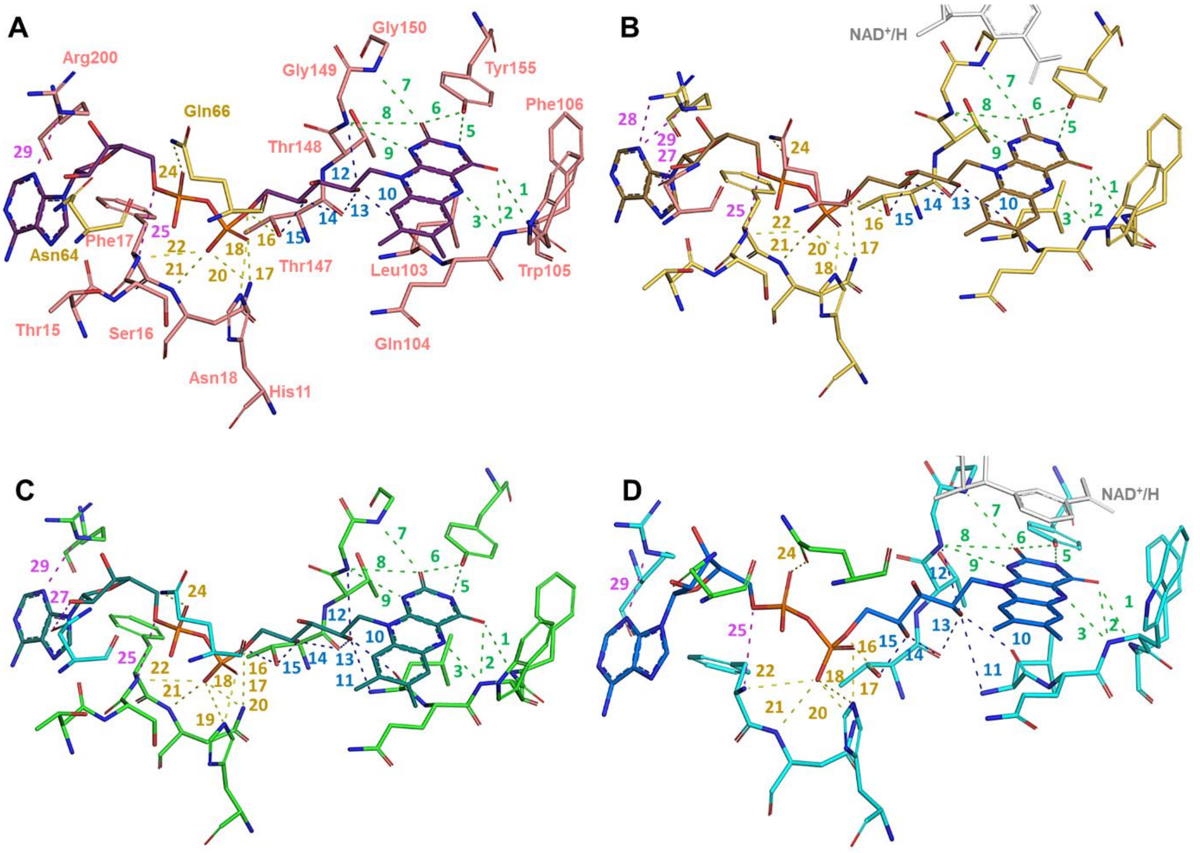


**Figure S7. Hydrogen bond interactions at the FAD binding site in the two homodimers of the structure of the complex hNQO1-NAD$^+$/H.** Stick representation of the FADs and all residues involved in the formation of the FAD binding site in one of the homodimers (**A)** and **B)** for chains A (salmon) and B (yellow)) and in the other one (**C)** and **D)** for chains C (green) and D (cyan)). Only residues of panel A) have been labeled for clarity. The color code of the protein residues is the same as that shown in Figure 2. All hydrogen bond interactions established between the FAD molecules and the protein residues have been labeled and are shown as dashes in the same color code as in Figure 5. The water molecule is represented as red sphere. The two NAD$^+$/H molecules are shown as gray sticks.

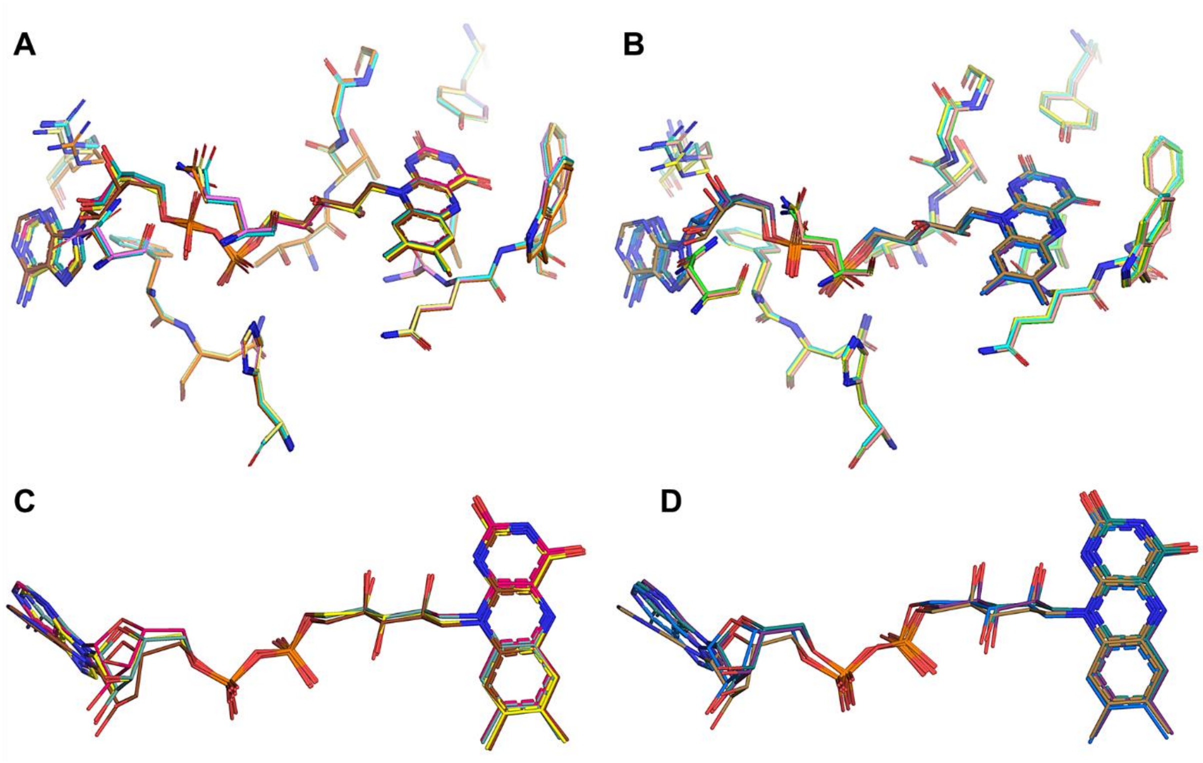


**Figure S8. FAD binding sites comparison of the free hNQO1 and hNQO1-NAD⁺/H structures. A)** Stick representation of the FAD molecules and the residues that comprise their binding sites of the two homodimers of the ASU in free hNQO1. **B)** Stick representation of the FAD molecules and the residues that comprise their binding sites of the two homodimers of the ASU in the complex hNQO1-NAD⁺/H. In both cases, the color code of the protein residues is the same as that in Figures S3 for the free hNQO1 and Figure 2 for the hNQO1-NAD⁺/H. The NAD⁺/H molecules have been removed from the complex for clarity. **C)** and **D)** Top view of the FADs represented in A) and B), respectively.

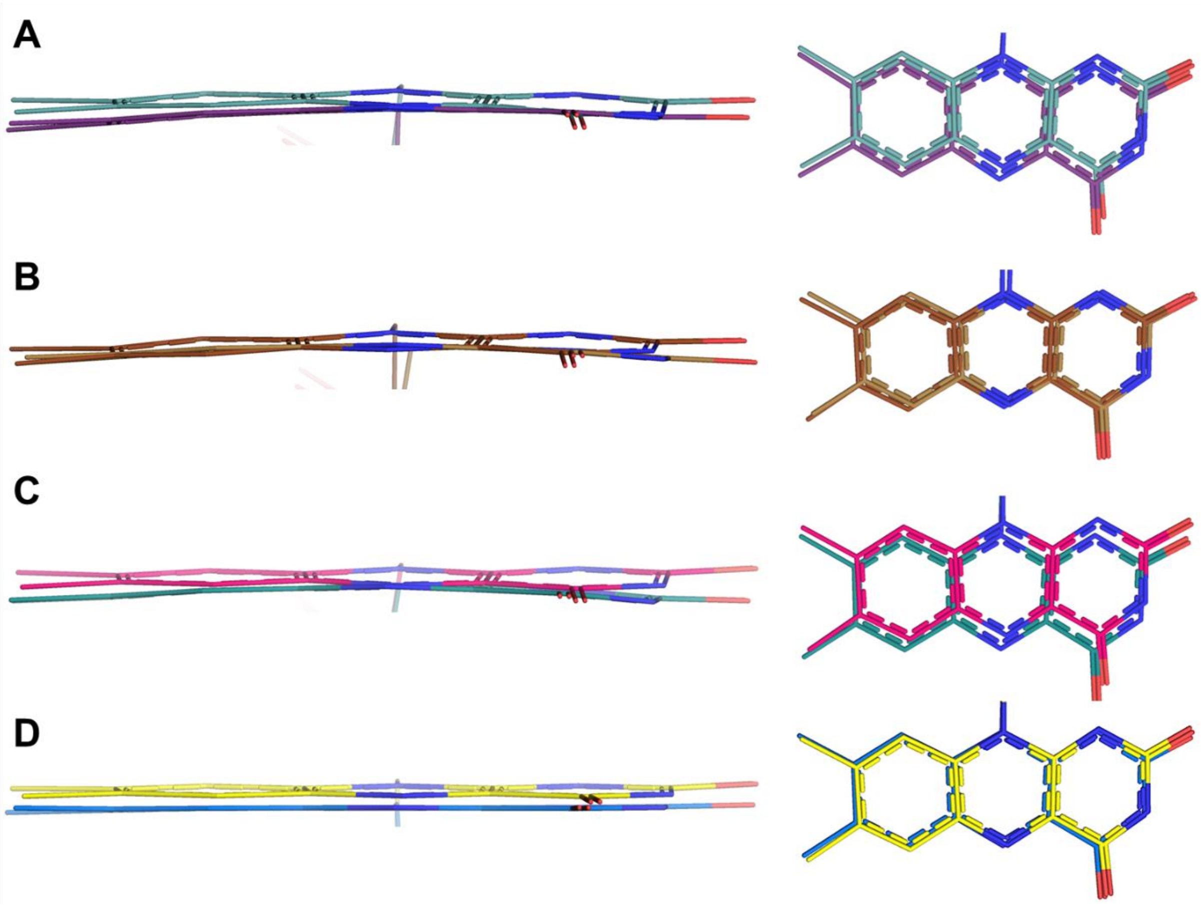


**Figure S9. Conformational changes in the isoalloxazine ring of the FADs. A)** Superposition of the isoalloxazine rings of chains A of the free hNQO1 (cyan) and the complex (violet). **B)** Superposition of the isoalloxazine rings of chains B of the free hNQO1 (dark yellow) and the complex (brown). **C)** Superposition of the isoalloxazine rings of chains C of the free hNQO1 (pink) and the complex (green). **D)** Superposition of the isoalloxazine rings of chains D of the free hNQO1 (yellow) and the complex (blue).

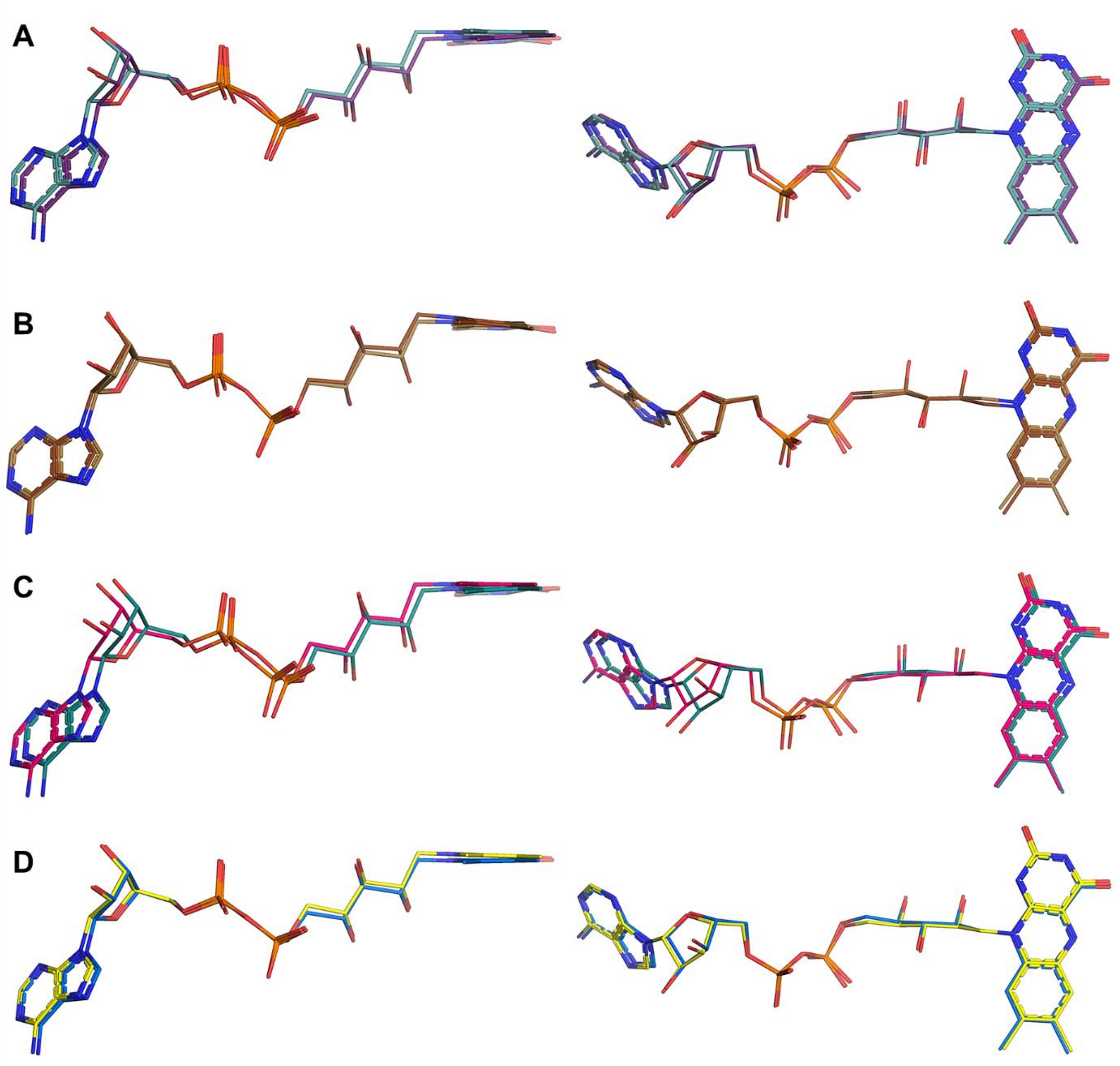


**Figure S10. Conformational changes in the FADs upon binding of NAD⁺/H. A)** Top (left) and side (right) views of the superposition of the FADs of chains A of the free hNQO1 (cyan) and the complex (violet). **B)** Top (left) and side (right) views of the superposition of the FADs of chains B of the free hNQO1 (dark yellow) and the complex (brown). **C)** Top (left) and side (right) views of the superposition of the FADs of chains C of the free hNQO1 (pink) and the complex (green). **D)** Top (left) and side (right) views of the superposition of the FADs of chains D of the free hNQO1 (yellow) and the complex (blue).

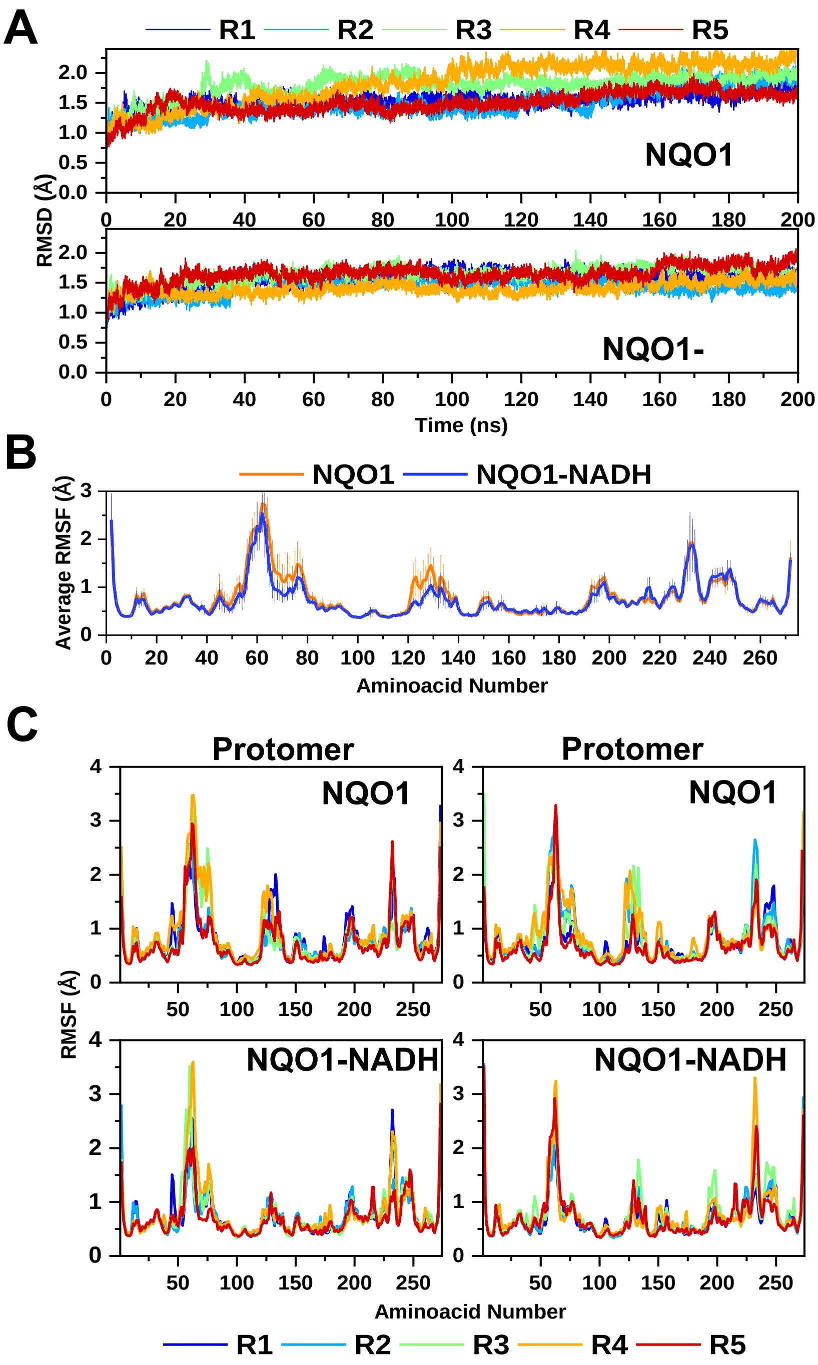


**Figure S11. Overall features of the 200 ns MD simulations for homodimers of hNQO1 and hNQO1-NADH models. A)** RMSD (Cα) of the five replicates (R1 to R5) throughout the MD simulation when overlapped to their corresponding starting models for the homodimers of hNQO1 (top) and the complex hNQO1-NADH (bottom). **B)** Average root mean square fluctuation (RMSF) along the amino acid sequence of hNQO1 and hNQO1-NADH homodimers. Average values correspond to the mean of the five MD replicates in the 20-200 ns range and include at each position the residues of the two protomers of the homodimer. Average values are shown in bold lines, while their corresponding standard deviation for each residue are shown in thick vertical lanes. **C)** Root mean square fluctuation (RMSF) along the amino acid sequence of hNQO1 and hNQO1-NADH by protomer within the homodimer along the 20-200 ns range of the MD simulation. Data corresponding to the same replicate have the same color for the two protomers of the homodimer in the case of the hNQO1 (top panels) and hNQO1-NADH complex (low panels).

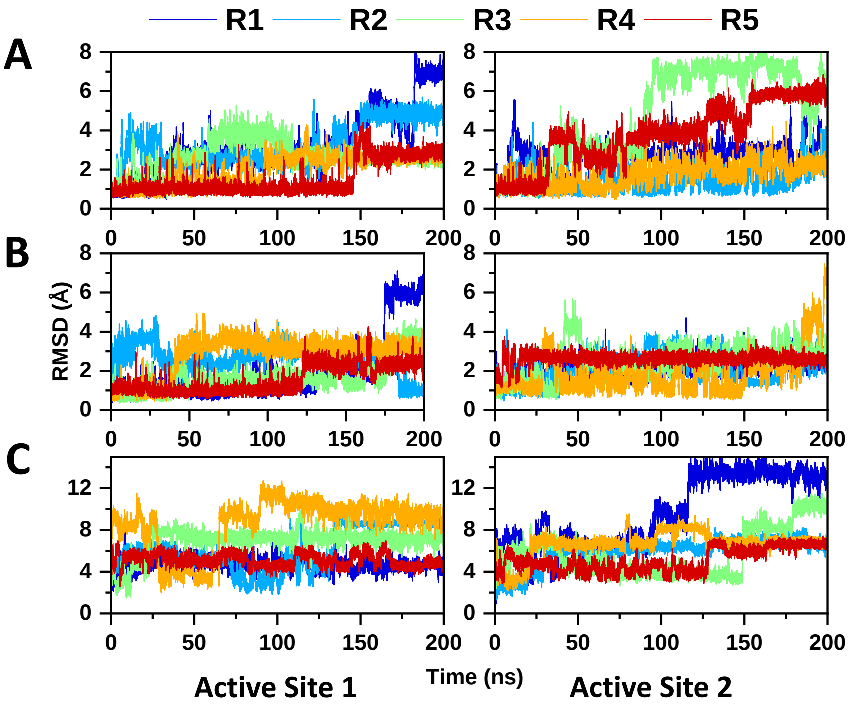


**Figure S12. Evolution of RMSD of the FAD and NADH ligands at active site 1 and active site 2 throughout the MD simulation. A)** FAD in free hNQO1 homodimer. **B)** FAD and **C)** NADH in the NADH bound hNQO1 homodimer. Data corresponding to the same replicate have the same colour for the ligands at the two active sites of the homodimer, and for both ligands in the case of the hNQO1-NADH complex.

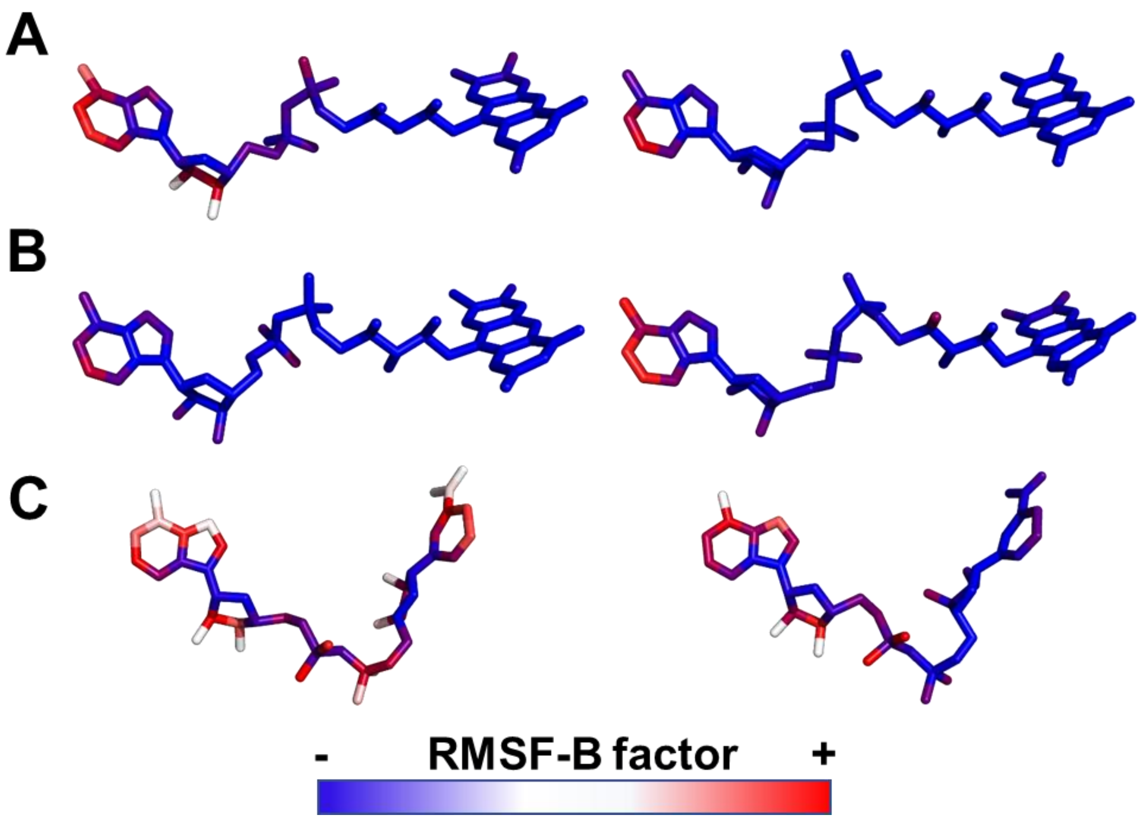


**Figure S13. B-factor equivalent from Root Mean Square Fluctuation (RMSF-B factor, 20-220 nm2) for atom of the ligands at active site 1 and active site 2**. **A)** FAD for replicate 2 (R2, taken as representative) of the MD simulation in free hNQO1 homodimer. **B)** FAD and **C)** NADH for replicate 4 (R4, taken as representative) of the MD simulation in the NADH bound hNQO1 homodimer.

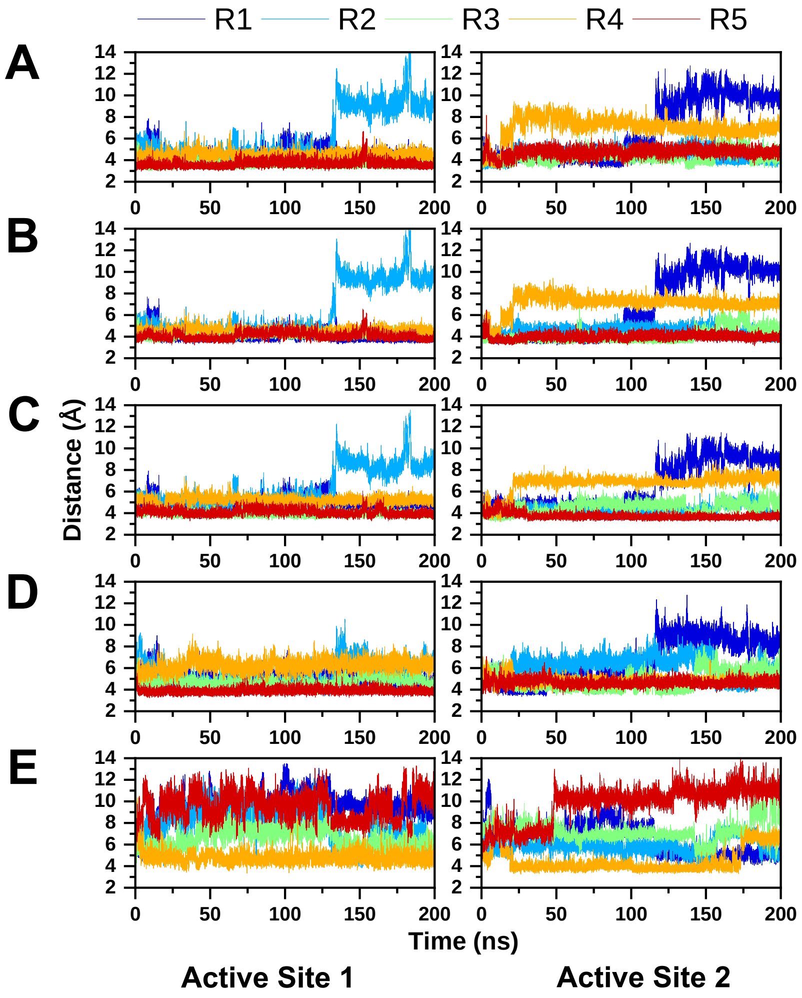


**Figure S14. Evolution of key distances between the nicotinamide of FAD, the isoalloxazine ring of FAD and the side chains of Tyr126 and Tyr128 through the 200 ns MD simulation of the hNQO1-NADH complex. A)** N5 of the FAD isoalloxazine to C4 of the nicotinamide. **B)** Centre of the nicotinamide ring of NADH to the center of the pyrazine ring of the FAD isoalloxazine. **C)** Centre of the nicotinamide ring of NADH to the center of the pyrimidine ring of the FAD isoalloxazine. **D)** Centre of the nicotinamide ring of NADH to the center of the aromatic ring of Tyr126. **E)** Centre of the nicotinamide ring of NADH to the center of the aromatic ring of Tyr128. Data corresponding to the same replicate have the same color at the two active sites of the homodimer and for all the shown distances.

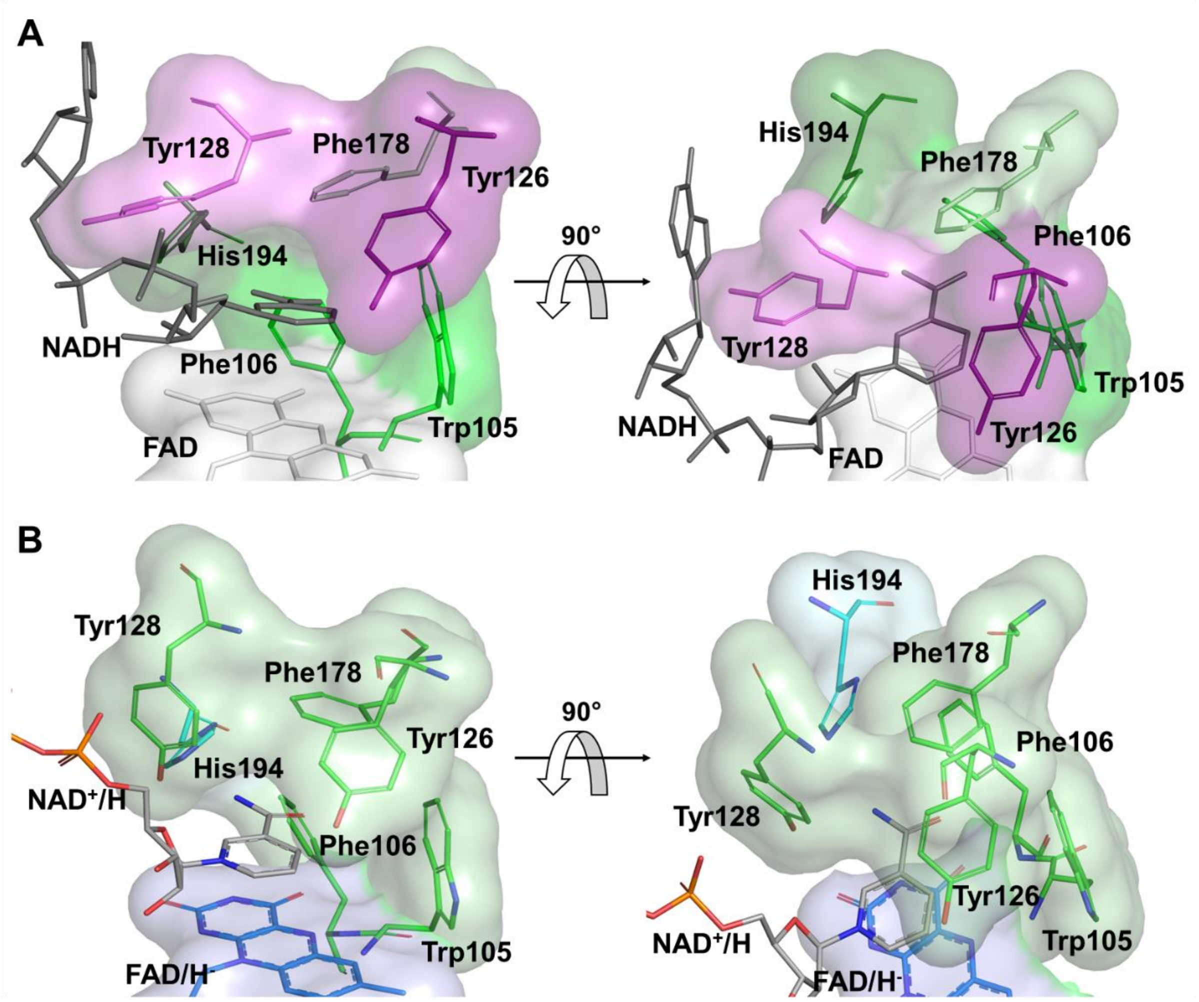


**Figure S15. Stacking of the nicotinamide moiety of the NADH at the catalytic site.** **A)** Front (left) and top (right) view of the stacking of the nicotinamide moiety at the catalytic site 1 in the simulations (frame at 30ns of MD simulations, replicate 2). All residues that form the nicotinamide binding pocket and the NADH molecule, are shown as sticks in the same color code as in the simulations image (Figure 7). **B)** Front (left) and top (right) view of the stacking of the nicotinamide moiety of the $NAD^+/H_D$ molecule at the catalytic site in the crystal structure of the complex (this study). All residues that form the nicotinamide binding pocket and the NADH molecules, are shown as sticks.

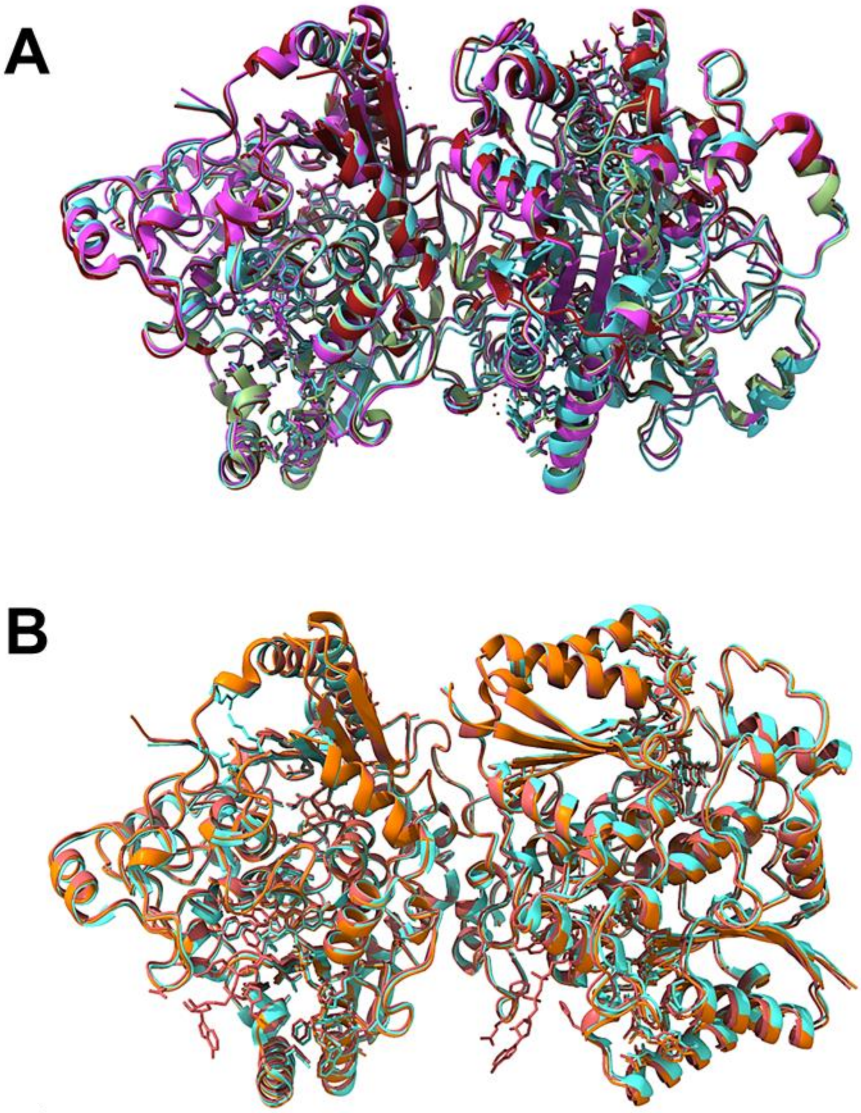


**Figure S17. Comparison of the packing in the two homodimers of hNQO1 from different structures obtained from large crystals. A)** Superposition of the two homodimers observed in the large crystals of the free hNQO1 in PDB 5A4K (light green) (W.-D. Lienhart et al. 2017) and PDB 5EA2 (red) (Pidugu et al. 2016), the hNQO1 in complex with BPPSA PDB 6FY4 (pink) (Strandback et al. 2020), and the two homodimers in the large crystals of PDB 8OK0 (light blue) (Grieco et al. 2023). **B)** Superposition of the two homodimers in the microcrystals of the free hNQO1 (orange) and hNQO1-NAD$^{+}$/H (red), this study, and PDB 8C9J (light blue) (Doppler et al. 2023).

**Table S1:** Surface and buried areas of the free hNQO1 and complex hNQO1-$NAD^+$/H structures.

| Structure | Chains | Surface area ($Å^2$) | Buried area ($Å^2$) |
|---|---|---|---|
| Free hNQO1 | A:B | 23,400 | 5,480 |
| | C:D | 23,400 | 5,490 |
| hNQO1-$NAD^+$/H | A:B | 22,400 | 5,420 |
| | C:D | 22,300 | 5,390 |

**Table S2:** Intermolecular hydrogen bond interactions established at the interface residues in the homodimer composed by A and B of the free hNQO1.

| # | Residue (A) /atom | Residue (B) /atom | Distance (Å) |
|---|---|---|---|
| 1 | Glu13 [OE2] | Arg52 [NE] | 3.6 |
| 2 | Glu13 [OE1] | Arg52 [NH2] | 3.0 |
| 3 | Glu13 | Arg52 [NE] | 3.5 |
| 4 | Tyr42 [OH] | Ile50 [O] | 2.9 |
| 5 | Ile50 [O] | Tyr42 [OH] | 3.5 |
| 6 | Arg52 [NH2] | Glu13 [OE1] | 2.6 |
| 7 | Trp105 [NE1] | Phe116 [O] | 3.6 |
| 8 | Gln105 [O] | Lys113 [NZ] | 2.6 |
| 9 | Val108 [O] | Lys113 [NZ] | 3.0 |
| 10 | Ala110 [N] | Glu117 [OE1] | 3.2 |
| 11 | Ala110 [N] | Glu117 [OE2] | 3.5 |
| 12 | Lys113 [NZ] | Val108 [O] | 2.8 |
| 13 | Lys113 [NZ] | Gln104 [O] | 2.7 |
| 14 | Phe116 [O] | Trp105 [NE1] | 3.5 |
| 15 | Glu117 [OE2] | Ala110 [N] | 3.5 |
| 16 | Glu117 [OE1] | Ala110 [N] | 3.3 |
| 17 | Tyr132 [OH] | His161 [ND1] | 2.9 |
| 18 | Ser153 [OG] | Gly235 [O] | 2.6 |
| 19 | Gln158 [O] | Met238 [N] | 2.9 |
| 20 | Gly159 [O] | His257 [NE2] | 2.9 |
| 21 | Ile160 [N] | Phe236 [O] | 3.2 |
| 22 | His161 [ND1] | Tyr132 [OH] | 2.9 |
| 23 | Asp163 [OD2] | His258 [NE2] | 2.7 |
| 24 | Asp163 [N] | Gly256 [O] | 3.2 |
| 25 | Gly235 [O] | Ser153 [OG] | 2.7 |
| 26 | Phe236 [O] | Ile160 [N] | 3.2 |
| 27 | Met238 [N] | Gln158 [O] | 3.0 |
| 28 | Gly256 [O] | Asp163 [N] | 3.1 |
| 29 | His257 [NE2] | Gly159 [O] | 2.8 |
| 30 | His258 [NE2] | Asp163 [OD2] | 2.7 |
| 31 | His258 [NE2] | Asp163 [OD2] | 2.7 |
| 32 | Gly260 [O] | Ser262 [OG] | 2.7 |
| 33 | Ser262 [OG] | Gly260 [O] | 2.7 |

**Table S3:** Intermolecular hydrogen bond interactions established at the interface residues in the homodimer composed by chains C and D of the free hNQO1.

| # | Residue (C) /atom | Residue (D) /atom | Distance (Å) |
|---|---|---|---|
| 1 | Glu13 [OE1] | Arg52 [NH2] | 3.1 |
| 2 | Thr15 [OG1] | Asn64 [OD1] | 3.6 |
| 3 | Tyr42 [OH] | Ile50 [O] | 3.3 |
| 4 | Ile50 [O] | Tyr42 [OH] | 3.4 |
| 5 | Arg52 [NH2] | Glu13 [OE1] | 3.4 |
| 6 | Gln104 [O] | Lys113 [NZ] | 2.8 |
| 7 | Trp105 [NE1] | Phe116 [O] | 3.6 |
| 8 | Val108 [O] | Lys113 [NZ] | 2.7 |
| 9 | Ala110 [N] | Glu117 [OE1] | 3.3 |
| 10 | Ala110 [N] | Glu117 [OE2] | 3.4 |
| 11 | Lys113 [NZ] | Gln104 [O] | 2.7 |
| 12 | Lys113 [NZ] | Val108 [O] | 2.8 |
| 13 | Phe116 [O] | Trp105 [NE1] | 3.5 |
| 14 | Glu117 [OE1] | Ala110 [N] | 3.2 |
| 15 | Glu117 [OE2] | Ala110 [N] | 3.3 |
| 16 | Tyr132 [OH] | His161 [ND1] | 2.9 |
| 17 | Ser153 [OG] | Gly235 [O] | 2.9 |
| 18 | Gln158 [O] | Met238 [O] | 3.0 |
| 19 | Gly159 [O] | His257 [NE2] | 2.9 |
| 20 | Ile160 [N] | Phe236 [O] | 3.1 |
| 21 | His161 [ND1] | Tyr132 [OH] | 3.0 |
| 22 | Asp163 [N] | Gly256 [O] | 3.2 |
| 23 | Phe236 [O] | Ile160 [N] | 3.0 |
| 24 | Met238 [N] | Gln158 [O] | 2.8 |
| 25 | Gly256 [O] | Asp163 [N] | 3.2 |
| 26 | His257 [NE2] | Asp163 [OD2] | 2.7 |
| 27 | His257 [NE2] | Gly159 [O] | 2.9 |
| 28 | Ser262 [OG] | Gly260 [O] | 2.8 |

**Table S4:** Intermolecular hydrogen bond interactions established at the interface residues in the homodimer composed by A and B of the complex hNQO1-NAD$^+$/H.

| # | Residue (A) /atom | Residue (B) /atom | Distance (Å) |
|---|---|---|---|
| 1 | Glu13 [OE2] | Arg52 [NE] | 3.4 |
| 2 | Glu13 [OE1] | Arg52 [NH2] | 2.6 |
| 3 | Arg52 [NH2] | Glu13 [OE1] | 2.7 |
| 4 | Gln104 [O] | Lys113 [NZ] | 2.7 |
| 5 | Trp105 [NE1] | Phe116 [0] | 3.5 |
| 6 | Val108 [O] | Lys113 [NZ] | 2.7 |
| 7 | Lys113 [NZ] | Val108 [O] | 2.8 |
| 8 | Lys113 [NZ] | Gln104 [O] | 2.7 |
| 9 | Glu117 [OE1] | Ala110 [N] | 3.2 |
| 10 | Glu117 [OE2] | Ala110 [N] | 3.4 |
| 11 | Tyr132 [OH] | His161 [ND1] | 2.8 |
| 12 | Aer153 [OG] | Gly235 [O] | 2.6 |
| 13 | Gln158 [O] | Met238 [N] | 2.8 |
| 14 | Gly159 [O] | His257 [NE2] | 2.8 |
| 15 | Ile160 [N] | Phe236 [O] | 3.0 |
| 16 | His161 [ND1] | Tyr132 [OH] | 2.7 |
| 17 | Asp163 [N] | Gly256 [O] | 3.1 |
| 18 | Asp163 [OD2] | His258 [NE2] | 2.7 |
| 19 | Gly235 [O] | Ser153 [OG] | 2.6 |
| 20 | Phe236 [O] | Ile160 [N] | 3.1 |
| 21 | Met238 [N] | Gln158 [O] | 3.0 |
| 22 | Gly256 [O] | Asp163 [N] | 3.1 |
| 23 | His257 [NE2] | Gly159 [O] | 3.1 |
| 24 | His258 [NE2] | Asp163 [OD2] | 2.7 |
| 25 | Gly260 [O] | Ser262 [OG] | 2.6 |
| 26 | Ser262 [OG] | Gly260 [O] | 2.5 |

**Table S5:** Intermolecular hydrogen bond interactions established at the interface residues in the homodimer composed by chains C and D of the complex hNQO1-NAD$^+$/H.

| # | Residue (C) /atom | Residue (D) /atom | Distance (Å) |
|---|---|---|---|
| 1 | Glu13 [OE1] | Arg52 [NH2] | 2.6 |
| 2 | Arg52 [NH2] | Glu13 [OE1] | 2.8 |
| 3 | Gln104 [O] | Lys113 [NZ] | 2.7 |
| 4 | Gln104 [OE1] | Glu117 [OE2] | 3.0 |
| 5 | Val108 [O] | Lys113 [NZ] | 2.8 |
| 6 | Ala110 [N] | Glu117 [OE1] | 3.2 |
| 7 | Ala110 [N] | Glu117 [OE2] | 3.2 |
| 8 | Lys113 [NZ] | Gln104 [O] | 2.6 |
| 9 | Lys113 [NZ] | Val108 [O] | 2.9 |
| 10 | Glu117 [OE1] | Ala110 [N] | 3.2 |
| 11 | Glu117 [OE2] | Ala110 [N] | 3.2 |
| 12 | Tyr132 [OH] | His161 [ND1] | 2.7 |
| 13 | Ser153 [OG] | Gly235 [O] | 2.8 |
| 14 | Gln158 [O] | Met238 [M] | 2.9 |
| 15 | Gly159 [O] | His257 [NE2] | 2.7 |
| 16 | Ile160 [N] | Phe236 [O] | 3.1 |
| 17 | His161 [ND1] | Tyr132 [OH] | 2.7 |
| 18 | Asp163 [N] | Gly256 [O] | 3.0 |
| 19 | Asp163 [OD2] | His258 [NE2] | 2.6 |
| 20 | Gly235 [O] | Ser153 [OG] | 2.6 |
| 21 | Phe236 [O] | Ile160 [N] | 2.9 |
| 22 | Met238 [M] | Gln158 [O] | 2.8 |
| 23 | Gly256 [O] | Asp163 [N] | 3.1 |
| 24 | His257 [NE2] | Gly159 [O] | 2.8 |
| 25 | His258 [NE2] | Asp163 [OD2] | 2.7 |
| 26 | Gly260 [O] | Ser262 [OG] | 2.9 |
| 27 | Ser262 [OG] | Gly260 [O] | 2.8 |

**Table S6:** Intramolecular hydrogen bond interactions established at the interface by residues of chain A of the free hNQO1.

| # | Residue /atom | Residue /atom | Distance (Å) |
|---|---|---|---|
| 1 | Glu13 [OE2] | Ser12 [OG] | 2.6 |
| 2 | Asn64 [O] | Asn64 [ND2] | 3.2 |
| 3 | Gln104 [N] | Gly107 [O] | 3.1 |
| 4 | Gly107 [N] | Gln104 [O] | 3.3 |
| 5 | Ala110 [O] | Gly114 [N] | 3.0 |
| 6 | Ala110 [O] | Lys113 [N] | 3.3 |
| 7 | Lys113 [N] | Pro109 [O] | 3.0 |
| 8 | Phe116 [N] | Lys113 [O] | 3.4 |
| 9 | Gly117 [OE2] | Lys113 [NZ] | 3.0 |
| 10 | Glu117 [N] | Lys113 [O] | 3.2 |
| 11 | Phe120 [N] | Phe116[O] | 3.0 |
| 12 | Tyr132 [N] | Phe178 [O] | 3.0 |
| 13 | Ser153 [OG] | Ser151 [OG] | 3.2 |
| 14 | Met154 [N] | Ser151 [O] | 3.1 |
| 15 | Ser156 [N] | Ser153 [O] | 3.3 |
| 16 | Ser156 [OG] | Ser153 [O] | 2.7 |
| 17 | Gln158 [N] | Gln158 [E1] | 3.1 |
| 18 | His161[N] | Met154 [O] | 3.1 |
| 19 | Val166 [N] | Asp163 [O] | 3.3 |
| 20 | Trp169 [N] | Val166 [O] | 3.4 |
| 21 | Gly174 [N] | Trp169 [O] | 3.6 |
| 22 | Gly174 [N] | Pro170 [O] | 3.2 |
| 23 | Gly174 [N] | Trp169 [O] | 3.6 |
| 24 | Leu230 [N] | Phe228 [O] | 3.5 |
| 25 | Gln243 [NE2] | Met238 [O] | 3.5 |
| 26 | His258 [ND1] | Gly260 [N] | 3.2 |
| 27 | Lys261 [N] | His258 [O] | 2.9 |

**Table S7:** Intramolecular hydrogen bond interactions established at the interface by residues of chain B of the free hNQO1.

| # | Residue /atom | Residue /atom | Distance (Å) |
|---|---|---|---|
| 1 | Gln105 [N] | Gly107 [O] | 3.1 |
| 2 | Gly107 [N] | Gln104 [O] | 3.3 |
| 3 | Lys113 [N] | Pro109 [O] | 3.0 |
| 4 | Lys113 [N] | Ala110 [O] | 3.3 |
| 5 | Lys113 [NZ] | Glu117 [OE2] | 3.0 |
| 6 | Gly114 [N] | Ala110 [O] | 3.0 |
| 7 | Phe116 [N] | Lys113 [O] | 3.4 |
| 8 | Glu117 [N] | Lys113 [O] | 3.2 |
| 9 | Phe120 [N] | Phe116 [O] | 3.0 |
| 10 | Tyr132 [N] | Phe178 [O] | 3.0 |
| 11 | Ser153 [OG] | Ser151 [OG] | 3.2 |
| 12 | Met154 [N] | Ser151 [O] | 3.1 |
| 13 | Ser156 [N] | Ser153 [O] | 3.3 |
| 14 | Ser156 [OG] | Ser153 [O] | 2.7 |
| 15 | Gln158 [N] | Gln158 [OE1] | 3.1 |
| 16 | Gly159 [N] | Ser156 [O] | 3.1 |
| 17 | His161 [N] | Met154 [O] | 3.1 |
| 18 | Gly162 [N] | Gly159 [O] | 3.1 |
| 19 | Val166 [N] | Asp163 [O] | 3.3 |
| 20 | Trp169 [O] | Gly174 [N] | 3.6 |
| 21 | Trp169 [N] | Val166 [O] | 3.4 |
| 22 | Pro170 [O] | Gly174 [N] | 3.2 |
| 23 | Leu230 [N] | Phe228 [O] | 3.5 |
| 24 | Gln243 [NE2] | Met238 [O] | 3.4 |
| 25 | Gly260 [N] | His258 [ND1] | 3.2 |
| 26 | Lys261 [N] | His258 [O] | 2.9 |

**Table S8:** Intramolecular hydrogen bond interactions established at the interface by residues of chain C of the free hNQO1.

| # | Residue /atom | Residue /atom | Distance (Å) |
|---|---|---|---|
| 1 | Ser12 [OG] | Glu13 [OE2] | 2.6 |
| 2 | Gln104 [N] | Gly107 [O] | 3.1 |
| 3 | Gly107 [N] | Gln104 [O] | 3.3 |
| 4 | Lys113 [N] | Ala110 [O] | 3.3 |
| 5 | Lys113 [N] | Pro109 [O] | 3.0 |
| 6 | Lys113 [NZ] | Glu117 [OE2] | 3.0 |
| 7 | Gly114 [N] | Ala110 [O] | 3.0 |
| 8 | Phe116 [N] | Lys113 [O] | 3.4 |
| 9 | Glu117 [N] | Lys113 [O] | 3.2 |
| 10 | Phe120 [N] | Phe116 [O] | 3.0 |
| 11 | Tyr132 [N] | Phe178 [O] | 3.0 |
| 12 | Ser156 [N] | Ser153 [O] | 3.3 |
| 13 | Ser156 [OG] | Ser153 [O] | 2.7 |
| 14 | Gln158 [N] | Gln158 [OE1] | 3.1 |
| 15 | Gly159 [N] | Ser156 [O] | 3.1 |
| 16 | His161 [N] | Met154 [O] | 3.2 |
| 17 | Gly162 [N] | Gly159 [O] | 3.1 |
| 18 | Val166 [N] | Asp163 [O] | 3.3 |
| 19 | Trp169 [N] | Val166 [O] | 3.4 |
| 20 | Gly174 Ǹ] | Pro170 [O] | 3.2 |
| 21 | Gly174 Ǹ] | Trp169 [O] | 3.6 |
| 22 | Gln243 [NE2] | Met238 [O] | 3.4 |
| 23 | Lys261 [N] | His258 [O] | 2.9 |

**Table S9:** Intramolecular hydrogen bond interactions established at the interface by residues of chain D of the free hNQO1.

| # | Residue /atom | Residue /atom | Distance (Å) |
|---|---|---|---|
| 1 | Asn64 [ND2] | Asn64 [O] | 3.2 |
| 2 | Gln104 [N] | Gly107 [O] | 3.1 |
| 3 | Gly107 [N] | Gln104 [O] | 3.3 |
| 4 | Lys113 [NZ] | Glu117 [OE2] | 3.0 |
| 5 | Lys113 [N] | Ala110 [O] | 3.3 |
| 6 | Lys113 [N] | Pro109 [O] | 3.0 |
| 7 | Gly114 [N] | Ala110 [O] | 3.0 |
| 8 | Phe116 [N] | Lys113 [O] | 3.4 |
| 9 | Glu117 [N] | Lys113 [O] | 3.2 |
| 10 | Phe120 [N] | Phe116 [O] | 3.0 |
| 11 | Tyr132 [N] | Phe178 [O] | 3.0 |
| 12 | Ser153 [OG] | Ser151 [OG] | 3.1 |
| 13 | Met154 [N] | Ser151 [O] | 3.1 |
| 14 | Ser156 [OG] | Ser153 [O] | 2.7 |
| 15 | Ser156 [N] | Ser153 [O] | 3.3 |
| 16 | Gln158 [N] | Gln158 [OE1] | 3.1 |
| 17 | Gly159 [N] | Ser156 [O] | 3.1 |
| 18 | His161 [N] | Met154 [O] | 3.1 |
| 19 | Val166 [N] | Asp163 [O] | 3.3 |
| 20 | Trp169 [N] | Val166 [O] | 3.4 |
| 21 | Gly174 [N] | Trp169 [O] | 3.6 |
| 22 | Gly174 [N] | Pro170 [O] | 3.2 |
| 23 | Leu230 [N] | Phe228 [O] | 3.5 |
| 24 | Gln243 [NE2] | Met238 [O] | 3.4 |
| 25 | His258 [N] | His258 [ND1] | 3.3 |
| 26 | Lys261 [N] | His258 [O] | 2.9 |

**Table S10:** Intramolecular hydrogen bond interactions established at the interface by residues of chain A of the complex hNQO1-NAD+/H.

| # | Residue / atom | Residue / atom | Distance (Å) |
|---|---|---|---|
| 1 | Asn64 [O] | Asn64 [ND2] | 3.1 |
| 2 | Gln104 [N] | Gly107 [O] | 3.0 |
| 3 | Gln104 [O] | Gly107 [N] | 3.1 |
| 4 | Pro109 [O] | Lys113 [N] | 2.9 |
| 5 | Ala110 [O] | Lys113 [N] | 3.2 |
| 6 | Ala110 [O] | Gly114 [N] | 2.8 |
| 7 | Lys113 [O] | Phe116 [N] | 3.4 |
| 8 | Lys113 [O] | Glu117 [N] | 3.2 |
| 9 | Lys113 [NZ] | Glu117 [OE2] | 2.9 |
| 10 | Phe116 [O] | Phe120 [N] | 2.9 |
| 11 | Tyr132 [N] | Phe178 [O] | 3.0 |
| 12 | Ser151 [O] | Met154 [N] | 3.0 |
| 13 | Ser151 [OG] | Ser153 [OG] | 3.1 |
| 14 | Ser153 [O] | Ser156 [OG] | 2.7 |
| 15 | Ser153 [O] | Ser156 [N] | 3.3 |
| 16 | Met154 [O] | His161 [N] | 3.1 |
| 17 | Met154 [O] | Gly162 [N] | 3.4 |
| 18 | Gln158 [N] | Gln158 [OE1] | 3.1 |
| 19 | Gly159 [N] | Ser156 [O] | 2.9 |
| 20 | Gly162 [N] | Gly159 [O] | 3.1 |
| 21 | Asp163 [O] | Val166 [N] | 3.2 |
| 22 | Trp169 [N] | Val166 [O] | 3.5 |
| 23 | Trp169 [O] | Gly176 [N] | 3.5 |
| 24 | Pro170 [O] | Gly174 [N] | 3.0 |
| 25 | Phe178 [O] | Tyr132 [N] | 3.0 |
| 26 | Ser225 [O] | Phe228 [N] | 3.0 |
| 27 | Phe232 [O] | Gly235 [N] | 3.0 |
| 28 | Met238 [O] | Gln243 [NE2] | 3.1 |
| 29 | His258 [O] | Lys261 [N] | 3.0 |
| 30 | Gly260 [N] | His258 [ND1] | 3.1 |

**Table S11:** Intramolecular hydrogen bond interactions established at the interface by residues of chain B of the complex hNQO1-NAD$^+$/H.

| # | Residue / atom | Residue / atom | Distance (Å) |
|---|---|---|---|
| 1 | Gln104 [N] | Gly107 [O] | 3.0 |
| 2 | Gln104 [O] | Gly107 [N] | 3.1 |
| 3 | Pro109 [O] | Lys113 [N] | 2.9 |
| 4 | Ala110 [O] | Lys113 [N] | 3.2 |
| 5 | Ala110 [O] | Gly114 [N] | 2.8 |
| 6 | Lys113 [O] | Phe116 [N] | 3.4 |
| 7 | Lys113 [O] | Glu117 [N] | 3.2 |
| 8 | Lys113 [NZ] | Glu117 [OE2] | 2.9 |
| 9 | Gly114 [N] | Ala110 [O] | 2.8 |
| 10 | Phe116 [O] | Phe120 [N] | 2.9 |
| 11 | Tyr132 [N] | Phe178 [O] | 3.1 |
| 12 | Ser153 [O] | Ser156 [N] | 3.3 |
| 13 | Ser153 [O] | Ser156 [OG] | 2.7 |
| 14 | Met154 [O] | His161 [N] | 3.1 |
| 15 | Ser156 [N] | Ser153 [O] | 3.3 |
| 16 | Ser156 [O] | Gly159 [N] | 2.9 |
| 17 | Ser156 [OG] | Ser153 [O] | 2.7 |
| 18 | Gln158 [N] | Gln158 [OE1] | 3.1 |
| 19 | Val166 [N] | Asp165 [O] | 3.2 |
| 20 | Val166 [O] | Trp169 [N] | 3.5 |
| 21 | Trp169 [O] | Gly174 [N] | 3.5 |
| 22 | Pro170 [O] | Gly174 [N] | 3.0 |
| 23 | Ser225 [O] | Phe228 [N] | 3.0 |
| 24 | Phe232 [O] | Gly235 [N] | 3.0 |
| 25 | Met238 [O] | Gln243 [NE2] | 3.2 |
| 26 | His258 [O] | Lys261 [N] | 3.0 |
| 27 | His258 [ND1] | Gly260 [N] | 3.1 |

**Table S12:** Intramolecular hydrogen bond interactions established at the interface by residues of chain C of the complex hNQO1-NAD$^+$/H.

| # | Residue / atom | Residue / atom | Distance (Å) |
|---|---|---|---|
| 1 | Asn64 [N] | Asn64 [O] | 3.1 |
| 2 | Gln104 [N] | Gly107 [O] | 3.0 |
| 3 | Gln104 [O] | Gly107 [N] | 3.1 |
| 4 | Pro109 [O] | Lys113 [N] | 2.9 |
| 5 | Ala110 [O] | Lys113 [N] | 3.2 |
| 6 | Ala110 [O] | Gly114 [N] | 2.8 |
| 7 | Lys113 [NZ] | Glu117 [OE2] | 2.9 |
| 8 | Phe116 [N] | Lys113 [O] | 3.4 |
| 9 | Phe116 [O] | Phe120 [N] | 2.9 |
| 10 | Glu117 [N] | Lys113 [O] | 3.2 |
| 11 | Tyr132 [N] | Phe178 [O] | 3.0 |
| 12 | Ser225 [O] | Phe228 [N] | 3.0 |
| 13 | Phe232 [O] | Gly235 [N] | 3.0 |
| 14 | Met238 [O] | Gln243 [NE2] | 3.2 |
| 15 | His258 [O] | Lys261 [N] | 3.0 |
| 16 | His258 [ND1] | Gly260 [N] | 3.1 |

**Table S13:** Intramolecular hydrogen bond interactions established at the interface by residues of chain D of the complex hNQO1-NAD$^+$/H.

| # | Residue /atom | Residue /atom | Distance (Å) |
|---|---|---|---|
| 1 | Asn64 [N] | Asn64 [O] | 3.0 |
| 2 | Gln104 [N] | Gly107 [O] | 3.0 |
| 3 | Gln104 [O] | Gly107 [N] | 3.1 |
| 4 | Pro109 [O] | Lys113 [N] | 2.9 |
| 5 | Ala110 [O] | Lys113 [N] | 3.2 |
| 6 | Ala110 [O] | Gly114 [N] | 2.8 |
| 7 | Lys113 [NZ] | Glu117 [OE2] | 2.9 |
| 8 | Lys113 [O] | Phe116 [N] | 3.3 |
| 9 | Lys113 [O] | Glu117 [N] | 3.2 |
| 10 | Phe116 [O] | Phe120 [N] | 2.9 |
| 11 | Tyr132 [N] | Phe178 [O] | 3.0 |
| 12 | Ser151 [O] | Met154 [N] | 3.0 |
| 13 | Ser151 [OG] | Ser153 [OG] | 3.1 |
| 14 | Ser153 [O] | Ser156 [N] | 3.3 |
| 15 | Ser153 [O] | Ser156 [OG] | 2.7 |
| 16 | Met154 [O] | His161 [N] | 3.1 |
| 17 | Met154 [O] | Gly162 [N] | 3.4 |
| 18 | Ser156 [O] | Gly159 [N] | 2.9 |
| 19 | Gln158 [OE1] | Gln158 [N] | 3.1 |
| 20 | Gly159 [O] | Gly162 [N] | 3.1 |
| 21 | Asp163 [O] | Val166 [N] | 3.2 |
| 22 | Val166 [O] | Trp169 [N] | 3.5 |
| 23 | Trp169 [O] | Gly174 [N] | 3.5 |
| 24 | Gly174 [N] | Pro170 [O] | 3.0 |
| 25 | Ser225 [O] | Phe228 [N] | 3.0 |
| 26 | Met238 [O] | Gln243 [NE2] | 3.2 |
| 27 | His258 [O] | Lys261 [N] | 3.0 |
| 28 | His258 [ND1] | Gly260 [N] | 3.1 |

**Table S14.** Percentage of time that the hydrogen bond/polar contacts are maintained for the FAD atoms at each active site of the free hNQO1 homodimer for each replicate along the 200 ns MD simulation.

| # | FAD atom | Residue [atom] | Active Site 1 | | | | | | Active Site 2 | | | | |
|---|---|---|---|---|---|---|---|---|---|---|---|---|---|
| | | | R1 | R2 | R3 | R4 | R5 | | R1 | R2 | R3 | R4 | R5 |
| 1 | O4 | Phe106 [N] | 60 | 97 | 92 | 98 | 97 | | 78 | 71 | 97 | 96 | 86 |
| 2 | O4 | Trp105 [N] | 68 | 84 | 90 | 85 | 86 | | 95 | 77 | 86 | 77 | 89 |
| 3 | N5 | Trp105 [N] | 65 | 100 | 98 | 100 | 100 | | 98 | 98 | 99 | 99 | 100 |
| 5 | N3 | Try155 [OH] | 54 | 98 | 81 | 97 | 93 | | 72 | 85 | 92 | 87 | 9 |
| 6 | O2 | Try155 [OH] | 37 | 53 | 66 | 22 | 40 | | 49 | 44 | 28 | 22 | 36 |
| 7 | O2 | Gly150 [N] | 51 | 92 | 77 | 95 | 82 | | 83 | 93 | 95 | 47 | 9 |
| 8 | O2 | Gly149 [N] | 77 | 95 | 80 | 90 | 81 | | 96 | 72 | 9 | 48 | 84 |
| 9 | N1 | Gly149 [N] | 35 | 61 | 52 | 68 | 47 | | 35 | 41 | 75 | 33 | 50 |
| 10 | O2´ | Gly149 [N] | 67 | 46 | 57 | 57 | 65 | | 52 | 70 | 44 | 41 | 54 |
| 11 | O2´ | Leu103 [O] | 35 | 91 | 86 | 94 | 79 | | 27 | 89 | 65 | 94 | 92 |
| 12 | O2´ | Leu103 [N] | 9.3 | 18 | 18 | 21 | 11 | | 5.1 | 13 | 5.1 | 36 | 22 |
| 13 | O2´ | Thr147 [O] | 44 | 35 | 28 | 30 | 34 | | 11.0 | 17 | 20 | 31 | 30 |
| 14 | O4´ | Thr147 [O] | 38 | 79 | 85 | 85 | 74 | | 22 | 76 | 15 | 93 | 86 |
| 15 | O4´ | Thr147 [OG1] | 57 | 100 | 100 | 100 | 99 | | 26 | 90 | 17 | 100 | 100 |
| 16 | O5´ | Thr147 [OG1] | 36 | 66 | 76 | 80 | 50 | | 59 | 66 | 80 | 72 | 60 |
| 17 | O5´ | Asn18 [ND2] | 43 | 80 | 71 | 67 | 29 | | 77 | 36 | 63 | 45 | 43 |
| 18 | O2P | His11 [NE2] | 99 | 100 | 100 | 100 | 100 | | 100 | 99 | 100 | 99 | 99 |
| 19 | O1P | His11 [NE2] | 63 | 42 | 48 | 59 | 64 | | 28 | 66 | 46 | 77 | 51 |
| 20 | O1P | Asn18 [ND2] | 100 | 100 | 100 | 100 | 100 | | 100 | 99 | 100 | 100 | 100 |
| 21 | O1P | Asn18 [N] | 98 | 98 | 99 | 99 | 98 | | 99 | 97 | 89 | 98 | 967 |
| 22 | O1P | Phe17 [N] | 97 | 99 | 99 | 98 | 95 | | 98 | 96 | 94 | 97 | 95 |
| 23 | O2A | Gln66 [OE1]* | 1.1 | 1.2 | 0.3 | 2.6 | 2.4 | | 3.8 | 0.2 | 0.1 | 0.6 | 4.5 |
| 24 | O2A | Gln66 [NE2]* | 50 | 35 | 16 | 19 | 44 | | 25 | 31 | 1.8 | 12 | 42 |
| 25 | O5B | Phe17 [N] | 25 | 40 | 15 | 29 | 10 | | 40 | 15 | 11 | 8.2 | 19 |
| 26 | O2B | Asn64 [ND2]* | 14 | 9.3 | 0.5 | 0.4 | 1.1 | | 0.9 | 0.7 | 1.3 | 0.2 | 22 |
| 27 | O2B | Asn64 [OD1]* | 1 | 9.6 | 0.7 | 0.2 | 1.6 | | 0.4 | 0.5 | 1.3 | 0.1 | 24 |
| 28 | N3A | Arg200 [NH2] | 0.8 | 2.7 | 1.3 | 0.5 | 0.5 | | 2.1 | 0.7 | 1.1 | 0.6 | 7.0 |
| 29 | N3A | Arg200 [NE] | 0.1 | 0.3 | 0.3 | 0.3 | 0.2 | | 2.8 | 0.1 | 0.1 | 0.3 | 0.8 |

*All of the FAD/protein interactions have been highlighted using the same color code and numbering as in Figure 5. Green for the isoalloxazine moiety, blue for the ribitol moiety, yellow for the phosphates moiety, and pink for the ribose and adenine moieties. The cut-off to consider a hydrogen bond interaction is 3.6 Å.*

**Most interactions shown for FADs at active sites 1 and 2 refer respectively to those with chains A and B, with the only exception of those labelled with * that respectively correspond to chains B and A in the free hNQO1 homodimer.*

**Table S15.** Percentage of time that the hydrogen bond/polar contacts are maintained for the FAD atoms at each active site of the complex hNQO1-NADH homodimer for each replicate along the 200 MD simulation.

| # | FAD atom | Residue [atom] | Active Site 1 | | | | | | Active Site 2 | | | | |
|---|---|---|---|---|---|---|---|---|---|---|---|---|---|
| | | | R1 | R2 | R3 | R4 | R5 | | R1 | R2 | R3 | R4 | R5 |
| 1 | O4 | Phe106 [N] | 91 | 95 | 81 | 98 | 85 | | 31 | 88 | 60 | 84 | 100 |
| 2 | O4 | Trp105 [N] | 76 | 87 | 92 | 91 | 93 | | 78 | 85 | 93 | 97 | 88 |
| 3 | N5 | Trp105 [N] | 100 | 100 | 98 | 100 | 99 | | 41 | 99 | 66 | 91 | 100 |
| 5 | N3 | Try155 [OH] | 96 | 83 | 78 | 94 | 95 | | 36 | 87 | 48 | 51 | 98 |
| 6 | O2 | Try155 [OH] | 20 | 72 | 84 | 62 | 65 | | 38 | 65 | 39 | 91 | 25 |
| 7 | O2 | Gly150 [N] | 98 | 77 | 99 | 90 | 98 | | 55 | 99 | 66 | 67 | 99 |
| 8 | O2 | Gly149 [N] | 88 | 97 | 86 | 99 | 91 | | 64 | 90 | 68 | 73 | 97 |
| 9 | N1 | Gly149 [N] | 47 | 53 | 70 | 73 | 49 | | 44 | 54 | 44 | 55 | 70 |
| 10 | O2´ | Gly149 [N] | 70 | 51 | 87 | 54 | 54 | | 68 | 75 | 76 | 62 | 50 |
| 11 | O2´ | Leu103 [O] | 87 | 62 | 96 | 87 | 86 | | 62 | 47 | 44 | 77 | 93 |
| 12 | O2´ | Leu103 [N] | 15 | 11 | 13 | 15 | 12 | | 35 | 4.7 | 9.2 | 23 | 15 |
| 13 | O2´ | Thr147 [O] | 30 | 40 | 44 | 34 | 32 | | 64 | 24 | 31 | 24 | 34 |
| 14 | O4´ | Thr147 [O] | 80 | 71 | 81 | 79 | 71 | | 17 | 17 | 17 | 66 | 83 |
| 15 | O4´ | Thr147 [OG1] | 100 | 99 | 87 | 100 | 100 | | 22 | 19 | 22 | 74 | 100 |
| 16 | O5´ | Thr147 [OG1] | 59 | 46 | 44 | 80 | 54 | | 23 | 59 | 54 | 69 | 86 |
| 17 | O5´ | Asn18 [ND2] | 43 | 62 | 10 | 81 | 46 | | 27 | 82 | 80 | 47 | 85 |
| 18 | O2P | His11 [NE2] | 99 | 100 | 100 | 100 | 100 | | 99 | 99 | 100 | 99 | 100 |
| 19 | O1P | His11 [NE2] | 63 | 63 | 37 | 40 | 51 | | 68 | 27 | 45 | 67 | 46 |
| 20 | O1P | Asn18 [ND2] | 100 | 99 | 100 | 100 | 100 | | 100 | 100 | 100 | 100 | 100 |
| 21 | O1P | Asn18 [N] | 99 | 99 | 99 | 99 | 99 | | 99 | 99 | 98 | 99 | 100 |
| 22 | O1P | Phe17 [N] | 97 | 99 | 98 | 99 | 98 | | 98 | 98 | 98 | 97 | 100 |
| 23 | O2A | Gln66 [OE1]* | 1.2 | 6.7 | 0.0 | 0.1 | 3.4 | | 0.0 | 2.2 | 0.4 | 0.4 | 1.6 |
| 24 | O2A | Gln66 [NE2]* | 37 | 54 | 25 | 9.7 | 45 | | 42 | 51 | 25 | 23 | 29 |
| 25 | O5B | Phe17 [N] | 17 | 32 | 1.3 | 2.2 | 18 | | 2.2 | 69 | 63 | 6.1 | 43 |
| 26 | O2B | Asn64 [ND2]* | 0.3 | 13 | 0.2 | 0.0 | 1.8 | | 0.6 | 1.6 | 1.8 | 0.5 | 2.2 |
| 27 | O2B | Asn64 [OD1]* | 1.0 | 17 | 0.1 | 0.0 | 1.8 | | 0.1 | 2.2 | 3.9 | 0.6 | 3.9 |
| 28 | N3A | Arg200 [NH2] | 0.1 | 0.7 | 0.4 | 24 | 0.2 | | 0.2 | 6.0 | 1.0 | 1.3 | 0.0 |
| 29 | N3A | Arg200 [NE] | 0.2 | 0.1 | 0.1 | 18 | 0.2 | | 0.2 | 5.0 | 0.3 | 0.2 | 0.0 |

*All of the FAD/protein interactions have been highlighted using the same color code and numbering as in Figure 5. Green for the isoalloxazine moiety, blue for the ribitol moiety, yellow for the phosphates moiety, and pink for the ribose and adenine moieties. The cut-off to consider a hydrogen bond interaction is 3.6 Å.*

**Most interactions shown for FADs at active sites 1 and 2 refer respectively to those with chains A and B, with the only exception of those labelled with * that respectively correspond to chains B and A in the hNQO1-NADH homodimer.*

**Table S16.** Percentage of polar contacts maintained for the NADH atoms and the protein at each active site of the complex hNQO1-NADH homodimer for each replicate along the 200 ns MD simulation.

| # | FAD atom | Residue [atom] | Active Site 1 | | | | | | Active Site 2 | | | | |
|---|---|---|---|---|---|---|---|---|---|---|---|---|---|
| | | | R1 | R2 | R3 | R4 | R5 | | R1 | R2 | R3 | R4 | R5 |
| 3 | N7N | His161 [NE2]* | 0.7 | 0.1 | 51 | 0.1 | 62 | | 0.2 | 12 | 0.3 | 2.8 | 0.1 |
| | N7N | Tyr126 [OH] | 1.1 | 26 | 0.0 | 0.0 | 2.2 | | 16 | 6.9 | 8.2 | 1.4 | 3.4 |
| 4 | N7N | Tyr128 [OH] | 0.0 | 1.8 | 0.1 | 6.6 | 0.1 | | 0.0 | 26 | 0.0 | 5.9 | 2.7 |
| | O7N | Tyr126 [OH] | 12 | 20 | 0.0 | 0.1 | 0.0 | | 9.4 | 2.4 | 5.6 | 0.0 | 1.6 |
| | O7N | Tyr128 [OH] | 0.0 | 0.0 | 0.0 | 0.0 | 0.0 | | 0.0 | 8.2 | 0.0 | 1.9 | 2.4 |
| | O7N | Trp105 [NE1]* | 35 | 0.0 | 0.0 | 0.0 | 0.0 | | 31 | 0.0 | 28 | 0.0 | 0.0 |
| 5 | O4D | Tyr128 [OH] | 0.0 | 1.0 | 0.0 | 5.7 | 0.8 | | 1.7 | 25 | 0.1 | 7.1 | 0.1 |
| | O4D | His194 [NE2]* | 0.0 | 0.0 | 0.0 | 0.0 | 0.0 | | 0.0 | 0.0 | 0.0 | 0.0 | 0.0 |
| | O3D | Tyr128 [OH] | 2.8 | 0.0 | 14 | 0.6 | 14 | | 26 | 3.4 | 28 | 1.7 | 0.0 |
| | O2D | His161 [NE2]* | 38 | 0.0 | 0.0 | 0.0 | 0.0 | | 2.6 | 0.0 | 28 | 0.0 | 0.0 |
| 7 | O2A | His194 [NE2]* | 0.0 | 0.0 | 0.1 | 0.1 | 0.0 | | 0.0 | 0.0 | 0.0 | 0.0 | 0.0 |
| | O1A | His194 [NE2]* | 0.1 | 0.1 | 0.0 | 0.1 | 0.0 | | 0.0 | 0.1 | 0.1 | 0.0 | 0.1 |
| | O2B | Asn233 [N] | 0.0 | 0.0 | 0.1 | 0.0 | 0.0 | | 1.9 | 0.0 | 0.0 | 0.1 | 0.0 |
| | O2B | Asn233 [NE2] | 0.1 | 0.0 | 0.2 | 0.0 | 0.0 | | 2.2 | 2.1 | 0.1 | 1.0 | 0.2 |

*#NADH/protein interactions corresponding to the numbering in Figure 6.*
**Most interactions shown for NADHs at active sites 1 and 2 refer respectively to those with chains B and A, with the only exception of those labelled with * that respectively correspond to chains A and B in the hNQO1-NADH homodimer. The cut-off to consider a polar interaction is 3.6 Å.*

**Table S17.** Percentage of polar contact maintained between the FAD and the NADH at each active site of the complex hNQO1-NADH homodimer for each replicate along the 200 ns MD simulation.

| # | NADH atom | FAD atom | Active Site 1 | | | | | Active Site 2 | | | | |
|---|---|---|---|---|---|---|---|---|---|---|---|---|
| | | | R1 | R2 | R3 | R4 | R5 | R1 | R2 | R3 | R4 | R5 |
| | O7N | N1 | 1.7 | 29 | 0.1 | 64 | 0.0 | 4.8 | 11 | 3.7 | 34 | 3.2 |
| | C4N | N5 | 1.2 | 0.1 | 43 | 0.2 | 51 | 2.2 | 5.0 | 7.5 | 1.3 | 1.5 |
| 6 | O2D | O3' | 0.0 | 9.3 | 0.0 | 21 | 0.0 | 0.0 | 17 | 1.4 | 0.4 | 0.1 |

*[#]NADH/protein interactions corresponding to the numbering in Figure 6.*
*The cut-off to consider a polar contact or hydrogen bond interaction is 3.6 Å.*

**Table S18.** Percentage of *pi-stacking* interactions maintained for the NADH at each active site of the complex hNQO1-NADH homodimer for each replicate along the 200 ns MD simulation.

| # | NADH ring | FAD or residue ring | Active Site 1 | | | | | Active Site 2 | | | | |
|---|---|---|---|---|---|---|---|---|---|---|---|---|
| | | | R1 | R2 | R3 | R4 | R5 | R1 | R2 | R3 | R4 | R5 |
| 1 | Nicotinamide | Pyrazine of Isoalloxazine (N5-N10) | 22 | 8.2 | 93 | 8.6 | 89 | 15 | 19 | 33 | 4.0 | 5.7 |
| 2 | Nicotinamide | Pyrimidine of Isoalloxazine (N1-N3) | 58 | 1.7 | 24 | 2.3 | 48 | 37 | 11 | 62 | 0.4 | 54 |
| | Nicotinamide | Tyr126 | 3.3 | 0.0 | 64 | 0.4 | 44 | 8.3 | 18 | 23 | 2.8 | 88 |
| 4 | Nicotinamide | Tyr128 | 1.8 | 6.5 | 4.0 | 0.0 | 71 | 10 | 0.5 | 20 | 0.3 | 1.1 |

*[#]NADH/protein interactions corresponding to the numbering in Figure 6.*
*The cut-off to consider a pi-stacking interaction is 4 Å between the geometric centre of each interacting ring.*
